# Poison frog chemical defenses are influenced by environmental availability and dietary selectivity for ants

**DOI:** 10.1101/2022.06.14.495949

**Authors:** Nora A. Martin, Camilo Rodríguez, Aurora Alvarez-Buylla, Katherine Fiocca, Colin R. Morrison, Adolfo Chamba-Carrillo, Ana B. García-Ruilova, Janet Rentería, Elicio E. Tapia, Luis A. Coloma, David A. Donoso, Lauren A. O’Connell

## Abstract

The ability to use small molecule alkaloids as defensive chemicals, often acquired via trophic interactions, has evolved in many organisms. Animals with diet-derived defenses must balance food choices to maintain their defense reservoirs along with other physiological needs. Poison frogs accumulate skin alkaloids from their arthropod diet, but whether they show preference for specific prey remains unexplored. Here, we explore the role of leaf litter prey availability and diet preferences in shaping poison frog chemical defenses along a geographic gradient. We examined skin alkaloid composition, stomach contents and leaf litter ants in aposematic diablito frogs (*Oophaga sylvatica*) at five sites in northwestern Ecuador, and in sympatric, cryptic Chimbo rocket frogs (*Hyloxalus infraguttatus*) at one site. We found that differential availability of leaf litter ants influenced alkaloid profiles across diablito populations, and low levels of alkaloids were observed in the sympatric, ‘undefended’ Chimbo rocket frog. Ants were the primary dietary component of the defended species, while ‘undefended’ species ate other prey categories including beetles and larvae in addition to ants. A prey selection analysis suggested that defended and ‘undefended’ frogs both feed on a high proportion of specific small ant genera that naturally contain alkaloids, suggesting that selectivity for toxic prey is not restricted to classically aposematic and highly toxic species. These findings suggest that poison frogs’ use of feeding resources relative to availability may be an understudied and important selection factor in the evolution of acquired defenses.

**Resumen:** La capacidad de usar alcaloides como productos químicos defensivos ha evolucionado en muchos organismos, a menudo a través de interacciones tróficas. Los animales con defensas que se derivan de la dieta deben balancear la selección de los alimentos para mantener sus reservas defensivas junto con otras necesidades fisiológicas. Las ranas venenosas acumulan alcaloides en la piel procedentes de su dieta de artrópodos, pero aún no se ha explorado si muestran preferencia por presas específicas. En este estudio, investigamos el papel de la disponibilidad de presas en la hojarasca y las preferencias dietarias en la configuración de las defensas químicas de las ranas venenosas a lo largo de un gradiente geográfico. Examinamos la composición de alcaloides en la piel, el contenido estomacal y la presencia de hormigas de la hojarasca en ranas aposemáticas diablito (Oophaga sylvatica) en cinco sitios del noroeste de Ecuador, así como en la especie críptica y simpátrica rana cohete de Chimbo (Hyloxalus infraguttatus) en un sitio. Encontramos que la disponibilidad diferencial de hormigas en la hojarasca influyó en los perfiles de alcaloides en las poblaciones de diablito, y se detectaron niveles bajos de alcaloides en la rana cohete de Chimbo, previamente considerada ‘indefensa’. Las hormigas fueron el componente principal de la dieta en la especie defendida, mientras que la especie ‘indefensa’ consumió otras categorías de presas, incluidos escarabajos y larvas, además de hormigas. Un análisis de selección de presas indicó que tanto las ranas defendidas como las indefensas prefieren ciertos géneros de hormigas pequeñas que contienen alcaloides de forma natural, lo que sugiere que la preferencia por presas tóxicas no está restringida a especies clásicamente aposemáticas y altamente tóxicas. Estos hallazgos sugieren que las preferencias de comportamiento pueden ser un factor de selección poco estudiado, pero crucial, en la evolución de las defensas adquiridas.

## Introduction

Many organisms use chemical defenses to protect themselves from predators or pathogens (Mebs 2002). These defenses often involve small molecule alkaloids synthesized by plants or microbes, and some taxa can acquire them through dietary sequestration (Roberts and Wink 1998; Agrawal et al. 2012; Santos et al. 2016). Phytophagous insects represent the most well-studied taxa, including some species that specialize in particular plant species and accumulate specific secondary metabolites for chemical communication or defense (Roberts and Wink 1998; Walsh and Tang 2017; Beran and Petschenka 2022). Although our understanding of chemical defense in vertebrates is more sparse than invertebrates, poison frogs are a well known example of chemical defenses acquired through an arthropod-based diet (Savitzky et al. 2012). A key component of poison frog diets includes alkaloid-rich arthropods, and these frogs have evolved the physiological mechanisms to tolerate and integrate the toxin in their tissues to deter predators (Alvarez-Buylla et al., 2023). Yet, it remains unclear whether the ecological principles underlying arthropod diet specialization on toxic plants also apply to vertebrates that sequester their chemical defenses from arthropods.

The chemical repertoire of alkaloid-defended species varies within and between populations. For example, the composition and concentration of piperidine alkaloids in the Norwegian spruce (*Picea abies*) differ by location (Virjamo and Julkunen-Tiitto 2016), while in fire ants (*Solenopsis* spp.) vary within species (Deslippe and Guo 2000). In species with acquired chemical defenses, variation in alkaloid profiles is generally attributed to spatio-temporal shifts in the availability of alkaloid-containing food, often resulting from environmental variation in temperature, rainfall, and other climatic factors. For instance, in Argentine *Melanophryniscus* toads (Daly et al. 2007), Malagasy *Mantella laevigata* frogs (Moskowitz et al. 2018), and various species of Neotropical poison frogs (Family Dendrobatidae; Saporito et al. 2006; Saporito, et al. 2007; Prates et al. 2019; Moskowitz et al. 2020), differences in the composition of skin alkaloids across localities and seasons correspond with differences in stomach contents and leaf litter arthropod communities. Dietary selectivity may also influence the alkaloid profile of chemically defended species by favoring the consumption of food items that contain specific defensive alkaloids. Yet, organisms must make dietary decisions based on handling time and nutritional value, in addition to food availability and maintenance of chemical defenses. For instance, *Chiasmocleis leucosticta* frogs preferred smaller ants over larger, more aggressive genera (Meuer et al. 2021), while lab-reared non-toxic *Dendrobates tinctorius* preferred protein-rich larvae over other prey types including ants (Moskowitz et al. 2022). As these studies suggest, variation in the environment, the availability of alkaloid-containing prey, prey phenotype and foraging behavior are likely important factors in diet-acquired defense evolution. However, there is limited understanding of how these factors interact to influence species’ food choices and their ability to sequester chemical defenses from specific dietary sources.

Neotropical poison frogs acquire alkaloids from alkaloid-containing arthropods rather than synthesizing them de novo (Daly et al. 1994). Chemical defenses in dendrobatids have evolved independently at least four times in parallel with dietary specialization on ants and mites (Santos et al. 2003; Darst et al. 2005). However, recent evidence suggests that diet specialization alone does not explain the defended phenotype, as ‘undefended’ species often consume alkaloid-containing arthropods and have low but detectable alkaloid levels (Sanches et al. 2023; Tarvin et al. 2024). Yet, it is unclear if selectivity for specific alkaloid-containing prey plays a role in the ability of poison frogs to dietarily acquire their chemical defenses, as most studies focus on diet without assessing environmental availability of arthropod prey (McElroy and Donoso 2019). This is especially important as temporal and geographic variations in abiotic factors such as temperature, altitude, and precipitation affect the composition and richness of leaf litter arthropod communities (Brühl et al. 1999; Silva and Brandão 2014; Gibb et al. 2015; Tiede et al. 2017; Hoenle et al. 2022; Basset et al. 2023). Thus, we currently lack a framework for understanding the evolution of diet-acquired defenses in poison frogs, as the role of prey availability in diet specialization has not been studied in depth.

Here, we tested whether dendrobatid poison frogs that acquire chemical defenses from their diet exhibit dietary prey selectivity. To test this hypothesis, we sampled stomach contents, skin alkaloids, and surrounding leaf litter ant communities from five populations of the aposematic, chemically defended *Oophaga sylvatica* and one sympatric population of the cryptic, chemically ‘undefended’ *Hyloxalus infraguttatus.* We compared skin alkaloid profiles among poison frog populations along a geographical gradient, predicting intraspecific variation linked to environmental factors, such as altitude, temperature and precipitation, and higher alkaloid composition in the sympatric defended species. We compared stomach contents across poison frog populations and predicted within and between species variation, where chemically defended frogs would eat more ants and mites than ‘undefended’ frog species. We sorted ants by genus from stomachs and leaf litter, characterized their morphology, and predicted that sympatric defended and ‘undefended’ frogs will show distinct dietary selectivity for ant genera despite access to the same ant communities. We further predicted that differences in ant communities across localities correspond with differences in chemical defense between poison frogs. Together, our between- and within-species comparisons of poison frog stomach contents and prey availability aim to disentangle the relationship between dendrobatid diet and alkaloid acquisition, which has broader implications for our general understanding of trophic interactions and the evolution of chemical defenses across taxa.

## Methods

### Study system and sample collection

Diablito frogs (*Oophaga sylvatica*) were collected in May 2019, during daylight in secondary forests near the towns of Ceiba (N=10; 207 m.a.s.l), Cristóbal Colón (N=11; 221 m.a.s.l), Puerto Quito (N=11; 302 m.a.s.l), Santo Domingo de los Tsáchilas (N=10; 632 m.a.s.l), and La Maná (N=20; 480 m.a.s.l), in Northwestern Ecuador (Figure 1A). As *O. sylvatica* suffers from illegal poaching for the pet trade, coordinates for collection can be obtained from the corresponding authors. Chimbo rocket frogs (*Hyloxalus infraguttatus*) were collected during daylight from La Maná (N=9). While behavioral observations were not performed, frogs were collected during active foraging hours (06:00 - 19:00; Funkhouser 1956; AmphibiaWeb 2025). Frogs were anesthetized 3 – 6 hours after collection with 20% benzocaine gel applied to the ventral skin and euthanized. For each individual, the dorsal skin was dissected and stored in methanol in glass vials. The stomach contents were stored in 100% ethanol in 1.5 ml plastic tubes. Remaining frog tissues were either preserved in 100% ethanol or RNAlater (Thermo Scientific, Waltham, MA, USA), or deposited in the amphibian collection of Centro Jambatu de Investigación y Conservación de Anfibios in Quito, Ecuador. Collections and exportation of specimens were done under permits (Collection permit: No. 0013-18 IC-FAU-DNB/MA; Export permit: No. 214-2019-EXP-CM-FAU-DNB/MA; CITES export permit No. 19EC000036/VS) issued by the Ministerio de Ambiente de Ecuador. The Administrative Panel on Laboratory Animal Care of Stanford University approved all frog-related procedures (Protocol #34153).

**Figure 1.**
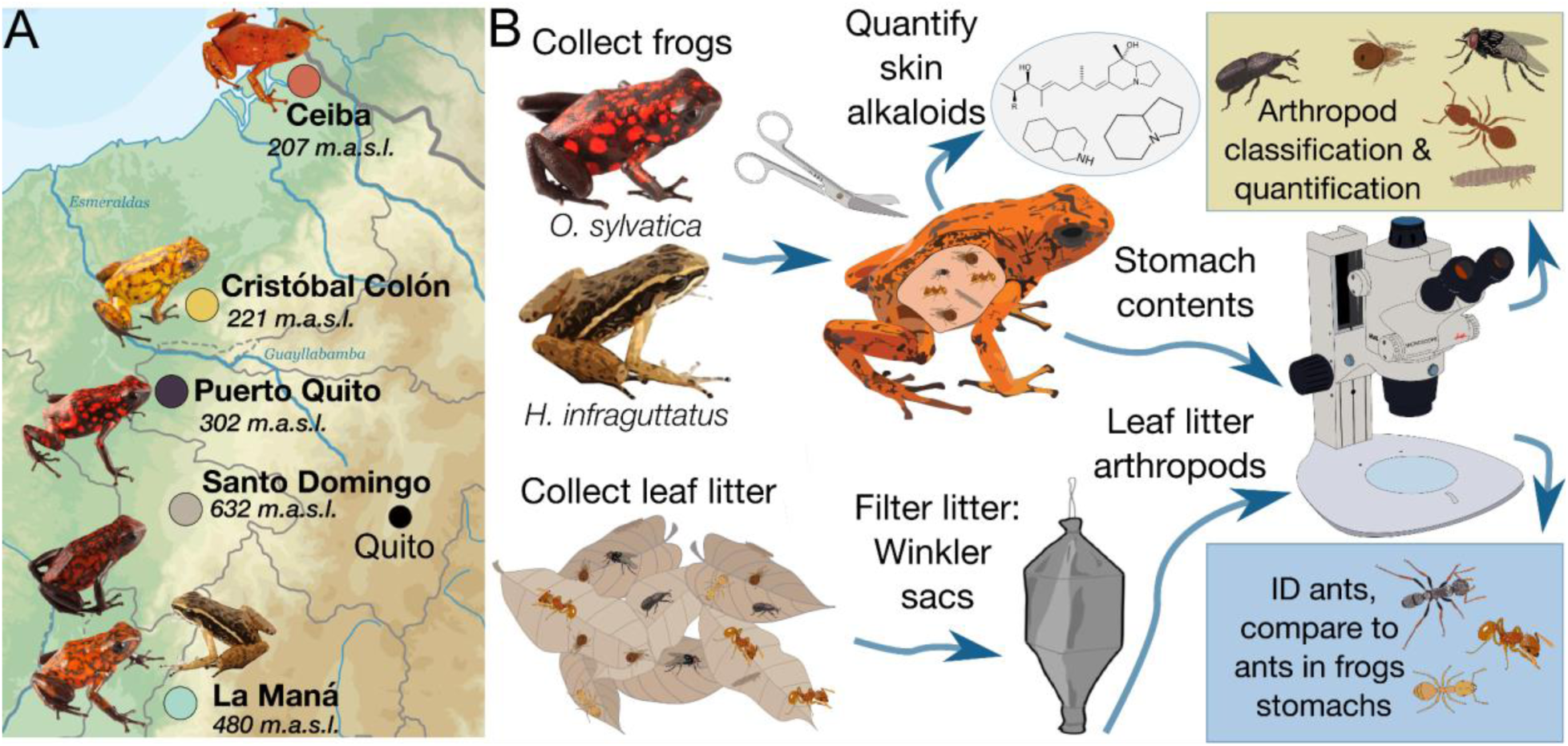
Map of frog populations and experimental workflow. **(A)** Collection sites are shown along a geographical gradient on a topographic map of western Ecuador. Note that both *Hyloxalus infraguttatus* and *Oophaga sylvatica* were collected from La Maná. ma.s.l. = meters above sea level. **(B)** A flowchart depicts the main steps of data collection for frog stomach content and leaf litter samples, in order to compare ant genera abundances between these groups.

### Alkaloid extraction and quantification

Skins were removed from methanol with forceps and weighed. From the methanol in which the skin was stored, 1 ml was syringe filtered through a 0.45 um PTFE filter (Thermo Scientific, 44504-NP) into the new glass vial (Wheaton, PTFE caps, 60940A-2) supplemented with 25 ug (-)-nicotine (Sigma Aldrich, N3876-100ML). All tubes were then capped, vortexed, and stored for 24 hours at -80°C to precipitate lipids and proteins. After precipitating for 24 hours, the supernatant was filtered through a 0.45 um PTFE syringe filter into a new glass vial. A 100 uL aliquot was added to a gas chromatography/mass spectrometry (GC/MS) autosampler vial, and the remaining solution was stored at -80°C.

Alkaloid detection was performed using gas chromatography-mass spectrometry (GC-MS) following the protocols described elsewhere (Saporito et al. 2010; Alvarez-Buylla et al. 2024), and using a Shimadzu GCMS-QP2020 instrument with a Shimadzu 30m x 0.25 mmID SH-Rxi-5Sil MS column. In brief, the separation of alkaloids was achieved with helium as the carrier gas (flow rate: 1 mL/min) using a temperature program increasing from 100 to 280°C at a rate of 10°C/minute. This was followed by a 2-minute hold and an additional ramp to 320°C at a rate of 10°C/minute for column protection reasons, and no alkaloids appeared during this part of the method. Compounds were analyzed with electron impact-mass spectrometry (EI-MS). The GC-MS data files were exported as .CDF files, and the Global Natural Products Social Molecular Networking (GNPS) software was used to perform the deconvolution and library searching against the AMDIS (NIST) database to identify all compounds (Wang et al. 2016; Aksenov et al. 2021). For deconvolution (identification of peaks and abundance estimates), the default parameters were used. Through the deconvolution process, molecular features were reported as rows/observations, while m/z intensities were reported as columns/variables. Automatic library search was obtained from reference libraries of natural products (NIST, Wiley, University of CORSICA, GNPS), and our resulting dataset was filtered to keep only the nicotine standard and alkaloids previously found in poison frogs or compounds with the same base ring structure and R group positions as those classes defined in the Daly poison frog alkaloid database (Daly et al. 2005). Once the feature table from the GNPS deconvolution was filtered to include only poison frog alkaloids and nicotine, the abundance values (ion counts) were normalized by dividing by the nicotine standard and skin weight. The resulting filtered and normalized feature table was used for all further analyses and visualizations.

### Frog stomach contents identification

Whole stomachs were stored in 100% ethanol at -20°C until processing. Stomach contents were sorted and photographed. Prey items in the photographs were identified to the lowest possible taxonomic rank and then grouped into broad diet categories: ants, mites, larvae, beetles, and “other” (i.e., any non-larval arthropods not described by the other four groups). Note that most larvae belong to Dipteran or Coleopteran taxa, but are considered in our distinct “larvae” category given their differences in appearance from adults. The vast majority of identifiable prey remained whole, with the exception of ants, whose heads frequently detached from their bodies upon ingestion. To prevent overcounting, only whole ant specimens, partial ant specimens with heads, or individual ant heads were counted. Ant specimens were identified to genus using a reference collection of Ecuadorian ants (Donoso and Ramón 2009; Salazar et al. 2015; Donoso 2017).

### Leaf litter communities and ant morphology

Leaf litter samples were extracted and collected using 18 - 20 Winkler sacs from all localities except for *La Ceiba* due to time and resource limitations. Samples were collected in May–June 2019 within one square meter of where a frog had been previously collected that day. Leaf litter arthropods were extracted from 1 m^2^ and hung within Winkler sacs for 24 hours, during which the arthropods were collected into 70% ethanol. Only ants were identified, as described above. Collection of ant specimens was done under permits issued by the Ministerio de Ambiente de Ecuador to Museo de Historia Natural Gustavo Orcés at Escuela Politécnica Nacional (MAE-DNB-CM-2017-0068). Ant morphology was characterized by 17 traits related to size, texture, spine count, and coloration for leaf litter ant genera that were also found in frog stomach contents. Morphological traits were measured at the species level using data from the Global Ants Database, which provides standardized trait information across ant species (Parr et al. 2017).

### Data Analysis

Statistical analyses and figures were generated using R Studio (version 2025.05).

#### Alkaloid comparisons

We used a Kruskal-Wallis test to examine overall differences between *O. sylvatica* and *Hyloxalus infraguttatus* populations in summed toxicity across the 79 alkaloids in the 13 structural families. A pairwise Wilcoxon test was used to determine *O. sylvatica* population differences between the 13 structural families in both species, with p-values adjusted for multiple testing using the False Discovery Rate control (FDR).

#### Frogs’ diet comparisons

We visualized interactions between arthropod prey taxa and poison frog species in a bipartite network based on the average abundance of arthropod prey for every frog population. We used the species specificity index (ssi), as implemented in the specieslevel() function of the ‘bipartite’ package, to quantify the variability in dietary interactions between arthropod prey taxa and poison frog species in a bipartite network. This index reflects the degree to which each frog species interacts unevenly across prey taxa, with values ranging from 0 (indicating low variability and generalist behavior) to 1 (indicating high variability and specialist behavior). Additionally, we used generalized linear mixed models (GLMM) to test for compositional differences of frog diet categories using the function glmmTMB() within the package ‘glmmTMB’ (Brooks et al. 2017). We used a negative binomial distribution appropriate for count data with overdispersion. We tested for diet differences by including species/population, prey type, and their interaction as main effects. Frog individual tags were included as a random variable to account for repeated sampling of prey categories within individuals. We computed estimated marginal means to test for pairwise comparisons between populations using the ‘emmeans’ package (Lenth 2023).

#### Ant-vs mite-derived alkaloids

To infer potential dietary origins of frogs’ skin alkaloids, we cross-referenced the structural classes of alkaloids with the determined arthropod source reported by Santos et al. (2016), which assigns compounds as derived from ants, mites, or both. We tested whether the proportion of ant-based alkaloids is greater than the proportion of mite-based alkaloids across frog populations using an anova followed by a Tukey post-hoc test for multiple comparisons. We visualized the proportion of ant- and mite-derived alkaloids using a chord diagram. Additionally, we tested whether total alkaloid abundance, as well as abundance within structural families, correlated with the number of ants and mites consumed across populations.

#### Skin alkaloids vs. leaf litter ant communities along a geographical gradient

We analyzed compositional differences in skin alkaloid profiles of *O. sylvatica* populations and *H. infraguttatus*, and their surrounding leaf litter ant communities in two separate non-metric multidimensional scaling (NMDS), using the function metaMDS() within the package ’vegan’ (Oksanen et al. 2022). Statistical differences between and within populations were assessed using a permutational multivariate analysis of variance (PERMANOVA) on Bray-Curtis dissimilarities. P-values for pairwise comparisons were adjusted using the function pairwise.adonis(). Additionally, we used the envfit() function to understand the influence of altitude, ambient temperature and precipitation on both alkaloid composition and leaf litter ant communities. These environmental variables were selected to capture spatial and temporal variation in ecological conditions that could influence ant availability and frogs’ foraging behavior. Ambient temperature and precipitation data was extracted from WorldClim 2.1, at a spatial resolution of 30 arc-seconds (∼1 km²) for each study site (Fick and Hijmans 2017). Similarly, we used envfit() to explore the contribution of specific alkaloid classes and ant genus to the ordination spaces. We used Procrustes analysis to test similarities between the NMDS ordinations of skin alkaloids and ant community composition across sites. The Procrustes correlation was calculated using the procrustes() function from the ‘vegan’ package, which aligns the two ordinations by scaling, rotating, and translating one configuration to best match the other. Statistical significance of the correlation was evaluated with 999 permutations using the protest() function.

#### Stomach vs. leaf litter ant communities and frogs’ preference for ant genera

To test for within and between species differences in ant abundance between leaf litter and stomach contents, we employed a negative binomial generalized linear model using the glm.nb() function from the ‘MASS’ package (Venables and Ripley 2002). The model used total ant abundance per sample as the response variable, with group (stomach vs. leaf) and population as predictors. Pairwise comparisons were conducted using estimated marginal means. To examine if frogs show selectivity for consuming specific ant genera, we calculated a linear selectivity index by subtracting the relative abundance of ants found on the leaf litter from those found in frog stomach contents (Strauss, 1979). The interpretation of these selectivity values was guided by the methodology in McElroy and Donoso (2019) (McElroy & Donoso, 2019), which involves generating a null distribution of selectivity values via simulation for each ant species. By comparing the observed selectivity values to this null distribution, we classified the ants as ‘selected’ if the values were above, ‘neutral’ if they were within, and ‘avoided’ if they were below the null distribution, thereby delivering a statistically robust assessment of the frogs’ selective foraging behaviors.

#### Ant morphology vs frog selectivity

To determine whether the morphological and life history traits of ant species in the leaf litter influences frog selectivity, we performed a Principal Component Analysis (PCA). The PCA aimed at reducing the dimensionality of 17 morphological and life history traits across 17 ant genera found both in the leaf litter and in *O. sylvatica* stomachs. To build the PCA, we used the function dudi.pca() from the ‘ade4’ package (Dray & Dufour, 2007). Scores of components with eigenvalues higher than 1 were selected and used as response variables in a pairwise Wilcoxon test to determine differences between selectivity categories for ants in all diablito populations. P-values were adjusted for multiple testing using the False Discovery Rate control (FDR).

## Results

### Alkaloids differ between species and across diablito frog populations

Skin extracts consisted of 79 alkaloids. The summed amount of alkaloids varied across species and populations (Kruskal-Wallis; X^2^(5) = 41.542, p < 0.001; Figure 2A), with *O. sylvatica* having more alkaloids than *H. infraguttatus* (*H. infraguttatus* vs. all other *O. sylvatica* populations, p < 0.001; Table S1). Within *O. sylvatica*, the Ceiba population had less toxins than all others (p < 0.001; Table S1), while frogs from Santo Domingo had on average the highest alkaloid load (Table S1).

**Figure 2:**
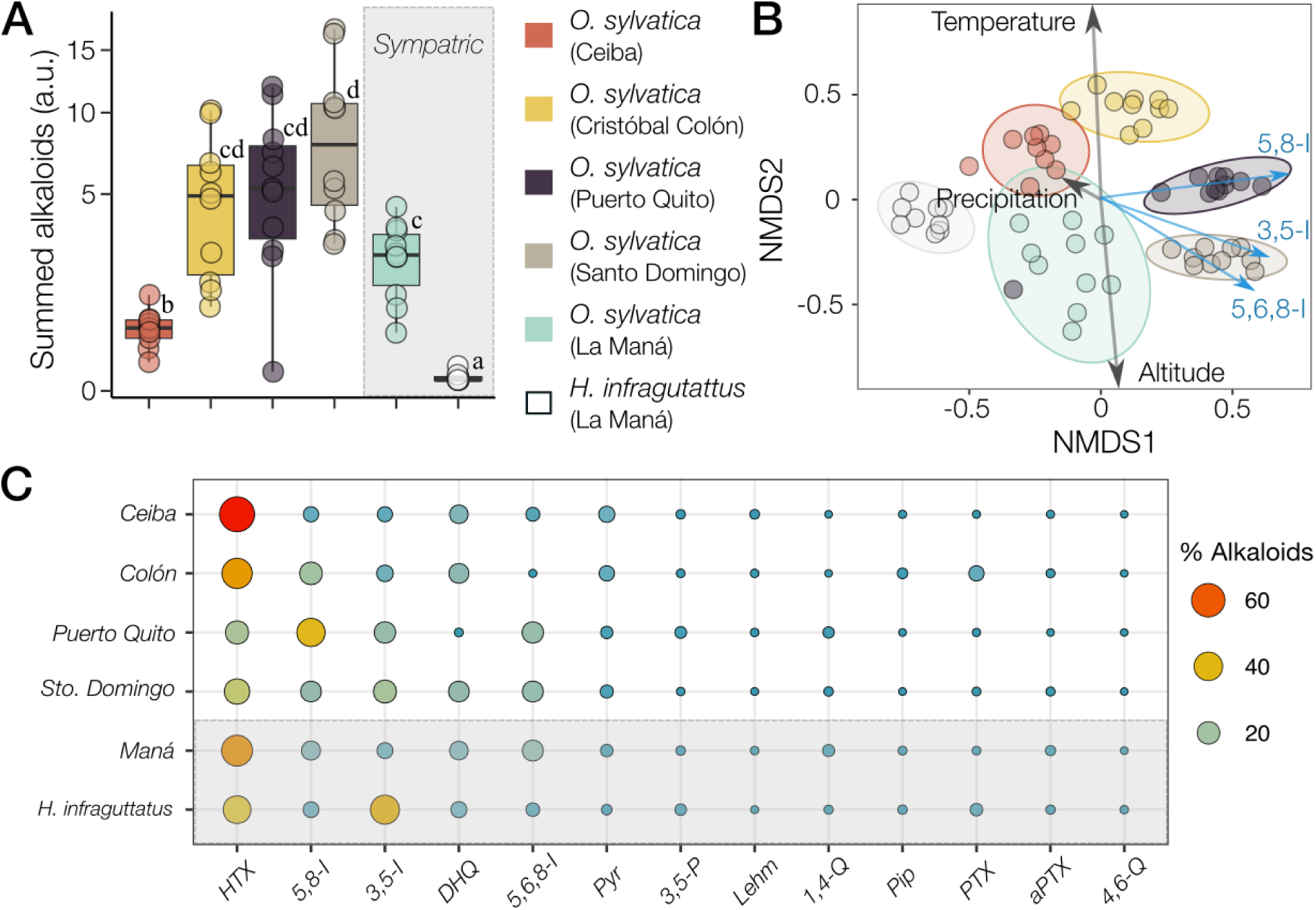
Alkaloid profiles differ between species and populations. **(A)** The summed relative abundances of 79 alkaloids are visualized as box plots, where each dot represents a frog. The y axis is square-root transformed for visual clarity and summed alkaloids are expressed in arbitrary units (a.u.). Populations are represented with different colors. *Hyloxalus infraguttatus*, collected near the town of La Maná, is shown on the far right. Groups not connected by the same letter are significantly different (Table S1). **(B)** Non-metric multidimensional scaling (NMDS) biplot based on Bray-Curtis dissimilarities show alkaloid profile differences between populations based on the relative abundances of 79 unique alkaloids (Stress = 0.16). Each dot represents a single frog’s alkaloid profile. Ellipses represent 95% confidence intervals. Environmental factors (black arrows) and alkaloids (blue arrows) in correlation with the ordination are shown. **(C)** Bubble - heatmap shows percent of all summed alkaloids grouped by alkaloid structural class and population. Histrionicotoxins (HTX); 5,8-disubstituted indolizidines (5,8-I); 3,5-disubstituted indolizidine (3,5-I); Decahydroquinoline (DHQ); 5,6,8-trisubstituted indolizidines (5,6,8-I); Pyrrolidine (Pyr); 3,5-disubstituted pyrrolizidine (3,5-P); Lehmizidine (Lehm); 1,4-disubstituted quinolizidine (1,4-Q); Piperidine (Pip); Pumiliotoxin (PTX); Allopumilliotoxin (aPTX); 4,6-disubstituted quinolizidine (4,6-Q).

We next visualized overall alkaloid compositional differences across *O. sylvatica* populations and *H. infraguttatus* using an NMDS (Figure 2B). The NMDS suggested a two dimensional solution (stress = 0.156) and showed distinct clusters of alkaloid composition. The abundance of all 79 alkaloids varied significantly across groups (PERMANOVA, F_(4)_ = 10.178, p < 0.001). A post-hoc pairwise comparison indicated significant differences between all possible population pairs (Table S2; Figure 2B), suggesting each group has a unique alkaloid profile. Fitting environmental variables into the NMDS indicated that altitude (r² = 0.72, p = 0.001) and temperature (r² = 0.68, p = 0.001) significantly influenced alkaloid composition across the geographical gradient (Figure 2B). Given their strong inverse correlation and closely aligned vectors in NMDS space, we report both as reflecting a shared environmental gradient.

Skin alkaloids fell into one of 13 structural families: Histrionicotoxins (HTX); 5,8-disubstituted indolizidines (5,8-I); 3,5-disubstituted indolizidine (3,5-I); Decahydroquinoline (DHQ); 5,6,8-trisubstituted indolizidines (5,6,8-I); Pyrrolidine (Pyr); 3,5-disubstituted pyrrolizidine (3,5-P); Lehmizidine (Lehm); 1,4-disubstituted quinolizidine (1,4-Q); Piperidine (Pip); Pumiliotoxin (PTX); Allopumilliotoxin (aPTX); 4,6-disubstituted quinolizidine (4,6-Q). When examined more closely, histrionicotoxins, decahydroquinolines and indolizidines were the most abundant alkaloid classes relative to the total alkaloid content in both *H. infraguttatus* and all *O. sylvatica* populations sampled (Figure 2C). Several indolizidines, including 5,6,8-I (r² = 0.54, p = 0.001), 5,8-I (r² = 0.54, p = 0.001) & 3,5-I (r² = 0.46, p = 0.001) contributed significantly to alkaloid ordination (Figure 2B).

### Defended frogs consumed more ants relative to other prey types and compared to the diet of the undefended species

We found that the number of prey consumed in different categories differs significantly across populations and between species (GLMM, population x prey type: X^2^(20) = 83.59, p < 0.001; Figure 3A & B). Post hoc pairwise comparisons and the species selectivity index (*ssi*) showed that all *O. sylvatica* populations consumed significantly more ants than other prey categories (x̅ = 75%; emmeans _(ants vs. all prey)_: p-value = <0.001; *ssi_range_*= 0.66 - 0.85; Figure 3B, Table S3), whereas *H. infraguttatus* showed a more generalist dietary pattern, consuming a smaller but diverse array of arthropods including ants (45%), beetles (14%) and ‘other’ arthropods (25%; emmeans _(all prey comparisons)_: p-value = >0.05; *ssi* = 0.27; Figure 3B, Table S3). It is worth noting that only one *H. infraguttatus* individual had 18 ants in the stomach, which accounts for nearly half of the total consumed for this species in our dataset. When removing this individual, ants made up 36% of the total diet, followed by ‘other’ arthropods (29.6%) and beetles (16.5%).

**Figure 3:**
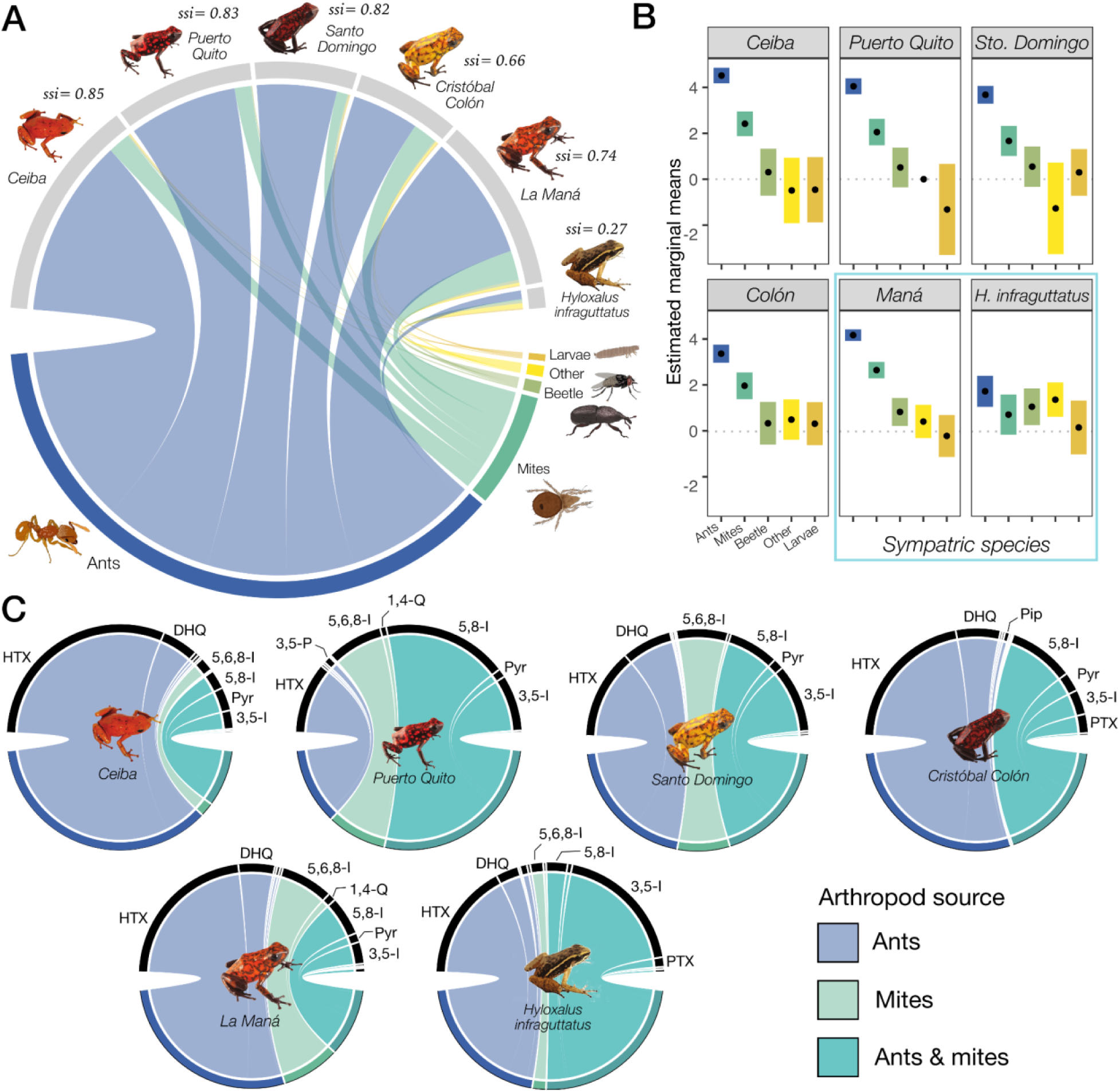
Diet differs between frog species and across diablito frog populations. **(A)** Chord diagram showing a bipartite network of the dietary interactions between two frog species, the non-toxic *Hyloxalus infraguttatus* and the toxic *O. sylvatica*, and their arthropod prey in five localities. The thickness of the connecting bars represents the average number of prey items consumed, indicating its relative importance in the network. Species specificity index (*ssi*) is shown next to each frog species, ranging from 0 (generalist) to 1 (specialist). **(B)** Pairwise comparison of prey item consumption between frog species. Dots represent the mean and bars represent 95% confidence intervals for the estimated marginal means. Non-overlapping bars indicate statistically significant differences. **(C)** Chord diagrams showing the most abundant alkaloid families and their putative arthropod source, across frog populations (*sensu* Santos et al. 2016). The thickness of the connecting bars represents the summed alkaloid abundance per structural class. Histrionicotoxins (HTX); 5,8-disubstituted indolizidines (5,8-I); 3,5-disubstituted indolizidine (3,5-I); Decahydroquinoline (DHQ); 5,6,8-trisubstituted indolizidines (5,6,8-I); Pyrrolidine (Pyr); 3,5-disubstituted pyrrolizidine (3,5-P); Lehmizidine (Lehm); 1,4-disubstituted quinolizidine (1,4-Q); Piperidine (Pip); Pumiliotoxin (PTX); Allopumilliotoxin (aPTX); 4,6-disubstituted quinolizidine (4,6-Q).

### Frogs’ alkaloid composition suggests a predominantly ant-based diet

We found that across all frog populations, including the “non-toxic” species *H. infraguttatus*, ant-derived alkaloids appeared to represent the most abundant component of their chemical profile, while a smaller but important contribution was attributed to mite-derived alkaloids (Figure 3C). Tukey’s post hoc test revealed that the proportion of ant-derived alkaloids was significantly higher than mite-derived alkaloids (p<0.001). Neither total alkaloid abundance nor abundance within structural families showed a significant correlation with the number of ants or mites consumed across populations (All p-values > 0.05; TABLE S4).

### Alkaloid diversity among sites is associated with variation in leaf litter ant communities

As the diet of *O. sylvatica* is mainly composed of ants, we looked at compositional differences in leaf litter ant communities across the geographical gradient. A total of 46 ant genera were recovered from Winkler traps in leaf litter communities. The NMDS suggested a two-dimensional solution (stress = 0.130), representing partially overlapping clusters of antcommunities across localities in the ordination space (Figure 4A). PERMANOVA and post-hoc comparisons suggested significant differences in the composition of ant communities between all pairs of localities (PERMANOVA, F(3) = 5.0525, p < 0.001; Table S5, Figure 4A). Precipitation (r² = 0.41, p = 0.001), altitude (r² = 0.48, p = 0.001) and temperature (r² = 0.49, p = 0.001) significantly influenced ant availability across the geographical gradient (Figure 4B). A subset of ant genus, including *Octostruma* (r² = 0.59, p = 0.001), *Pachycondyla* (r² = 0.54, p = 0.001), *Rogeria* (r² = 0.51, p = 0.001) and *Solenopsis* (r² = 0.48, p = 0.001) significantly contributed to the ordination. We next asked whether availability of leaf litter ant communities between locations is associated with skin alkaloid diversity. The procrustes analysis suggested that variation in alkaloid diversity among sites is significantly correlated with variation in availability of ant communities (r = 0.47, p-value = 0.001, 999 permutations; Figure 4B).

**Figure 4:**
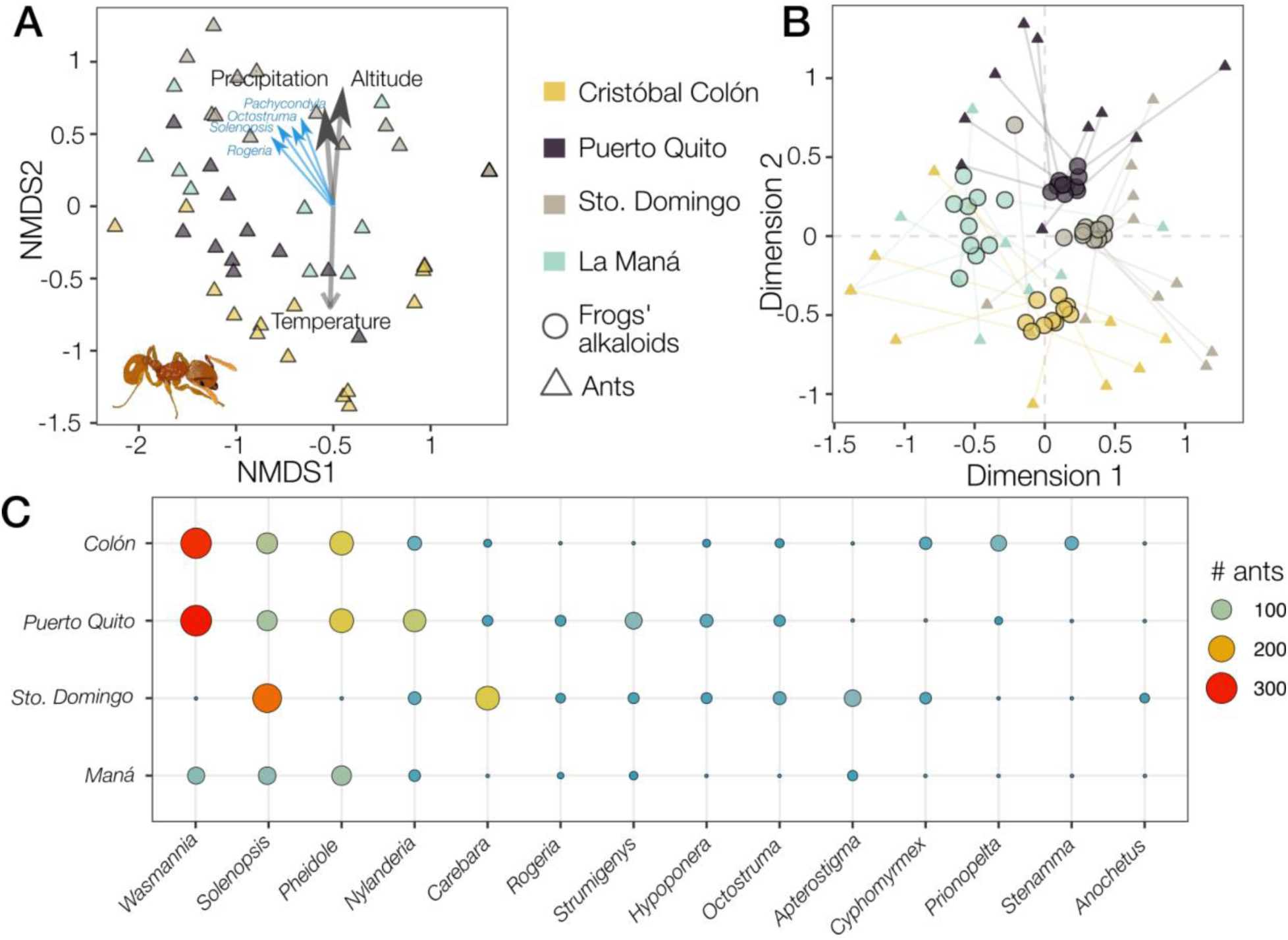
Ant community composition varies across localities. **(A)** Non-metric multidimensional scaling (NMDS) biplot based on Bray-Curtis dissimilarities showing ant communities across locations based on the relative abundances of 113 ant species (Stress = 0.156). Each triangle represents ant communities at one sampling point. Environmental factors (black arrows) and ant genera (blue arrows) in correlation with the ordination are shown. **(B)** Procrustes ordination plot showing correspondence between frog alkaloids and ant communities ordinations across localities. Procrustes residuals (lines) connect paired points between frog alkaloid profiles (circles) and ant community composition (triangles) at each site. Shorter lines indicate better alignment between the ordinations, suggesting stronger correspondence. **(C)** Bubble - heatmap shows the number of ants, grouped by genus (with ≥10 individuals), across frog populations. Each ant community was sampled using a trap placed next to the corresponding frog’s capture site.

### Frogs show different dietary selectivity for particular ant genera

We next asked if frogs eat specific ant genera selectively or if their ant diet reflects the genera of the surrounding leaf litter communities. From the 46 ant genera recovered from Winkler traps in leaf litter communities, only 17 of these were consumed by frogs across different species and populations. Our results indicated no significant differences in the total abundance of ants between frog stomach contents and leaf litter across all populations, except for Santo Domingo and *H. infraguttatus* where frogs had lower abundance of ants in their stomachs compared to the leaf litter (Table S6; Figure 5A). We found that *Solenopsis* was the most selected ant genus across all *O. sylvatica* populations, while *Pheidole* was the only selected ant genus by *H. infraguttatus*. Particularly, frogs from the Cristobal Colón population showed selectivity for *Paratrachymyrmex, Crematogaster and Pheidole*, whereas *Strumigenys* and *Cyphomyrmex* were selected in La Maná and Puerto Quito populations, respectively (Figure 5B).

**Figure 5:**
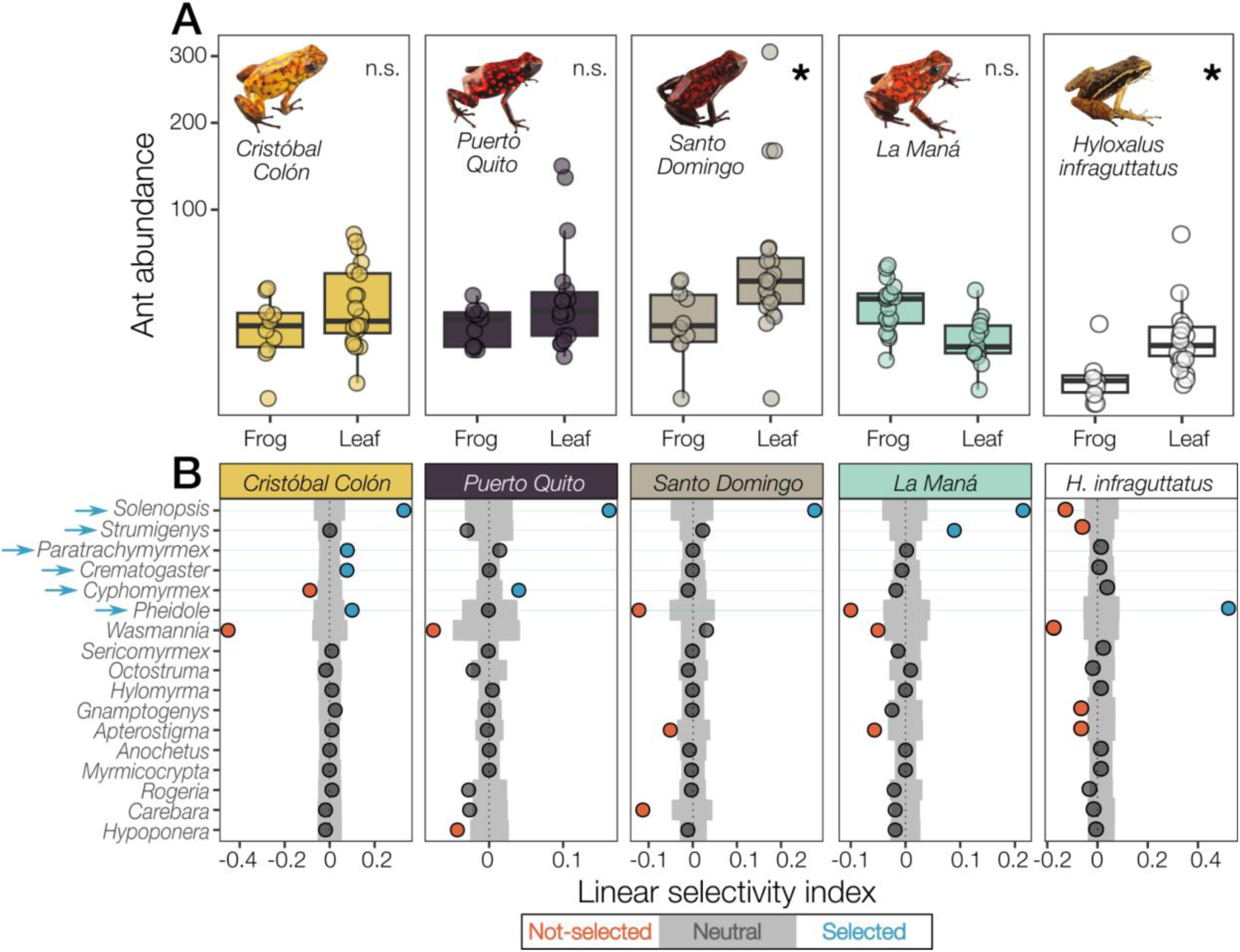
Relative abundance and selectivity for ant genera differs across species and within diablito localities. **(A)** Boxplot showing differences across populations in total abundance of ants within 17 ant genera found in both leaf litter and frog stomach samples. The y axis is square-root transformed for visual clarity. n.s. = non-significant. * p-value<0.05 **(B)** Linear selectivity index for 17 ant genera eaten by the toxic *Oophaga sylvatica* populations and the non-toxic *Hyloxalus infraguttatus*. Grey bars denote simulated null distribution. Points denote categorical selectivity as follows: ‘not-selected’ if they are below the null distribution (red dots), ‘neutral’ if they are within (black dots), and ‘selected’ if the values are above (blue dots). Blue arrows indicate overall selected ant genera.

On the other hand, the remaining ants were either not selected or occasionally consumed (Figure 5). For example, *Wasmannia* ants were avoided by all populations, except for Santo Domingo frogs where it was consumed in proportion to its availability (i.e., neutral). Similarly, *Apterostigma* ants were avoided by frogs in Santo Domingo and in the sympatric populations of *O. sylvatica* and *H. infraguttatus* in La Maná.

### Ant morphology influences prey selectivity in frogs

Finally, we asked whether *O. sylvatica* frogs select for specific ant traits. A principal components analysis on 17 morphological and life history ant traits yielded three principal components, each with eigenvalues exceeding 1. After inverting the scores of these components to aid interpretability, we found that the first component accounted for 55.66% of the total variance, primarily reflecting size and texture traits, with higher scores indicating larger ants. Component number two explained 11.02% of the variance and was strongly associated with color traits, where higher scores corresponded to ants with dark-brown body coloration. The third principal component, explaining 7.15% of the variance, was largely indicative of the presence of spines, with higher scores representing spiny ants (Figure 6). A pairwise Wilcoxon test on these principal components revealed that ants categorized as selected were significantly smaller than those with neutral selectivity (p-value = 0.03; Figure 6A; Table S7), but not than those not selected (p-value = 0.162; Figure 6A). Contrary, no differences were found in the traits of color and the presence of spines across the categories of selectivity (all p values >0.05; Figure 6B & C; Table S7).

**Figure 6:**
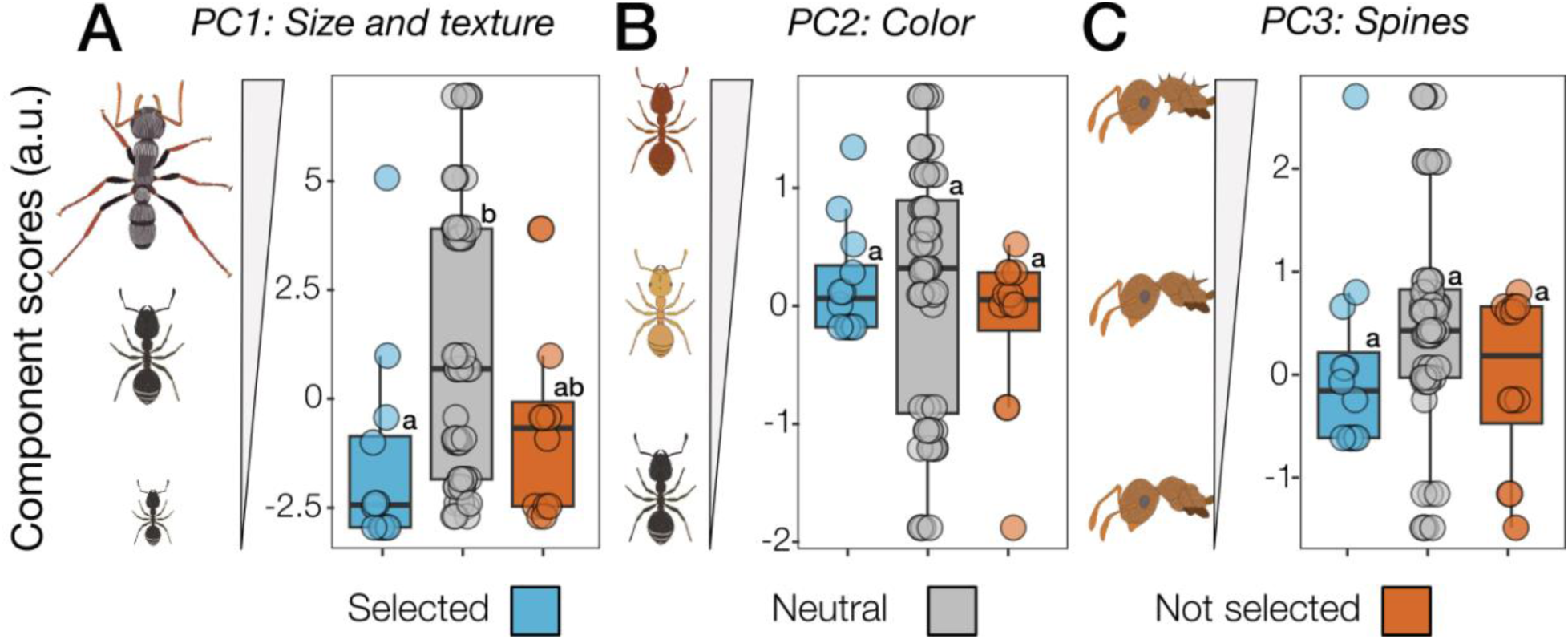
Boxplots showing differences between categorical selectivity in principal component scores on 17 ant traits. **(A)** First principal component (PC1) representing size and texture traits. High score values depict larger ants and low score values depict smaller ants. **(B)** Second principal component (PC2) representing body coloration. High score values depict dark-brown ants, middle scores depict light-brown ants, and low score values depict black ants. **(C)** Third principal component (PC3) representing number of spines. High score values depict spiny ants, while low score values depict non-spiny ants. Groups not connected by the same letter are significantly different.

## Discussion

We found that *O. sylvatica* alkaloid profiles varied between populations, corresponding with changes in the availability of leaf litter ants along a geographical gradient of temperature, precipitation, and altitude. Our results align with previous studies (McGugan et al., 2016; Moskowitz et al., 2020; Myers & Daly, 1976; Prates et al., 2019; Saporito et al., 2006; Stuckert et al., 2014), and provide further evidence of the importance of environmental availability of alkaloid containing prey in shaping the chemical repertoire in poison frogs. Overall, diablito populations with higher alkaloid loads were found at cooler, high-elevation sites, where leaf litter ant community composition was more diverse, except in La Maná, where both frog alkaloids and leaf litter ants were low despite the high altitude. As altitude and temperature vary along geographical gradients, they can drive changes in frog alkaloid profiles indirectly by shaping the composition and diversity of their arthropod prey (Brühl et al., 1999; Mackay et al., 1986; Moses et al., 2021; Silva & Brandão, 2014; Wise & Lensing, 2019). This is consistent with our previous work where we showed that alkaloid profiles, diet, and surrounding leaf litter communities in the diablito population from Santo Domingo were more abundant in frogs from a cooler, humid forest than in a hot, dry pasture (Moskowitz et al., 2020). Other factors like chemical diversity of arthropod prey have been shown to influence alkaloid variability in diablito frogs (McGugan et al., 2016). Future work comparing environmental arthropod chemistry is necessary to better understand the interplay between chemical repertoire and environmental availability of prey in organisms with diet-acquired defenses.

We found that *H. infraguttatus*, which is typically considered chemically ‘undefended’, has lower yet detectable amounts of alkaloids in the skin, compared to the sympatric chemically defended *O. sylvatica*. Our results align with the hypothesis that ‘undefended’ dendrobatids can accumulate alkaloids even in small amounts, and that the ability to acquire alkaloids for defense may be more common across dendrobatids (Tarvin et al., 2024). In this study histrionicotoxins and indolizidines made up the highest proportion of the alkaloid profiles in both *O. sylvatica* populations and *H. infraguttatus*, which correspond with what has been found in other dendrobatids, including *Oophaga* and *Hyloxalus* frogs (Jeckel et al., 2019; McGugan et al., 2016; Moskowitz et al., 2020; Saporito et al., 2006; Tarvin et al., 2024). Few studies have shown the variability of individual alkaloids and their influence on frog fitness. The potency or organismal effect of most poison frog alkaloids is unknown and there are debates about which assays are most appropriate to test potency (Weldon, 2017). Some poison frog alkaloids are toxic, while other alkaloids are more noxious (Daly et al., 2005; Nayik & Kour, 2022; Santos et al., 2016). Birds avoided eating food associated with skin secretions from one *Dendrobates tinctorius* population and this difference was partly explained by 15 alkaloids, many of them indolizidines (Lawrence et al., 2023). Given the variation in *O. sylvatica* alkaloid profiles, it is likely that there are differences in palatability to predators, although more studies on alkaloid ingestion with natural predators are needed to understand this relationship.

Organisms with diet-derived defenses must maintain their defensive reservoirs through balancing food choice and availability. In our study, sympatric groups of defended *O. sylvatica* ate more ants and mites than *H. infraguttatus,* which ate a less specialized diet of ants, beetles and arthropods gathered in the ‘other’ category, demonstrating dietary differences between species with similar prey access. Additionally, although *O. sylvatica* differed overall in diet and alkaloid profiles across populations, the majority of their stomach contents were constituted by ants, consistent with previous studies (McGugan et al., 2016; Moskowitz et al., 2020). Our results are also consistent with a broader dietary study of dendrobatids in which chemically defended species, such as *Epipedobates anthonyi, Dendrobates auratus, O. pumilio,* showed a greater degree of dietary specialization on ants than ‘undefended’ species, such as *Allobates femoralis, Allobates zaparo, Hyloxalus infraguttatus* (Caldwell, 1996; Darst et al., 2005). However, some cryptically colored dendrobatid species eat mainly ants and some chemically defended species have a broader diet than expected (Darst et al., 2005; Tarvin et al., 2024; Toft, 1995). Our data is consistent with growing evidence that ants make up a large portion of the diet in many *Hyloxalus* species (Darst et al., 2005; Sánchez Loja et al., 2023; Tarvin et al., 2024). Overall, our results support the broad trend of high alkaloid-bearing dendrobatids being ant and mite specialists, while providing further evidence that poison frogs with low alkaloid levels also consume important amounts of ants.

Consuming specific arthropod prey at rates disproportionate to their availability can influence poison frogs’ alkaloid profile, as certain alkaloid classes have known origins in specific arthropod taxa (Blum et al., 1980; McGugan et al., 2016; Santos et al., 2016; Saporito, Donnelly, et al., 2007; Saporito et al., 2004; Spande et al., 1999). In all *O. sylvatica* populations, frogs consistently selected *Solenopsis* ants, while *Strumigenys*, *Paratrachymyrmex*, *Crematogaster, Pheidole* and *Cyphomyrmex* ants were selectively consumed in specific populations, a pattern consistent with our previous findings (McGugan et al., 2016; Moskowitz et al., 2020). These ant genus are known sources of several alkaloid classes particularly abundant across diablito populations, including histrionicotoxins, decahydroquinolines, 3,5-disubstituted indolizidines, and pyrrolidines (Blum et al., 1980; Jones et al., 1982, 1999; McGugan et al., 2016; Moskowitz et al., 2020; Spande et al., 1999). Dietary selectivity in *O. sylvatica* may suggest a preference for specific ant prey based on their alkaloid content, or it may simply reflect that these frogs inhabit microhabitats where alkaloid-rich ants are abundant, leading to incidental consumption without active behavioral preference, as previously observed in *O. pumilio*. (Donnelly, 1991). Further behavioral assays are required to distinguish between these competing hypotheses. Additionally, given the varied, but overall high number of mites recovered from stomach contents across localities, our data suggest that mites are also an important defensive alkaloid source for *O. sylvatica*, probably of 5,8-disubstituted & 5,6,8-trisubstituted indolizidines and pumiliotoxins, as previously found in *O. pumilio* (Saporito et al., 2006; Saporito, Donnelly, et al., 2007). Mite taxonomy and chemistry is drastically understudied compared to ants and future studies should also make efforts to include mites in their analyses (but see Saporito, Donnelly, et al. 2007; Saporito et al. 2011; Saporito et al. 2015).

We found that *H. infraguttatus* selected *Pheidole* ants, which were also selected by *O. sylvatica* from the ‘Cristobal Colón’ locality within this study. Previous alkaloid sampling of *O. sylvatica* and *Pheidole* ants found evidence of overlapping alkaloids within the same sampling location in both frogs and ants (Moskowitz et al., 2020). Our data suggests that selection for alkaloid-rich ants is not exclusive to aposematic species. Recent evidence found that *Pheidole* ants constituted the largest portion of the diet in the cryptic *Allobates femoralis* frog, although they were not actively selected (Sanches et al., 2023). It is unclear whether *H. infraguttatus* frogs have less alkaloids because their diet is less ant and mite rich or if they lack the physiological mechanisms required for alkaloid sequestration in higher concentrations, as has been noted recently in other cryptic species (Alvarez-Buylla et al., 2023). For example, in laboratory toxin feeding trials *A. femoralis* was able to uptake alkaloids as effectively as *O. sylvatica*, but exhibited signs of physiological distress (Caty et al., 2025). Further captive feeding experiments would be helpful to fully understand the implications of dietary selectivity for the evolution of acquired chemical defenses among poison frogs.

Poison frogs may selectively consume certain ant genera based on their alkaloid content, reflecting underlying feeding preferences. However, dietary selectivity can also arise from factors beyond behavioral preference. For instance, *Solenopsis* and *Pheidole* ants may be consumed more frequently, regardless of their abundance, because they are smaller and require less handling effort than bigger, more aggressive ant genera, as observed in the chemically defended frog, *Chiasmocleis leucosticta* (Meurer et al., 2021). In contrast, the Neotropical toad, *Rhinella alata*, selectively consumed larger, more conspicuous ants despite their rarity, while avoiding the more abundant but smaller *Solenopsis* ants (McElroy and Donoso, 2019). Also, as we limited our genus-level analysis to ants here, we cannot determine whether frogs select ants over other prey categories that may offer different metabolic benefits or higher nutritional value. For example, when prey availability is controlled under lab conditions, lab-raised *Dendrobates tinctorius* frogs gravitate to the most nutritious prey (larvae), regardless of size and other prey options including ants (Moskowitz et al., 2022). A careful comparison of frog diet with all available arthropods in the leaf litter through metabarcoding could be useful to determine whether behavioral selectivity is consistent across prey categories. Although we found solid evidence on the relationship between alkaloid defenses, frog diets and arthropod availability, it is worth noting that alkaloid profiles in poison frogs reflect long-term accumulation, whereas dietary composition and prey availability are measured at a single time point, potentially capturing only a subset of the frogs’ long-term foraging patterns. Finally, we collected leaf litter samples with Winkler sacs, which often capture abundant small, specialist myrmicines compared to pitfall traps, which are better in capturing bigger scavengers like *Anochetus* or *Hypoponera* ants (Olson, 1991; Sabu & Shiju, 2010). Integrating multiple sampling methods may provide a more ecologically realistic view of the arthropod community available to poison frogs, including variation in foraging patterns that could underestimate prey availability.

## Summary

Our study demonstrates that dietary preferences and environmental prey availability shape poison frog chemical defenses. *Oophaga sylvatica* showed alkaloid profiles that varied with local leaf litter ant availability, while the sympatric *Hyloxalus infraguttatus*, traditionally considered ‘undefended’, carried small but detectable alkaloid levels. Both species selectively consumed certain small ant genera, suggesting toxic prey selection is not exclusive to aposematic species. These findings highlight dietary selectivity as a key factor in the evolution of chemical defenses. Future studies should replicate these findings in both natural and controlled settings, compare environmental arthropod chemistry to better understand prey availability, and use captive feeding experiments to test how diet influences alkaloid accumulation.

## Supporting information

reproducible report

## Acknowledgements

We thank Lola Guarderas (Wikiri) for logistical and resource support in Ecuador. We are thankful to the Peter Dorrestein lab at The University of California at San Diego for creating the GNPS environment that allows us to perform the metabolomic analysis of poison frog alkaloids. We would also like to thank Mabel Gonzalez, who was a visiting researcher in the Dorrestein Lab and helped us implement GNPS in our lab. LAC acknowledges the support of the Saint Louis Zoo.

We acknowledge that our work at Stanford University takes place on the ancestral and unceded land of the Muwekma Ohlone tribe.

## Data accessibility

Diet and alkaloid data is included in supplementary materials. Gas chromatography / mass spectrometry files and arthropod photos are available on DataDryad (submission pending acceptance).

## Author contributions

Conceptualization: NAM, LAO

Methodology: NAM, LAO, DAD, AAB

Formal Analysis: NAM, AAB,CR

Investigation: NAM, AAB, AC, JR, DAD

Resources: LAO, DAD

Data Curation: NAM, AAB, CM, DAD, CR

Writing – original draft preparation: NAM, LAO, AAB, DAD

Writing – review and editing: NAM, LAO, AAB, DAD, ET, AC, JR, LAC, CR, KF

Visualization: NAM, LAO, CR

Supervision: LAO

Project administration: LAO

Funding acquisition: LAO

---

**Table S1.**
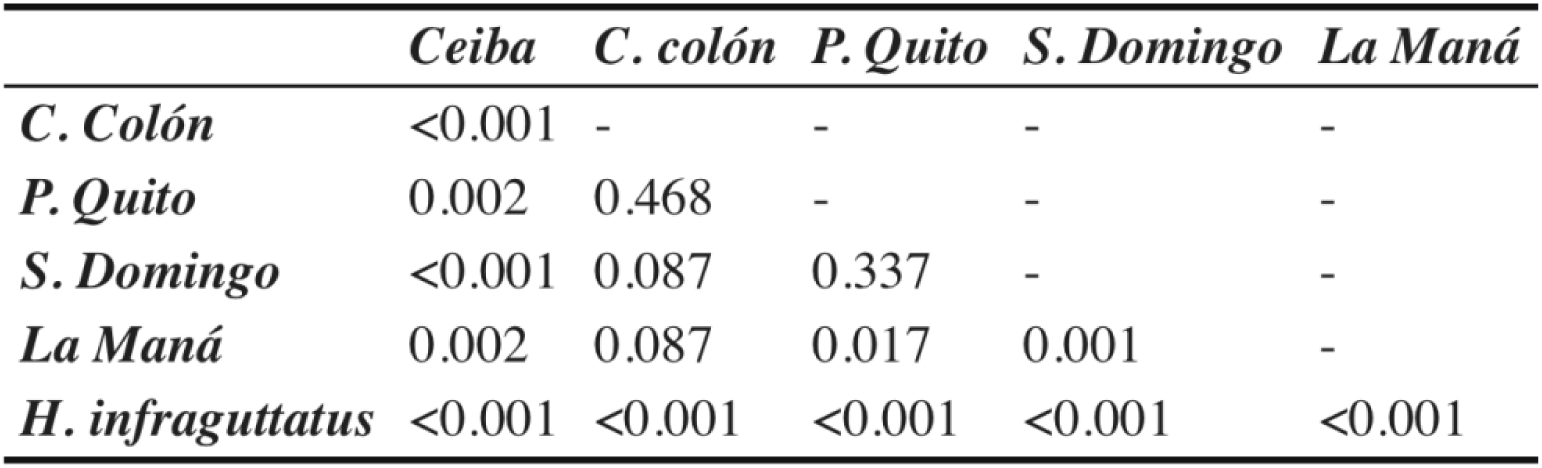
P-values of pairwise Wilcoxon test on differences in summed alkaloids between frog populations.

**Table S2.**
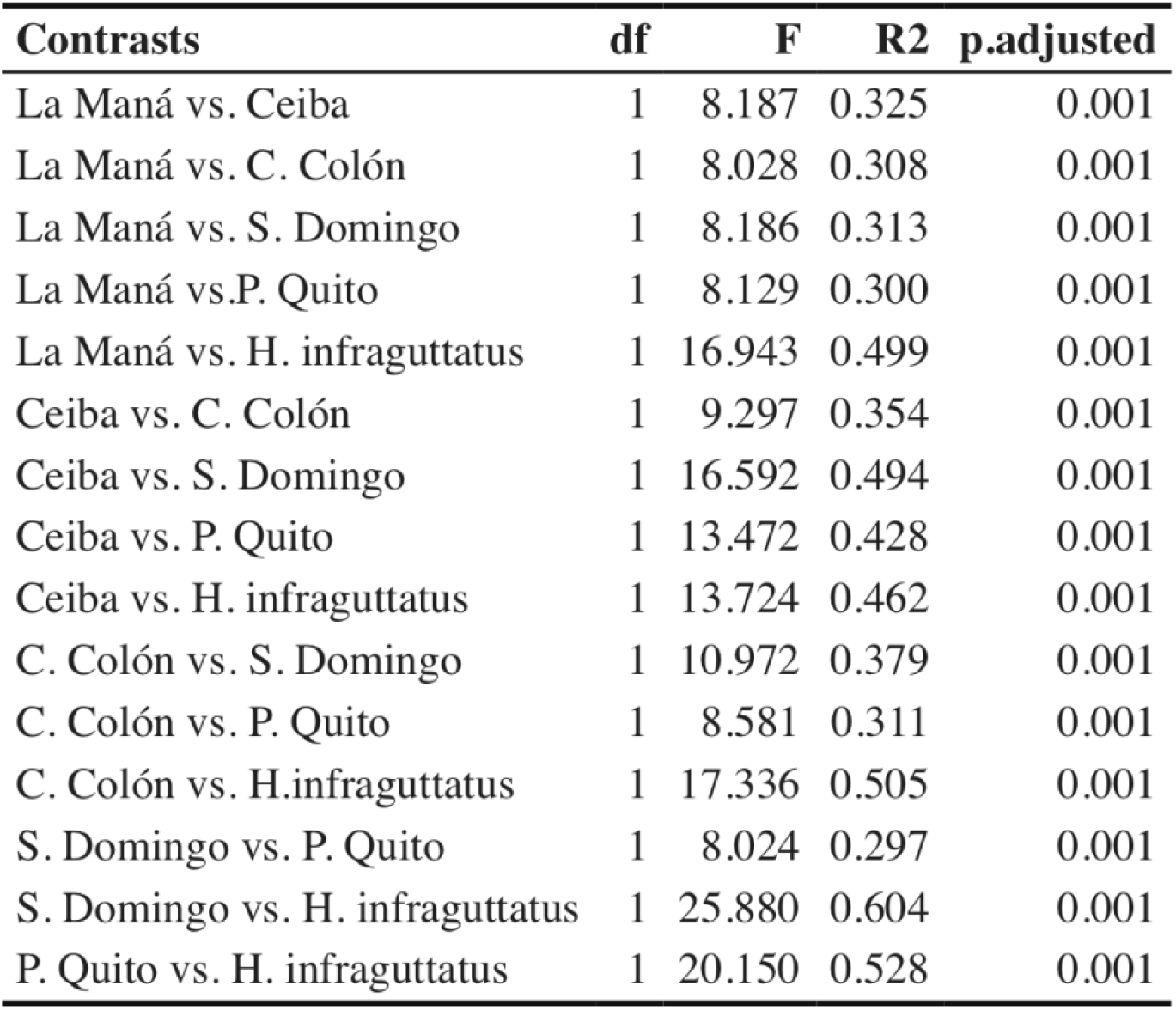
Summary of the results of pairwise comparisons of a PERMANOVA in alkaloid composition between *O. sylvatica* populations.

**Table S3.**
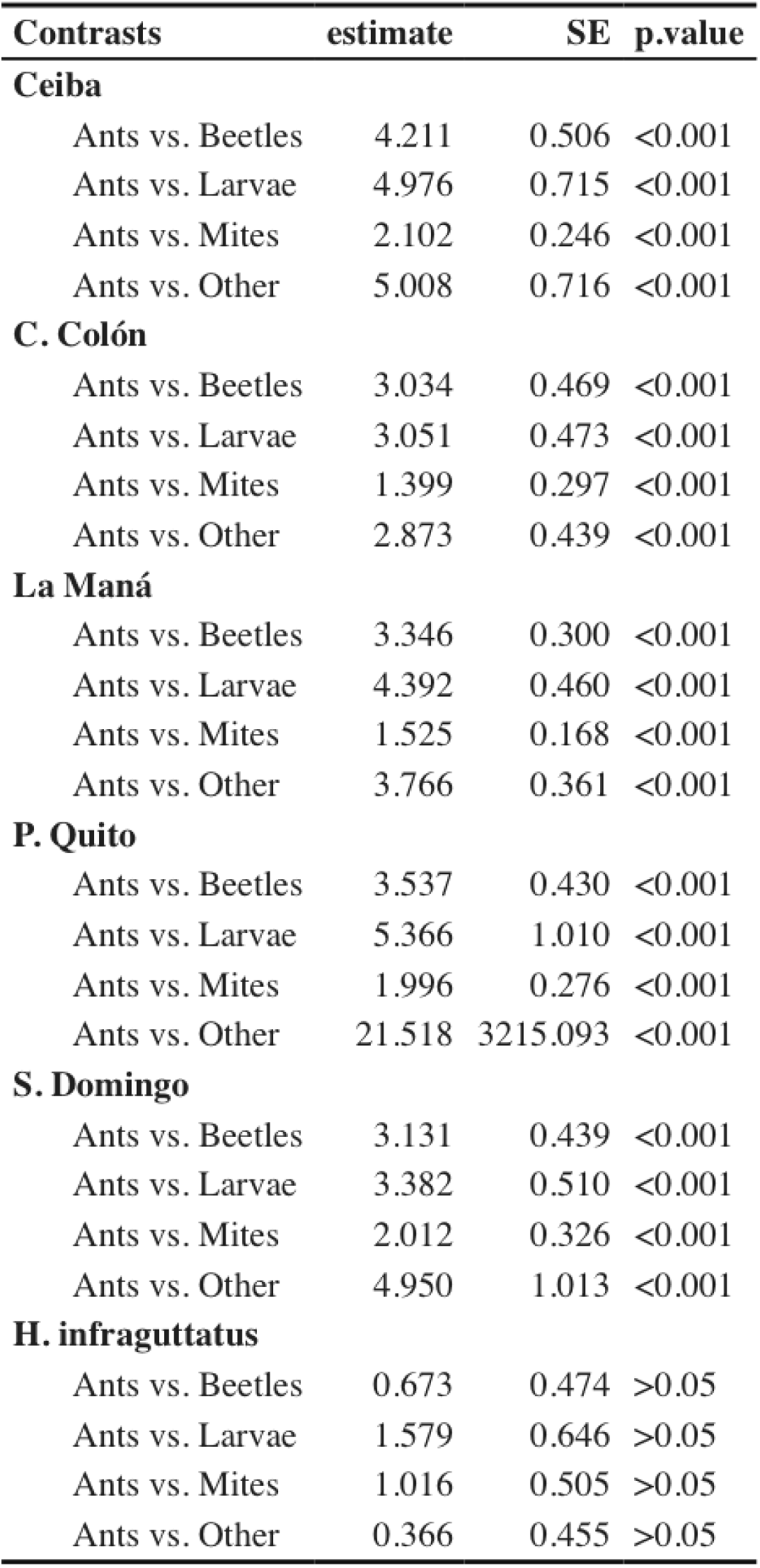
Summarized results of estimated marginal means between frog species and prey items. P-values were adjusted using Tukey’s method.

| **Contrasts** | **estimate** | **SE** | **p.value** |
| --- | --- | --- | --- |
| **Ceiba** |  |  |  |
| Ants vs. Beetles | 4.211 | 0.506 | <0.001 |
| Ants vs. Larvae | 4.976 | 0.715 | <0.001 |
| Ants vs. Mites | 2.102 | 0.246 | <0.001 |
| Ants vs. Other | 5.008 | 0.716 | <0.001 |
| **C. Colón** |  |  |  |
| Ants vs. Beetles | 3.034 | 0.469 | <0.001 |
| Ants vs. Larvae | 3.051 | 0.473 | <0.001 |
| Ants vs. Mites | 1.399 | 0.297 | <0.001 |
| Ants vs. Other | 2.873 | 0.439 | <0.001 |
| **La Maná** |  |  |  |
| Ants vs. Beetles | 3.346 | 0.300 | <0.001 |
| Ants vs. Larvae | 4.392 | 0.460 | <0.001 |
| Ants vs. Mites | 1.525 | 0.168 | <0.001 |
| Ants vs. Other | 3.766 | 0.361 | <0.001 |
| **P. Quito** |  |  |  |
| Ants vs. Beetles | 3.537 | 0.430 | <0.001 |
| Ants vs. Larvae | 5.366 | 1.010 | <0.001 |
| Ants vs. Mites | 1.996 | 0.276 | <0.001 |
| Ants vs. Other | 21.518 | 3215.093 | <0.001 |
| **S. Domingo** |  |  |  |
| Ants vs. Beetles | 3.131 | 0.439 | <0.001 |
| Ants vs. Larvae | 3.382 | 0.510 | <0.001 |
| Ants vs. Mites | 2.012 | 0.326 | <0.001 |
| Ants vs. Other | 4.950 | 1.013 | <0.001 |
| **H. infraguttatus** |  |  |  |
| Ants vs. Beetles | 0.673 | 0.474 | >0.05 |
| Ants vs. Larvae | 1.579 | 0.646 | >0.05 |
| Ants vs. Mites | 1.016 | 0.505 | >0.05 |
| Ants vs. Other | 0.366 | 0.455 | >0.05 |

**Table S4.**
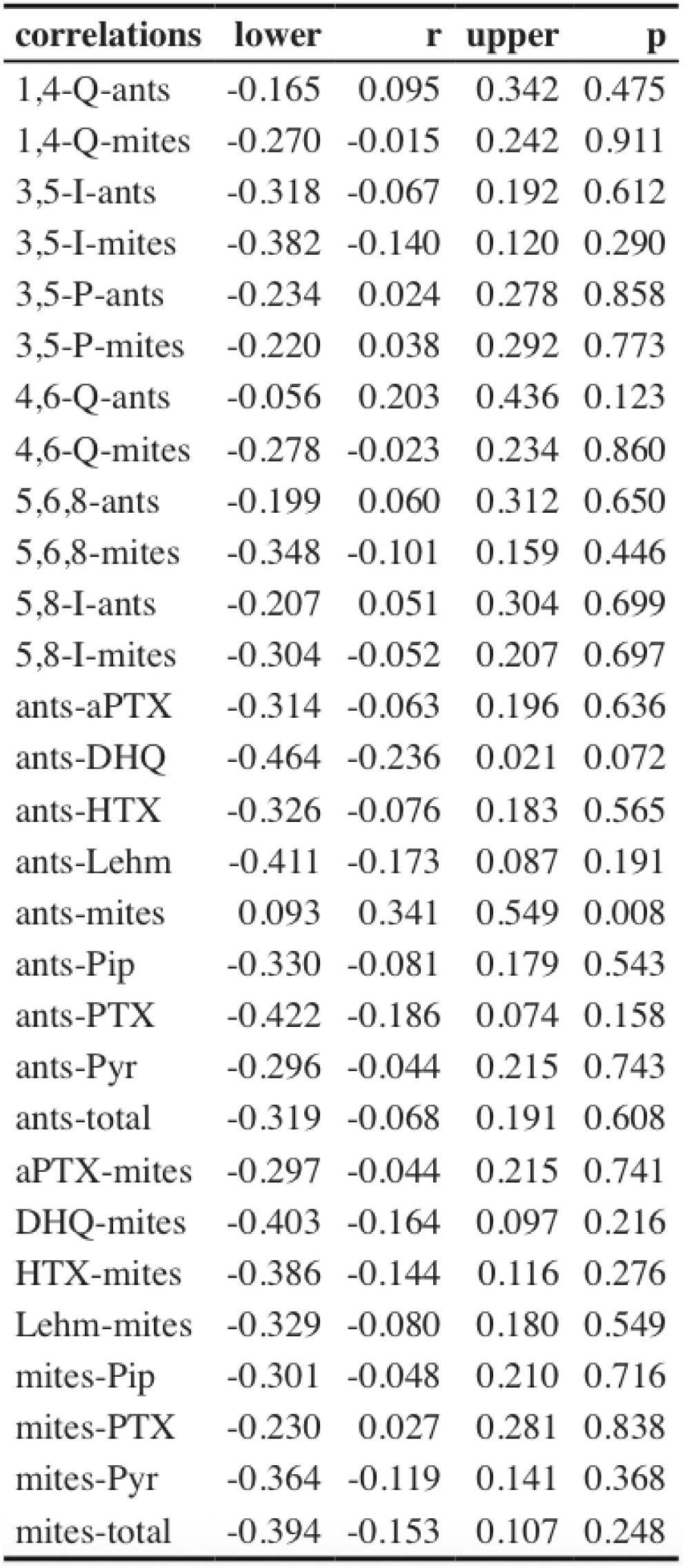
Summary of the results of pairwise correlations between each alkaloid class and ant & mite abundance.

**Table S5.**
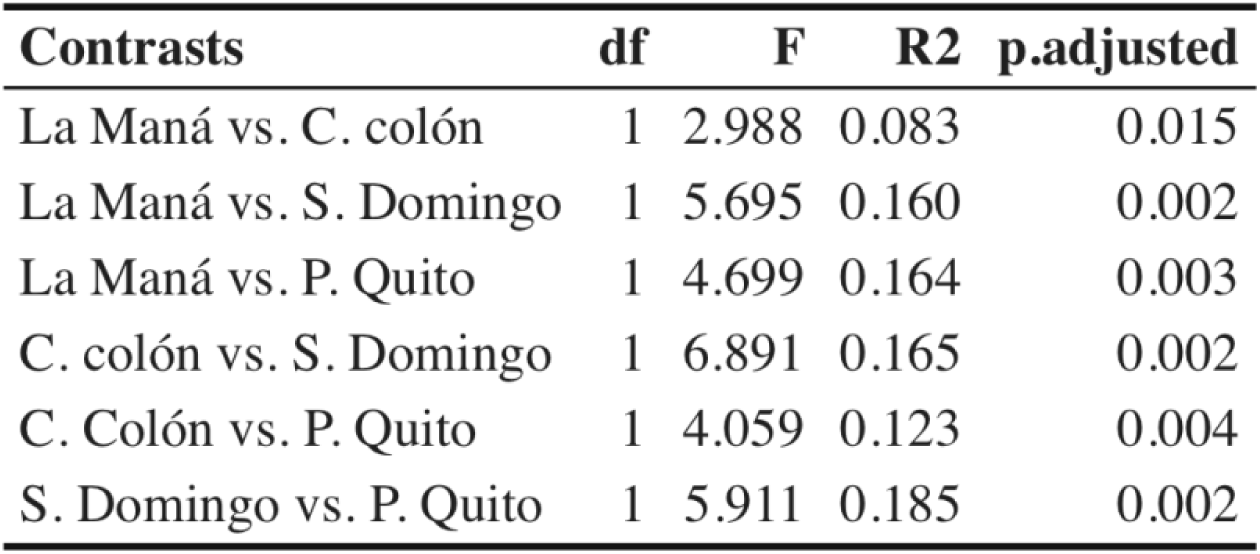
Summary of the results of pairwise comparisons of a PERMANOVA in leaf litter ant composition between study sites. P-values were adjusted using Tukey’s method.

| **Contrasts** | **df** | **F** | **R2** | **p.adjusted** |
| --- | --- | --- | --- | --- |
| La Maná vs. C. colón | 1 | 2.988 | 0.083 | 0.015 |
| La Maná vs. S. Domingo | 1 | 5.695 | 0.160 | 0.002 |
| La Maná vs. P. Quito | 1 | 4.699 | 0.164 | 0.003 |
| C. colón vs. S. Domingo | 1 | 6.891 | 0.165 | 0.002 |
| C. Colón vs. P. Quito | 1 | 4.059 | 0.123 | 0.004 |
| S. Domingo vs. P. Quito | 1 | 5.911 | 0.185 | 0.002 |

**Table S6.**
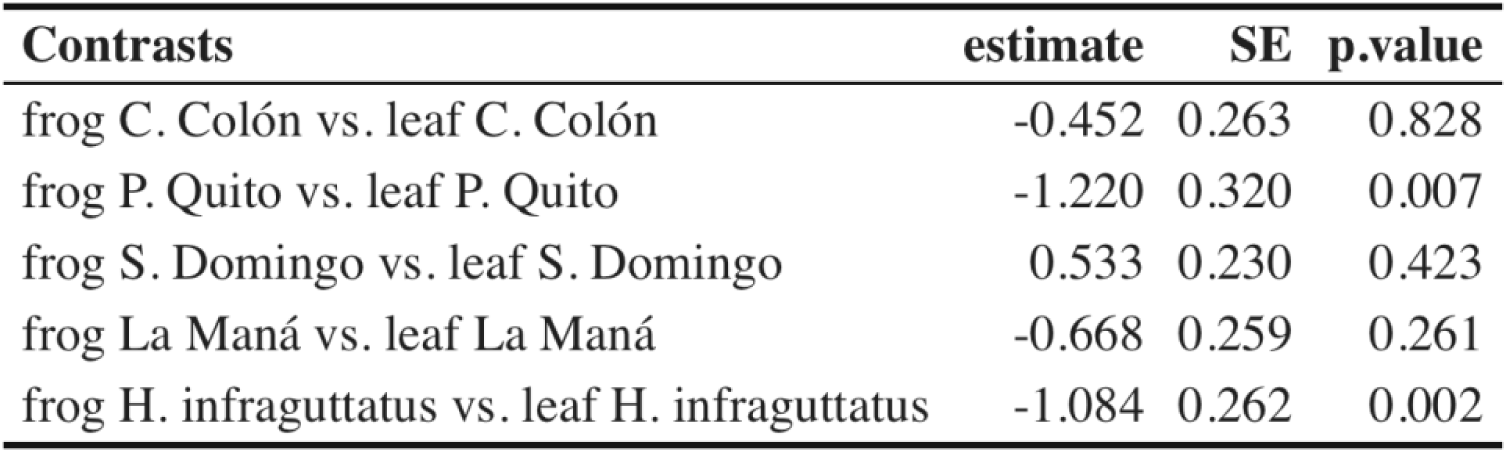
Summary of the results of pairwise comparisons of a Negative Binomial regression comparing ant abundance between *O. sylvatica* populations. P-values were adjusted using Tukey’s method.

**Table S7.**
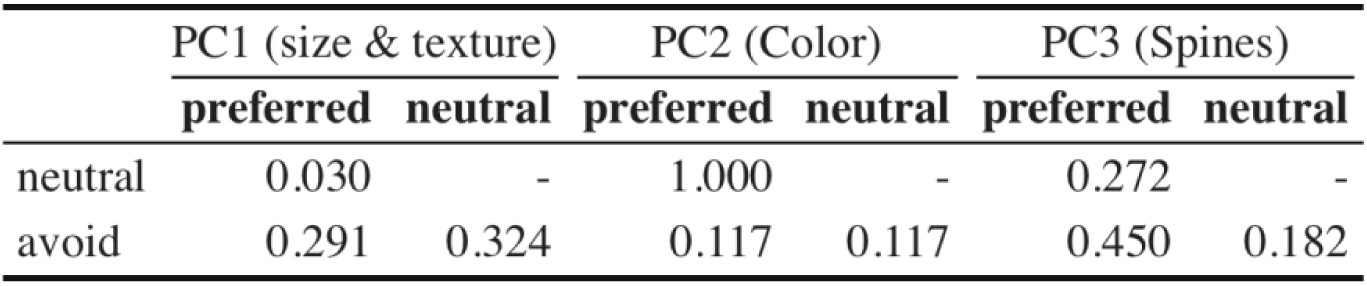
P-values of pairwise Wilcoxon test on differences in principal components between linear selectivity categories.

|  | PC1 (size & texture) |  | PC2 (Color) |  | PC3 (Spines) |  |
| --- | --- | --- | --- | --- | --- | --- |
|  | preferred | neutral | preferred | neutral | preferred | neutral |
| neutral | 0.030 | - | 1.000 | - | 0.272 | - |
| avoid | 0.291 | 0.324 | 0.117 | 0.117 | 0.450 | 0.182 |

