## Supplementary material for "Poison frog chemical defenses are influenced by environmental availability and dietary selectivity for ants": reproducible report: Reproducible report_rev.html


#### Table of contents

- Abstract
- Introduction
- Methods
  - Study system and sample collection
  - Alkaloid extraction and quantification
  - Frog stomach contents identification
  - Leaf litter communities and ant morphology
  - Data Analysis
    - Alkaloid comparisons
    - Frogs’ diet comparisons
    - Ant- vs mite -derive alkaloids
    - Skin alkaloids vs. leaf litter ant communities along a geographical gradient
    - Stomach vs. leaf litter ant communities and frogs’ preference for ant genera
    - Ant morphology vs frog selectivity
- Results
  - Alkaloids differ between species and across diablito frog populations
  - Defended frogs consumed more ants relative to other prey types and compared to the diet of the undefended species
  - Alkaloid diversity among sites is associated with variation in leaf litter ant communities
  - Frogs show different dietary selectivity for particular ant genera
  - Ant morphology influences prey selectivity in frogs
- Discussion
- Summary
- Acknowledgements
  - Data accessibility
  - Funding
  - Author contributions
- References

### Poison frog chemical defenses are influenced by environmental availability and dietary selectivity for ants

 Code

- Show All Code
- Hide All Code
- ---
- View Source

Nora A. Martin†¹, Camilo Rodríguez†¹, Aurora Alvarez-Buylla¹, Katherine Fiocca¹, Colin R. Morrison², Adolfo Chamba-Carrillo³, Ana B. García-Ruilova⁴, Janet Rentería⁵, Elicio E. Tapia⁶, Luis A. Coloma⁶, David A. Donoso\*⁷⁺⁸, Lauren A. O’Connell\*¹

¹ Department of Biology, Stanford University, Stanford, CA 94305, USA  
 ² Department of Integrative Biology, The University of Texas at Austin, Austin, TX 78712, USA  
 ³ Programa de Posgrado en Biodiversidad y Cambio Climático, Universidad Indoamérica, Quito, Ecuador  
 ⁴ División de Entomología, Instituto Nacional de Biodiversidad, Pje. Rumipamba N. 341 y Av. de los Shyris, Quito, Ecuador  
 ⁵ School of Biological Sciences, University of Bristol, Bristol, UK   
 ⁶ Centro Jambatu de Investigación y Conservación de Anfibios, Fundación Otonga, San Rafael, Quito, Ecuador  
 ⁷ Departamento de Biología, Escuela Politécnica Nacional, Ladrón de Guevara E11-253, Quito, Ecuador  
 ⁸ Grupo de Investigación en Ecología y Evolución en los Trópicos -EETrop-, Universidad de las Américas, Quito, Ecuador

Figure 1: Map of frog populations and experimental workflow. **(A)** Collection sites are shown on a topographic map of western Ecuador. Note that both *Hyloxalus infraguttatus* and *Oophaga sylvatica* were collected from La Maná. m.a.s.l. = meters above the sea level. **(B)** A flowchart depicts the main steps of data collection for frog stomach content and leaf litter samples, in order to compare ant genera abundances between these groups.

#### Data Analysis

All statistics and figures were generated in R Studio (version 1.1.442) running R (version 3.5.2).

Packages

```
library(Hmisc)
library(corrr)
library(corrplot)
library(bipartite)
library(sjPlot)
library(sjmisc)
library(sjlabelled)
library(sjtable2df)
library(caret)
library(MASS)
library(ggplot2)
library(emmeans)      ## estimated marginal means (least-squares means)
library(lmerTest)     ## p-values for lme4 models
library(lme4)         ## Mixed models 
library(lmtest)
library(MuMIn)
library(ggpubr)
library(nlme)
library(GGally)
library(PerformanceAnalytics)
library(psych)
library(knitr)
library(performance)
library(see)
library(viridis)       ## Viridis colour palette
library(car)
library(mvnTest)
library(rstatix)       ## different analyses (e.g., pairwise.t.test, Dunn's test)
library(FSA)           ## alternative for Dunn's test
library(repmis)
library(tidyverse)     ## the tidyverse
library(vegan)         ## Ordination, dissimilarity analyses
library(colorspace)    ## adjust colors
library(rcartocolor)   ## Carto palettes
library(ggforce)       ## sina plots
library(ggdist)        ## halfeye plots
library(ggridges)      ## ridgeline plots
library(ggbeeswarm)    ## beeswarm plots
library(gghalves)      ## off-set jitter
library(systemfonts)   ## custom fonts
library(kableExtra)
library(ggh4x)
library(ape)            
library(wesanderson)    ## Colour palette
library(grid)
library(png)
library(devtools)
library(pairwiseAdonis)
library(reshape2)       ## function melt
library(readr)          ## to export kable
library(rptR)           ## Repeatability
library(DescTools)
library(igraph)
library(remotes)
library(tinytable)
library(stats)
library(ggcorrplot)
library(glmmTMB)
library(DHARMa)
library(dietr)
library(FactoMineR)
library(factoextra)
library(modeldb)
library(ade4)
library(conflicted)
library(ggsankey)
library(raster)
library(circlize)
library(scico)
library(magrittr)

conflicts_prefer(dplyr::select)# Will prefer dplyr::select over any other package. 

conflicts_prefer(dplyr::filter)# Will prefer dplyr::filter over any other package. 

conflicts_prefer(base::attr)

conflicts_prefer(dplyr::summarize)

conflicts_prefer(dplyr::mutate)

conflicts_prefer(dplyr::rename)

conflicts_prefer(dplyr::summarise)

conflicts_prefer(base::union)

conflicts_prefer(magrittr::set_names)
```

All data sets used for this analyses can be accessed in: https://drive.google.com/open?id=1d0iVHDFW4sX9hHQyasMv9oO-aWF1Ff\_5&usp=drive\_fs

data

```
#LOBSU Mac
feat_tab <- read.csv('/Users/camilorl/Library/CloudStorage//Shared drives/LOBSU Manuscripts/O. sylvatica vs. H. infraguttatus diet (Nora)/Submission 2-JAE/Code&Data/featureTableY_normalized.csv')

#LOBSU Mac 
alkadiver <- read.csv(
  "/Users/camilorl/Library/CloudStorage//Shared drives/LOBSU Manuscripts/O. sylvatica vs. H. infraguttatus diet (Nora)/Submission 2-JAE/Code&Data/alkadiver.csv")

#My Mac 
#alkadiver <- read.csv(
#  '/Users/camilorodriguezlopez/Library/CloudStorage//Shared drives/LOBSU Manuscripts/O. sylvatica vs. H. infraguttatus diet (Nora)/Submission 2-JAE/Code&Data/alkadiver.csv') # has compositional differences in skin alkaloid profiles between *O. sylvatica* populations and *H.infragutatus*, and altitude for every locality.

#LOBSU Mac 
L_antdiver <- read.csv("/Users/camilorl/Library/CloudStorage//Shared drives/LOBSU Manuscripts/O. sylvatica vs. H. infraguttatus diet (Nora)/Submission 2-JAE/Code&Data/L_antdiver.csv")

#My Mac 
#L_antdiver <- read.csv('/Users/camilorodriguezlopez/Library/CloudStorage//Shared drives/LOBSU Manuscripts/O. sylvatica vs. H. infraguttatus diet (Nora)/Submission 2-JAE/Code&Data/L_antdiver.csv')#has the abundance of all ant genera collected from the leaf litter and from frogs' stomach contents in every locality.

#LOBSU Mac 
diet <- read.csv(
  "/Users/camilorl/Library/CloudStorage//Shared drives/LOBSU Manuscripts/O. sylvatica vs. H. infraguttatus diet (Nora)/Submission 2-JAE/Code&Data/diet.csv")

#My Mac 
#diet <- read.csv(
#  '/Users/camilorodriguezlopez/Library/CloudStorage//Shared drives/LOBSU Manuscripts/O. sylvatica vs. H. infraguttatus diet (Nora)/Submission 2-JAE/Code&Data/diet.csv')#has the type and number of prey items found in the stomach contents of frog individuals from all populations.

#LOBSU Mac 
ant_traits <- read.csv("/Users/camilorl/Library/CloudStorage//Shared drives/LOBSU Manuscripts/O. sylvatica vs. H. infraguttatus diet (Nora)/Submission 2-JAE/Code&Data/ant_traits.csv")

#My Mac 
#ant_traits <- read.csv('/Users/camilorodriguezlopez/Library/CloudStorage//Shared drives/LOBSU Manuscripts/O. sylvatica vs. H. infraguttatus diet (Nora)/Submission 2-JAE/Code&Data/ant_traits.csv')#has 17 morphological traits of all ant genera collected from the leaf litter in every locality.


#LOBSU Mac
ant_all <- read.csv("/Users/camilorl/Library/CloudStorage//Shared drives/LOBSU Manuscripts/O. sylvatica vs. H. infraguttatus diet (Nora)/Submission 2-JAE/Code&Data/ant_all.csv")

#My Mac
#ant_all <- read.csv('/Users/camilorodriguezlopez/Library/CloudStorage//Shared drives/LOBSU Manuscripts/O. sylvatica vs. H. infraguttatus diet (Nora)/Submission 2-JAE/Code&Data/ant_all.csv')# has abundance of all ant species captured in pitfall and Winkler traps for every place.
```

##### Alkaloid comparisons

We used a Kruskal-Wallis test to examine overall differences between *O. sylvatica* and *Hyloxalus infraguttatus* populations in summed toxicity across the 79 alkaloids in the 13 structural families. A pairwise Wilcoxon test was used to determine *O. sylvatica* population differences between the 13 structural families in both species, with p-values adjusted for multiple testing using the False Discovery Rate control (FDR).

Code

```
#Kruskal-Wallis test to compare skin alkaloids accross populations
kw_test <- alkadiver %>% 
  select(-latitude, -longitude, -altitude) %>% 
  mutate(total = select(., contains(".")) %>%
           rowSums(na.rm = TRUE)) %>% 
  do(tidy(kruskal.test(.$total ~ .$population)))

###-------------------------##
## NMDS for skin alkaloids ##
##-------------------------##

#the ordination was performed using Bray-Curtis dissimilarities
ord <- metaMDS((alkadiver %>% 
  select(where(is.numeric), -altitude, -latitude, -longitude)), 
  perm=9999, distance = "bray", 
               k = 2, autotransform = FALSE)

#the ordination was ran a second time starting at the previous best solution to ensure the stability and reliability of the results in the NMDS 
ord2<-metaMDS((alkadiver %>% 
  select(where(is.numeric), -altitude, -latitude, -longitude)), previous.best = TRUE)


#Include environmental variables 

# Create a vector with the "environmental variables" 
#For this I will use the coordinates of every locality and extract average historical temperature and precipitation, in addition to altitude, from World Clim.  

# Load average temperature and precipitation rasters from : https://www.worldclim.org/data/worldclim21.html 

# Adjust this path accordingly:

#My_Mac
#prec <- stack('/Users/camilorodriguezlopez/Library/CloudStorage//Shared drives/LOBSU Manuscripts/O. sylvatica vs. H. infraguttatus diet (Nora)/Submission 2-JAE/Code&Data/wc2.1_30s_prec_05.tif')  # layer for May
#temp <- stack('/Users/camilorodriguezlopez/Library/CloudStorage//Shared drives/LOBSU Manuscripts/O. sylvatica vs. H. infraguttatus diet (Nora)/Submission 2-JAE/Code&Data/wc2.1_30s_tavg_05.tif')  # layer for May

#LOBSU_Mac
prec <- stack('/Users/camilorl/Library/CloudStorage//Shared drives/LOBSU Manuscripts/O. sylvatica vs. H. infraguttatus diet (Nora)/Submission 2-JAE/Code&Data/wc2.1_30s_prec_05.tif')  # layer for May
temp <- stack('/Users/camilorl/Library/CloudStorage//Shared drives/LOBSU Manuscripts/O. sylvatica vs. H. infraguttatus diet (Nora)/Submission 2-JAE/Code&Data/wc2.1_30s_tavg_05.tif')  # layer for May

# Coordinates for every locality
coords <- alkadiver %>% 
  rename(lon = longitude , lat = latitude) %>% 
  select(lon, lat)

# Define a point (longitude, latitude)
coordinates(coords) <- ~lon + lat

# Extract temperature and precipitation for May (when the sampling was made) at the specified location
temp_value <- raster::extract(temp, coords)
prec_value <- raster::extract(prec, coords)

# Add location and altitude
env.vars <- alkadiver %>%
  select(altitude) %>% 
  bind_cols(data.frame(temp_value), data.frame(prec_value)) %>% 
  rename(Avtemp = wc2.1_30s_tavg_05, Avprec = wc2.1_30s_prec_05)

# fit environmental variables in the ordination
en <- envfit(ord2, env.vars, perm = 999) # We can see that altitude, latitude and longitude, are significantly correlated with the dimensions 


# fit alkaloids in the ordination to see their individual contribution
en.alka <- envfit(ord2, alkadiver %>%
  select(-latitude, -longitude, -altitude) %>%
  mutate(population = fct_relevel(population, "ceiba", "c_colon", "p_quito", "s_domingo", "la_mana", "H_infra")) %>%
  pivot_longer(cols = indo.137:pyrrz.526, names_to = "alkaloid", values_to = "abundance") %>%
  bind_cols(
    feat_tab %>%
      select(family, santos2016_source) %>%
      slice(rep(1:n(), times = 59))
  ) %>%
  select(frog_id,population,family, abundance) %>%
  pivot_wider(
    names_from = family,
    values_from = abundance,
    values_fn = sum,
    values_fill = 0
  ) %>%
    select(where(is.numeric), -frog_id, -population), perm = 999) 

sort(en.alka$vectors$r, decreasing = T) %>% 
  data.frame()

#PERMANOVA
permanova.output<-adonis2(alkadiver[,6:81]~alkadiver$population, 
                          permutations = 9999,method="bray")

#Pairwise adonis
pair.mod <-pairwise.adonis(alkadiver[,6:81],factors=alkadiver$population,
                           p.adjust = "BH")
```

##### Frogs’ diet comparisons

We visualized interactions between arthropod prey taxa and poison frog species in a bipartite network based on the average abundance of arthropod prey for every frog population. We used the species specificity index (*ssi*), as implemented in the **specieslevel()** function of the ‘*bipartite*’ package, to quantify the variability in dietary interactions between arthropod prey taxa and poison frog species in a bipartite network. This index reflects the degree to which each frog species interacts unevenly across prey taxa, with values ranging from 0 (indicating low variability and generalist behavior) to 1 (indicating high variability and specialist behavior). Additionally, we used generalized linear mixed models (GLMM) to test for compositional differences of frog diet categories using the function **glmmTMB()** within the package ‘*glmmTMB*’ (Brooks et al. 2017). We used a negative binomial distribution appropriate for count data with overdispersion. We tested for diet differences by including species/population, prey type, and their interaction as main effects. Frog individual tags were included as a random variable to account for repeated sampling of prey categories within individuals. We computed estimated marginal means to test for pairwise comparisons between populations using the ‘emmeans’ package (Lenth 2025).

Code

```
# to calculate the mean abundance of consumed prey for every frog population

sum_diet <- diet %>% # [-311,] remove the frog with 18 ants 
  group_by(population, prey) %>% 
  summarise(count_mean = mean(count)) %>%
  pivot_wider(id_cols = population, names_from = prey, 
              values_from = count_mean) %>% 
  data.matrix()


rownames(sum_diet) <- c("ceiba", "c.colon", "h.infragutatus", "mana",  
                         "p.quito", "s.domingo")

sum_diet <- sum_diet[,-c(1)]

# To calculate the specialization index
#dprime <- dfun(sum_diet, abuns=c(1,1,1,1,1))$dprime #I set hypothetical abundances of items in the leaf litter because we don't have the real information.
#However, even if I double the abundance of ants "abuns=c(2,1,1,1,1)", the specialization index is still higher for sylvatica and very low for infra.

#I now use the species specificity index, which which measures the variability in interaction strengths for a species, normalized to range from 0 (Low specificity - interactions are evenly distributed across all partners, i.e., generalist behavior) to 1 (High specificity - interactions are concentrated on a few partners, i.e., specialist behavior)

ssi <- specieslevel(sum_diet, index = "species specificity")

#To test for species differences (O. sylvatica vs. H. infraguttatus) in the number of prey items consumed

sd1 <- glmmTMB(count ~ population*prey,
               family=nbinom1(link="log"), data=diet)

anova.sd1 <- glmmTMB:::Anova.glmmTMB(sd1, typr="II")

#Check model fit by testing overdispersion 
disp.test <- function() {testDispersion(sd1)} #Remove "function() {}"
simulationOutputspp <- simulateResiduals(fittedModel = sd1, plot = F)
test.plots <- function() {plot(simulationOutputspp)} #Remove "function() {}"

#Calculate pairwise comparisons between prey items, within populations 
emm1 <- data.frame(emmeans(sd1, list(pairwise ~ population*prey), 
                           adjust = "tukey")$`emmeans of population, prey`)

emm1$prey <- factor(emm1$prey, levels = c("ants", "mites", "beetle", "other", "larvae"))

emm1$population <- factor(emm1$population, levels = c("ceiba", "puerto_quito","santo_domingo",  "cristobal_colon", "la_mana_Os", "la_mana_Hi"))

# To get rid of the very high values of puerto quito - other prey items
emm1$emmean[emm1$emmean < -10] <-  0
emm1$asymp.LCL[emm1$asymp.LCL < -10] <-  0
emm1$asymp.UCL[emm1$asymp.UCL > 10] <-  0
```

##### Ant- vs mite -derive alkaloids

To infer potential dietary origins of frogs’ skin alkaloids, we cross-referenced the structural classes of alkaloids with the determined arthropod source reported by (Juan C. Santos, Tarvin, and O’Connell 2016), which assigns compounds as derived from ants, mites, or both. We tested whether the proportion of ant-based alkaloids is greater than the proportion of mite-based alkaloids across frog populations using an anova followed by a Tukey post-hoc test for multiple comparisons. We visualized the proportion of ant- and mite-derived alkaloids using a chord diagram. Additionally, we tested whether total alkaloid abundance, as well as abundance within structural families, correlated with the number of ants and mites consumed across populations.

Code

```
## Test if the proportion of ant-based alkaloids is greater than the proportion of mite-based alkaloids across populations

antmitest <- alkadiver %>% 
  select(-latitude, -longitude, -altitude) %>% 
  mutate(total = select(., contains(".")) %>%
           rowSums(na.rm = TRUE)) %>%
  mutate(population = fct_relevel(population, "ceiba", "c_colon", "p_quito", "s_domingo", 
                                  "la_mana", "H_infra")) %>% 
  pivot_longer(cols = c(indo.137:pyrrz.526), 
               names_to = "alkaloid", values_to = "abundance") %>% 
  bind_cols(feat_tab %>% 
              select(family, santos2016_source) %>%
  slice(rep(1:n(), times = 59))) %>% 
  group_by(population, family, santos2016_source) %>% 
  summarize(abundance = sum(abundance)) %>% 
  mutate(family = factor(family, levels = c('HTX', '3,5-I', 'DHQ', 'Pyr', 
                                            '3,5-P', 'Lehm', 'Pip', '5,8-I', 
                                            '5,6,8-I', '1,4-Q', 'PTX', 
                                            '4,6-Q', 'aPTX'))) %>%
  group_by(population, santos2016_source) %>%
  summarise(total_abund = sum(abundance), .groups = "drop") %>%
  pivot_wider(names_from = santos2016_source, 
              values_from = total_abund, values_fill = 0) %>%
  mutate(total = Ants + Mites + Both,
         prop_ant = Ants / total,
         prop_mite = Mites / total,
         prop_both = Both / total) %>% 
  select(population, prop_ant, prop_mite, prop_both) %>% 
  pivot_longer(c(prop_ant:prop_both), names_to = "source", values_to = "proportion") %>%
  aov(proportion ~ source, data = .) %>%
  #plot()# to check residual normality
  #summary()
  TukeyHSD() %>%
  broom::tidy() %>% 
  kable(digits = 3, 
      table.attr = 'data-quarto-disable-processing="true"', "html",
      caption = "Summary of the results of Tukey post-hoc comparisons of an ANOVA in proportion of potential alkaloid arthropod source between frog populations") %>% 
  kable_classic(full_width = F, html_font = "Cambria") %>% 
  row_spec(0, bold = T)

# Correlation between #ants & # mites with alkaloids abundance 
cor_dietalk <- diet %>% 
  filter(prey %in% c("ants", "mites")) %>% 
  pivot_wider(names_from = "prey", values_from = "count", 
              id_cols = c(frog_id, population)) %>% 
  left_join(alkadiver %>% 
              select(-latitude, -longitude, -altitude) %>% 
              mutate(total = select(., contains(".")) %>%
                       rowSums(na.rm = TRUE)), by = "frog_id") %>% 
  na.omit() %>% 
  select(ants, mites, c(indo.137: total)) %>% 
  setNames(c(names(.)[1:2], feat_tab %>% 
               select(family) %>% 
               pull(), "total")) %>%
  {split.default(., names(.))} %>%                    # group columns by name
  map_dfc(~ rowMeans(as.data.frame(.), na.rm = TRUE)) %>% 
  corr.test() %>% 
  .$ci %>%
  rownames_to_column("correlations") %>%
  filter(str_detect(correlations, "mite|ant")) %>% 
  kable(digits = 3, 
      table.attr = 'data-quarto-disable-processing="true"', "html") %>% 
  kable_classic(full_width = F, html_font = "Cambria") %>% 
  row_spec(0, bold = T)
```

##### Skin alkaloids vs. leaf litter ant communities along a geographical gradient

We looked at compositional differences in skin alkaloid profiles of *O. sylvatica* populations and *H. infraguttatus*, and their surrounding leaf litter ant communities in two separate non-metric multidimensional scaling (NMDS), using the function **metaMDS()** within the package ‘*vegan*’ (Oksanen et al. 2024). Statistical differences between and within populations were assessed using a permutational multivariate analysis of variance (PERMANOVA) on Bray-Curtis dissimilarities. P-values for pairwise comparisons were adjusted using the function **pairwise.adonis()**. Additionally, we used the **envfit()** function to understand the influence of altitude, ambient temperature and precipitation on both alkaloid composition and leaf litter ant communities. These environmental variables were selected to capture spatial and temporal variation in ecological conditions that could influence ant availability and frogs’ foraging behavior. Ambient temperature and precipitation data was extracted from WorldClim 2.1, at a spatial resolution of 30 arc-seconds (~1 km²) for each study site (Fick and Hijmans 2017). Similarly, we used envfit() to explore the contribution of specific alkaloid classes and ant genus to the ordination spaces. We used Procrustes analysis to test similarities between the NMDS ordinations of skin alkaloids and ant community composition across sites. The Procrustes correlation was calculated using the **procrustes()** function from the ‘*vegan*’ package, which aligns the two ordinations by scaling, rotating, and translating one configuration to best match the other. Statistical significance of the correlation was evaluated with 999 permutations using the **protest()** function.

Code

```
### Winkler ###
#NOTE: S. Domingo and P. Quito seem to have one outlier each when performing the ordination. I remove them.
ant_co <- (ant_all %>%
  filter(TRAP == "Winkler") %>%
  select(SITE, SPECIES, X., altitude, latitude, longitude) %>% # important to avoid duplicates
  group_by(SITE,SPECIES) %>% # important to avoid duplicates
  mutate(row = row_number()) %>% 
  pivot_wider(names_from = SPECIES, values_from = X.) %>%
  select(-row) %>% 
  replace(is.na(.), 0) %>% 
  mutate(SITE = factor(SITE)) %>% 
  ungroup() %>% 
  mutate(SITE = fct_relevel(SITE, "Mana", "Colon", "Santo Domingo", 
                            "Puerto Quito")) %>%
  arrange(as.integer(SITE)) %>% 
      as.data.frame())[-c(47,65),]
  

#the ordination was performed using Bray-Curtis dissimilarities
ord.co <- metaMDS(ant_co %>%
                     select(where(is.numeric), -altitude, 
                            -latitude, -longitude), 
                  perm=9999, distance = "bray",
                  k = 2, autotransform = FALSE)

  #the ordination was ran a second time starting at the previous best solution to ensure the stability and reliability of the results in the NMDS 
ord.co2<-metaMDS(ant_co %>%
                     select(where(is.numeric), -altitude, 
                            -latitude, -longitude), 
                 previous.best = TRUE)

#Include environmental variables - altitude, latitude and longitude: 
# Create a vector with the "environmental variables" 

coords.2 <- ant_co %>% 
  rename(lon = longitude , lat = latitude) %>% 
  select(lon, lat)

# Define a point (longitude, latitude)
coordinates(coords.2) <- ~lon + lat

# Extract temperature and precipitation for May at the specified location
temp_value.2 <- raster::extract(temp, coords.2)
prec_value.2 <- raster::extract(prec, coords.2)

# Add location and altitude

env.vars.co <- ant_co %>%
  bind_cols(data.frame(temp_value.2), data.frame(prec_value.2)) %>% 
  rename(Avtemp = wc2.1_30s_tavg_05, Avprec = wc2.1_30s_prec_05) %>% 
  select(altitude, Avtemp, Avprec) 

en.co <- envfit(ord.co2, env.vars.co, perm = 999) # We can see that altitude, latitude and longitude, are significantly correlated with the dimensions 

en.co.ants <- envfit(ord.co2, ant_co %>%
         select(where(is.numeric), -altitude,
                -latitude, -longitude), perm = 999)

sort(en.co.ants$vectors$r, decreasing = T) %>% 
  data.frame() 


en.co$vectors$arrows %>%
  data.frame() %>%
  mutate(r2 = en.co$vectors$r) %>% 
  mutate("p-value" = en.co$vectors$pvals) %>% 
  kable(digits = 3,caption = "") %>% 
  kable_classic(full_width = F, html_font = "Cambria") %>% 
  row_spec(0, bold = T) 

#PERMANOVA
permanova.ants<-adonis2((ant_co %>% 
                          select(-SITE,-altitude, -latitude, -longitude)) ~ant_co$SITE, 
                          permutations = 9999,method="bray")

#Pairwise adonis
pair.per.ants <-pairwise.adonis((ant_co %>% select(-SITE,-altitude, -latitude, -longitude)),
                                 factors=ant_co$SITE, p.adjust = "BH")
```

##### Stomach vs. leaf litter ant communities and frogs’ preference for ant genera

To test for within and between species differences in ant abundance between leaf litter and stomach contents, we employed a negative binomial generalized linear model using the **glm.nb()** function from the ‘*MASS*’ package (Venables and Ripley 2002). The model used total ant abundance per sample as the response variable, with group (stomach vs. leaf) and population as predictors. Pairwise comparisons were conducted using estimated marginal means. To examine if frogs show selectivity for consuming specific ant genera, we calculated a linear selectivity index by subtracting the relative abundance of ants found on the leaf litter from those found in frog stomach contents (Strauss 1979). The interpretation of these selectivity values was guided by the methodology in (McElroy and Donoso 2019), which involves generating a null distribution of selectivity values via simulation for each ant species. By comparing the observed selectivity values to this null distribution, we classified the ants as ‘selected’ if the values were above, ‘neutral’ if they were within, and ‘avoided’ if they were below the null distribution, thereby delivering a statistically robust assessment of the frogs’ selective foraging behaviors.

Code

```
# To calculate the total ant abundance per sample only in Winkler sacs
sum_abun <-  L_antdiver %>% 
  rowwise(Trap, group, population) %>% 
  summarise(abundance = sum(c_across(Anochetus:Wasmannia))) %>% 
  filter(!(Trap == "Pitfall"))

#To check normality
shapiro.ants <- shapiro.test(sum_abun$abundance)

Histant <- sum_abun %>% 
  ggplot(aes(x=abundance)) +
  geom_histogram(aes(x=abundance, y=..density..), 
                 bins=5,col="black", fill="black",alpha = 0.15) +
  theme_bw(14) + 
  theme(panel.grid.major = element_blank(), 
        panel.grid.minor = element_blank(), 
        axis.text.x = NULL, legend.position = "none") +
  ylab("Density") + 
  theme(axis.text=element_text(size=9),
        axis.title=element_text(size=11)) +
  annotate("text", x = 250, y = 0.009, label = "Shapiro-Wilk test", 
           size = 3.5) +
  annotate("text", x = 250, y = 0.007, label = "p-value = <0.001", 
           size = 3.5) +
  annotate("text", x = 250, y = 0.005, label = "W = 0.83", 
           size = 3.5)

#### Ant abundance comparison between leafs and stomachs
# We used the data base "L_antdiver"

# Negative binomial logistic regression
sumabun.mod <- glm.nb(abundance ~ group*population, data = sum_abun)
sumabun.aov <- anova(sumabun.mod)

# Estimated marginal means
sumabun.emmeans <- emmeans(sumabun.mod, list(pairwise ~ group*population), 
                           adjust = "tukey")

# Calculate linear selectivity based on (McElroy & Donoso, 2019)

# Function to calculate null distribution 
custom_loop <- function(data) {
  for (i in 1:1000){                                          # number of loops
    j <- sample(2000:5000,1)                                  # number of ants sampled
    randomdraw <- sample(data$Genus,                             # draw species from env. proportion with replacement
                         size = j,
                        replace = TRUE, 
                        prob = data$leafprop)                      
    df_randomdraw <- as.data.frame(table(randomdraw))         # dataframe of species counts
    preyprop_sim <- df_randomdraw$Freq/j                      # vector of simulated prey species proportions
    Linear_sim <- as.data.frame(preyprop_sim - data$leafprop)       # simulated LinearSelectivity by subtracting REAL env_prop from SIMULATEd prey_prop + make dataframe
    colnames(Linear_sim) <- paste0("sim",i)                   # rename column from LinearS_sim --> sim1, sim2, sim3,...sim1000
  
  # add the simulated dataset to the dataframe with species, LinearS, sim1, sim2,...etc....  
    data <- cbind(data,Linear_sim) 
  }
  return(data)
}


# Calculate linear selectivity and null distribution using bootstrap with 1000 iterations 
L_simulations <- L_antdiver %>%
  group_by(group, population) %>% 
  select(where(is.numeric)) %>% 
  summarise_at(vars(Anochetus:Wasmannia), sum) %>%
  rowwise() %>% 
  mutate(across(Anochetus:Wasmannia, ~./sum(c_across(Anochetus:Wasmannia)))) %>%
  ungroup() %>% 
  melt() %>% 
  pivot_wider(names_from = group, values_from = value) %>% 
  rename(frogprop = frog, leafprop = leaf, Genus = variable) %>% 
  mutate(Linear = frogprop - leafprop) %>%
  group_by(population) %>% 
  do(custom_loop(.))

# melt all simulated linear selectivity for plotting null distribution 
Lsel_sim <- L_simulations %>% 
  select(-Linear) %>% 
  melt(id.vars = c("population", "Genus"), measure.vars = paste0("sim",c(1:1000))) %>% 
  group_by(population) %>% 
  arrange(desc(value)) %>%
  mutate(Genus = reorder(Genus, value)) 

# melt observed linear selectivity and assign category based on null distribution
Lsel_obs <- L_simulations %>% 
  melt(id.vars = c("population", "Genus"), measure.vars = c("Linear")) %>% 
  group_by(population) %>% # group by population to rearrange values
  arrange(desc(value)) %>%
  mutate(Genus = reorder(Genus, value)) %>%
  rename(Linear = value) %>% # arrange values by genus
  group_by(population, Genus) %>% 
  mutate(Cat.Linear = case_when(
    Linear > max(Lsel_sim$value) ~ "preferred", # selected are all values falling above null distribution   
    Linear >= min(Lsel_sim$value) & Linear <= max(Lsel_sim$value) ~ "neutral", # neutral are all values falling within null distribution
    Linear < min(Lsel_sim$value) ~ "avoid")) # avoid are all values falling under null distribution

#to arrange genus based on linear selectivity


Lsel_sim <- Lsel_sim %>%
  mutate(Genus = factor(Genus, levels = c("Hypoponera",
                                          "Carebara", "Rogeria",
                                          "Myrmicocrypta", "Anochetus",
                                          "Apterostigma", "Gnamptogenys",
                                          "Hylomyrma", "Octostruma",
                                          "Sericomyrmex",
                                          "Wasmannia","Pheidole", 
                                          "Cyphomyrmex",
                                          "Crematogaster", "Trachymyrmex",
                                          "Strumigenys", "Solenopsis")))

Lsel_obs <- Lsel_obs %>%
  mutate(Genus = factor(Genus, levels = c("Hypoponera",
                                          "Carebara", "Rogeria",
                                          "Myrmicocrypta", "Anochetus",
                                          "Apterostigma", "Gnamptogenys",
                                          "Hylomyrma", "Octostruma",
                                          "Sericomyrmex",
                                          "Wasmannia","Pheidole", 
                                          "Cyphomyrmex",
                                          "Crematogaster", "Trachymyrmex",
                                          "Strumigenys", "Solenopsis")))
```

##### Ant morphology vs frog selectivity

To determine whether the morphological and life history traits of ant species in the leaf litter influences frog selectivity, we performed a Principal Component Analysis (PCA). The PCA aimed at reducing the dimensionality of 17 morphological and life history traits across 17 ant genera found both in the leaf litter and in *O. sylvatica* stomachs. To build the PCA, we used the function **dudi.pca()** from the ‘*ade4*’ package (Dray and Dufour 2007). Scores of components with eigenvalues higher than 1 were selected and used as response variables in a pairwise Wilcoxon test to determine differences between selectivity categories for ants in all diablito populations. P-values were adjusted for multiple testing using the False Discovery Rate control (FDR).

Code

```
# Perform Principal Components Analysis for ant traits 
PCAtraits <- na.omit(ant_traits) %>% 
  filter(Genus %in% Lsel_obs$Genus) %>%  
  select(where(is.numeric)) %>% # retain only numeric columns
  dudi.pca(scale = T, center = T, scannf = FALSE, nf = 3) # we keep the first three components as they explain most of the variance (see scree plot)


# Plot eigenvalues
eigenvalues <- data.frame(eigenvalues = PCAtraits$eig) %>% 
  mutate("Component" = paste0("PC", 1:length(eigenvalues))) %>% 
  filter(Component %in% c("PC1", "PC2", "PC3", "PC4", "PC5")) %>%
  arrange(eigenvalues) %>%
  mutate(Component = reorder(Component, eigenvalues, decreasing = T)) %>%  
  ggplot(aes(x = Component, y = eigenvalues)) +
  geom_bar(stat = "identity", col = "black", fill = "darkgoldenrod", size = 0.3) + 
  geom_hline(yintercept = 1, lty = 2, col = "black") +
  theme_bw() + 
  theme(panel.grid.major = element_blank(),
        panel.grid.minor = element_blank(),
        axis.text.x = NULL, 
        plot.title = element_blank())

eigenvalues
```

Eigenvalues of the first 5 principal components. The first three components have eigenvalues higher than one

### Results

#### Alkaloids differ between species and across diablito frog populations

Skin extracts consisted of 79 alkaloids. The summed amount of alkaloids varied across species and populations (Kruskal-Wallis; X2(5) = 41.542, p < 0.001; Figure 2 A), with *O. sylvatica* having more alkaloids than *H. infraguttatus* (*H. infraguttatus vs. all other O. sylvatica populations*, p < 0.001; Table 1). Within *O. sylvatica*, the Ceiba population had less toxins than all others (p < 0.001; Table 1), while frogs from Santo Domingo had on average the highest alkaloid load (Table 1).

Code

```
#Pairwise Wilcoxon test
pwise.test <- alkadiver %>% 
  select(-latitude, -longitude, -altitude) %>% 
  mutate(total = select(., contains(".")) %>%
           rowSums(na.rm = TRUE)) %>%
  mutate(population = fct_relevel(population, "ceiba", "c_colon", "p_quito", "s_domingo", 
                                  "la_mana", "H_infra")) %>% 
  do(tidy(pairwise.wilcox.test(.$total, .$population,
                               p.adjust.method = "BH")$p.value)) %>% 
  mutate(rowName = c("C. Colón", "P. Quito", "S. Domingo", "La Maná", "H. infraguttatus")) %>% 
  column_to_rownames(var = "rowName") %>% 
  data.frame(.$x) %>% 
  select(-x) %>% 
  rename("C. colón" = c_colon, "Ceiba" = ceiba, "S. Domingo" = s_domingo, 
         "La Maná" = la_mana, "P. Quito" = p_quito)
  

# Results of pairwise comparisons

options(knitr.kable.NA = "-")# to remove "NA's" from the table

T1 <- pwise.test %>% 
  mutate(across(where(is.numeric), round, 3)) %>% 
  mutate(across(where(is.numeric), ~ ifelse(. < 0.001, "<0.001", as.character(.)))) %>% 
  kable(digits = 2, 
      table.attr = 'data-quarto-disable-processing="true"', "html",
      caption = "Table S1:P-values of pairwise Wilcoxon test on differences in summed alkaloids between frog populations") %>% 
  kable_classic(full_width = F, html_font = "Cambria") %>% 
  row_spec(0, italic = T, bold = T) %>% 
  column_spec(1, italic = T, bold = T)

T1
```

Table 1: P-values of pairwise Wilcoxon test on differences in summed alkaloids between frog populations

Table S1:P-values of pairwise Wilcoxon test on differences in summed alkaloids between frog populations

|  | Ceiba | C. colón | P. Quito | S. Domingo | La Maná |
| --- | --- | --- | --- | --- | --- |
| C. Colón | <0.001 | - | - | - | - |
| P. Quito | 0.002 | 0.468 | - | - | - |
| S. Domingo | <0.001 | 0.087 | 0.337 | - | - |
| La Maná | 0.002 | 0.087 | 0.017 | 0.001 | - |
| H. infraguttatus | <0.001 | <0.001 | <0.001 | <0.001 | <0.001 |

Table 1

Figure 2: **Alkaloid profiles differ between *Oophaga sylvatica* populations.** **(A)** The summed relative abundances of 79 alkaloids are visualized as box plots, where each dot represents a frog. The y axis is square-root transformed for visual clarity and summed alkaloids are expressed in arbitrary units (a.u.). Populations are represented with different colors. *Hyloxalus infraguttatus*, collected near the town of La Maná, is shown on the far right. Groups not connected by the same letter are significantly different (Table 1). **(B)** Non-metric multidimensional scaling (NMDS) biplot based on Bray-Curtis dissimilarities show alkaloid profile differences between populations based on the relative abundances of 79 unique alkaloids (Stress = 0.156). Each dot represents a single frog’s alkaloid profile Ellipses represent 95% confidence intervals. Environmental factors (black arrows) and alkaloids (blue arrows) in correlation with the ordination are shown. **(C)** Bubble - heatmap shows percent of all summed alkaloids grouped by alkaloid structural class and population. Histrionicotoxins (HTX); 5,8-disubstituted indolizidines (5,8-I); 3,5-disubstituted indolizidine (3,5-I); Decahydroquinoline (DHQ); 5,6,8-trisubstituted indolizidines (5,6,8-I); Pyrrolidine (Pyr); 3,5-disubstituted pyrrolizidine (3,5-P); Lehmizidine (Lehm); 1,4-disubstituted quinolizidine (1,4-Q); Piperidine (Pip); Pumiliotoxin (PTX); Allopumilliotoxin (aPTX); 4,6-disubstituted quinolizidine (4,6-Q).

Figure 2.png

We next visualized overall alkaloid compositional differences across *O. sylvatica* populations and *H. infraguttatus* using an NMDS (Figure 2 B). The NMDS suggested a two dimensional solution (stress = 0.156) and showed distinct clusters of alkaloid composition. The abundance of all 79 alkaloids varied significantly across groups (PERMANOVA, F(4) = 10.178, p < 0.001). A post-hoc pairwise comparison indicated significant differences between all possible population pairs (Table 2; Figure 2 B), suggesting each group has a unique alkaloid profile. Fitting environmental variables into the NMDS indicated that altitude (r² = 0.72, p = 0.001) and temperature (r² = 0.68, p = 0.001) significantly influenced alkaloid composition across the geographical gradient (Figure 2 B). Given their strong inverse correlation and closely aligned vectors in NMDS space, we report both as reflecting a shared environmental gradient.

Skin alkaloids fell into one of 13 structural families: Histrionicotoxins (HTX); 5,8-disubstituted indolizidines (5,8-I); 3,5-disubstituted indolizidine (3,5-I); Decahydroquinoline (DHQ); 5,6,8-trisubstituted indolizidines (5,6,8-I); Pyrrolidine (Pyr); 3,5-disubstituted pyrrolizidine (3,5-P); Lehmizidine (Lehm); 1,4-disubstituted quinolizidine (1,4-Q); Piperidine (Pip); Pumiliotoxin (PTX); Allopumilliotoxin (aPTX); 4,6-disubstituted quinolizidine (4,6-Q). When examined more closely, histrionicotoxins, decahydroquinolines and indolizidines were the most abundant alkaloid classes relative to the total alkaloid content in both H. infraguttatus and all O. sylvatica populations sampled (Figure 2 C). Several indolizidines, including 5,6,8-I (r² = 0.54, p = 0.001), 5,8-I (r² = 0.54, p = 0.001) & 3,5-I (r² = 0.46, p = 0.001) contributed significantly to alkaloid ordination (Figure 2 B).

Code

```
colnames(pair.mod) <- c("Contrasts", "df", "Sums of sq.", "F", 
                        "R2","p-value", "p.adjusted", "Sig.")

T2 <- pair.mod %>%
  select(Contrasts, df, F, R2, p.adjusted) %>% 
  mutate(Contrasts = c("La Maná vs. Ceiba", "La Maná vs. C. Colón", 
                   "La Maná vs. S. Domingo", "La Maná vs.P. Quito",
                   "La Maná vs. H. infraguttatus", "Ceiba vs. C. Colón", 
                   "Ceiba vs. S. Domingo", "Ceiba vs. P. Quito", 
                   "Ceiba vs. H. infraguttatus", "C. Colón vs. S. Domingo", 
                   "C. Colón vs. P. Quito", "C. Colón vs. H.infraguttatus",
                   "S. Domingo vs. P. Quito", "S. Domingo vs. H. infraguttatus",
                   "P. Quito vs. H. infraguttatus")) %>% 
  kable(digits = 3, 
      table.attr = 'data-quarto-disable-processing="true"', "html",
      caption = "Table S2: Summary of the results of pairwise comparisons of a PERMANOVA in alkaloid composition between frog populations") %>% 
  kable_classic(full_width = F, html_font = "Cambria") %>% 
  row_spec(0, bold = T) 

T2
```

Table 2: Summary of the results of pairwise comparisons of a PERMANOVA in alkaloid composition between *O. sylvatica* populations

Table S2: Summary of the results of pairwise comparisons of a PERMANOVA in alkaloid composition between frog populations

| Contrasts | df | F | R2 | p.adjusted |
| --- | --- | --- | --- | --- |
| La Maná vs. Ceiba | 1 | 8.187 | 0.325 | 0.001 |
| La Maná vs. C. Colón | 1 | 8.028 | 0.308 | 0.001 |
| La Maná vs. S. Domingo | 1 | 8.186 | 0.313 | 0.001 |
| La Maná vs.P. Quito | 1 | 8.129 | 0.300 | 0.001 |
| La Maná vs. H. infraguttatus | 1 | 16.943 | 0.499 | 0.001 |
| Ceiba vs. C. Colón | 1 | 9.297 | 0.354 | 0.002 |
| Ceiba vs. S. Domingo | 1 | 16.592 | 0.494 | 0.001 |
| Ceiba vs. P. Quito | 1 | 13.472 | 0.428 | 0.001 |
| Ceiba vs. H. infraguttatus | 1 | 13.724 | 0.462 | 0.001 |
| C. Colón vs. S. Domingo | 1 | 10.972 | 0.379 | 0.001 |
| C. Colón vs. P. Quito | 1 | 8.581 | 0.311 | 0.001 |
| C. Colón vs. H.infraguttatus | 1 | 17.336 | 0.505 | 0.001 |
| S. Domingo vs. P. Quito | 1 | 8.024 | 0.297 | 0.001 |
| S. Domingo vs. H. infraguttatus | 1 | 25.880 | 0.604 | 0.001 |
| P. Quito vs. H. infraguttatus | 1 | 20.150 | 0.528 | 0.001 |

Table 2

Code

```
#Boxplots 
f2a <- alkadiver %>% 
  select(-latitude, -longitude, -altitude) %>% 
  mutate(total = select(., contains(".")) %>%
           rowSums(na.rm = TRUE)) %>% 
  mutate(population = factor(population, 
                levels = c("ceiba", "c_colon", "p_quito",
                           "s_domingo", "la_mana", "ala_manaHI"))) %>% 
  ggplot(aes(y=total/10000000, x=population, fill=population)) + 
  geom_boxplot(outlier.shape = NA) + 
  geom_jitter(position=position_jitterdodge(0.1), shape = 21, 
              size = 4, alpha=0.6) +
  theme_classic(20) + 
  ylab("summed alkaloids (a.u.)") +
  scale_fill_manual(values=c("#d16b54", "#e8c95d", "#433447", "#b9b09f",
                             "#a9d8c8", "#ffffff"),
                    name = "populations", 
                    labels = c("Ceiba", "C. colon", "P. Quito", 
                               "S. Domingo", "La Maná", "H. infragutatus")) + 
  scale_y_sqrt() + 
  scale_y_continuous(trans = 'sqrt') +
  theme(axis.title.x = element_blank(), 
        axis.text.x = element_blank(), axis.title = element_text(size=15),
        plot.title = element_text(size=28,hjust=0.5),
        legend.position = "none")


# To make the NMDS plot in ggplot, we extract the dimension scores from the NMDS and add them to the main data
NMDSdims <- alkadiver %>% 
  mutate(NMDS1 = scores(ord2)$sites[,1], NMDS2 = scores(ord2)$sites[,2]) %>%
  data.frame()

# to extract the coordinates for the vectors of the environmental variables
en_coord <-  as.data.frame(scores(en, "vectors"))

en_alk_coord <-  as.data.frame(scores(en.alka, "vectors"))[c(1,4,6),]

# The contribution of every environmental variable
env.alka <- en$vectors$arrows %>%
  data.frame() %>%
  mutate(r2 = en$vectors$r) %>% 
  mutate("p-value" = en$vectors$pvals) %>% 
  kable(digits = 3, 
      table.attr = 'data-quarto-disable-processing="true"', "html") %>% 
  kable_classic(full_width = F, html_font = "Cambria") %>% 
  row_spec(0, bold = T)


#and the plot
f2b <- ggplot(NMDSdims, aes(x = NMDS1, y = NMDS2)) + 
  stat_ellipse(aes(colour = population, fill = population), 
               geom = "polygon", level = 0.95, alpha = 0.2, type = "t") +
  geom_point(aes(fill = population), pch = 21, size = 4, alpha = 0.5) + 
  scale_colour_manual(values = c("#e8c95d", "#d16b54", "gray90", "#a9d8c8", "#433447", "#b9b09f")) + 
  scale_fill_manual(values = c("#e8c95d", "#d16b54", "gray90", "#a9d8c8", "#433447", "#b9b09f")) + 
  geom_segment(aes(x = 0, y = 0, xend = NMDS1, yend = NMDS2), 
               data = en_coord, linewidth = 1, alpha = 0.5, colour = "grey30",
               arrow = arrow(type = "open", length = unit(0.1, "inches"))) + 
  geom_text(data = en_coord, aes(x = NMDS1-0.1, y = NMDS2), colour = "grey30", 
            fontface = "bold", label = row.names(en_coord)) +
  geom_segment(aes(x = 0, y = 0, xend = NMDS1, yend = NMDS2), 
               data = en_alk_coord, linewidth = 1, alpha = 0.5, colour = "red",
               arrow = arrow(type = "open", length = unit(0.1, "inches"))) + 
  geom_text(data = en_alk_coord, aes(x = NMDS1-0.1, y = NMDS2), colour = "red", 
            fontface = "bold", label = row.names(en_alk_coord)) +
  theme_bw() +
  theme(panel.grid.major = element_blank(), panel.grid.minor = element_blank(),
        axis.text.x = NULL, axis.text.y = NULL, legend.position = "none")


#Heatmap to visualize percentage of summed alkaloids grouped by alkaloid class and population

# To create a palette for a gradient color
pal <- wes_palette("Zissou1", 100, type = "continuous")

#By structural family
f2c <- alkadiver %>% 
  select(-latitude, -longitude, -altitude) %>% 
  #mutate(total = select(., contains(".")) %>%
  #         rowSums(na.rm = TRUE)) %>%
  mutate(population = fct_relevel(population, "ceiba", "c_colon", "p_quito", "s_domingo", 
                                  "la_mana", "H_infra")) %>% 
  pivot_longer(cols = c(indo.137:pyrrz.526), 
               names_to = "alkaloid", values_to = "abundance") %>% 
  bind_cols(feat_tab %>% 
              select(family, santos2016_source) %>%
  slice(rep(1:n(), times = 59))) %>% 
  group_by(family, population) %>% 
  summarise(sum.str.alk = sum(abundance)) %>% 
  group_by(population) %>% 
  mutate(percent = (sum.str.alk / sum(sum.str.alk))*100) %>% 
  mutate(family = factor(family, levels = c('HTX', '5,8-I', '3,5-I', 'DHQ', 
                                            '5,6,8-I', 'Pyr', '3,5-P', 'Lehm',
                                            '1,4-Q', 'Pip', 'PTX', '4,6-Q', 
                                            'aPTX'))) %>% 
  mutate(population = factor(population, levels = c("H_infra", "la_mana", "s_domingo", 
                                                    "p_quito", "c_colon", 
                                                    "ceiba"))) %>% 
  ggplot(aes(x = population, y = family, colour = percent)) + 
  geom_point(aes(size = percent, colour = percent)) +
  scale_color_gradientn(colours = pal) +
  theme_bw() + coord_flip() +
  theme(axis.text.x = element_text(angle = 45, hjust = 1, vjust = 1),
        legend.text.position = "right")


#By superfamily
#f2c <- alkadiver %>%  # to get the sum of alkaloids per individual
#  select(-altitude) %>%
#  melt() %>% 
#  mutate(toxin = case_when(grepl("indo", variable) ~ "indolizidine", 
 #                          grepl("allo", variable) ~ "allopumiliotoxin",
#                           grepl("quino", variable) ~ "quinolizidine", 
#                           grepl("pyrrz", variable) ~ "pyrrolizidine",
#                           grepl("deca", variable) ~ "decahydroquinoline", 
#                           grepl("hist", variable) ~ "histrionicotoxin",
#                           grepl("lehm", variable) ~ "lehmizidine", 
#                           grepl("piper", variable) ~ "piperidine", 
#                           grepl("pumi", variable) ~ "pumiliotoxin", 
#                           grepl("pyrro", variable) ~ "pyrrolidine")) %>%
#  select(-variable) %>% 
#  group_by(toxin, population) %>% 
#  summarise(sum.alk = sum(value)) %>% 
#  group_by(population) %>% 
#  mutate(percent = (sum.alk / sum(sum.alk))*100) %>% 
#  mutate(toxin = factor(toxin, levels = c('histrionicotoxin', 'indolizidine',
#                                          'decahydroquinoline', 'pumiliotoxin',
#                                          'pyrrolidine', 'pyrrolizidine', 
#                                          'quinolizidine', 'piperidine', 
#                                          'allopumiliotoxin', 'lehmizidine'))) %>% 
#  mutate(population = factor(population, levels = c("H_infra", "la_mana", "s_domingo", 
#                                                    "p_quito", "c_colon", 
#                                                    "ceiba"))) %>% 
#  ggplot(aes(x = population, y = toxin, colour = percent)) + 
#  geom_point(aes(size = percent, colour = percent)) +
#  scale_color_gradientn(colours = pal) +
#  theme_bw() + coord_flip() +
#  theme(axis.text.x = element_text(angle = 45, hjust = 1, vjust = 1))
```

#### Defended frogs consumed more ants relative to other prey types and compared to the diet of the undefended species

We found that the number of prey consumed in different categories differ significantly across populations and between species (GLMM, population x prey type: X2(20) = 83.59, p < 0.001; Figure 3 A & B). Post hoc pairwise comparisons and the species selectivity index (*ssi*) showed that all *O. sylvatica* populations consumed significantly more ants than other prey categories (x̅ = 75%; emmeans (ants vs. all prey): p-value = <0.001; ssirange= 0.66 - 0.85; Figure 3 B, Table 3), whereas *H. infraguttatus* showed a more generalist dietary pattern, consuming a smaller but diverse array of arthropods including ants (45%), beetles (14%) and ‘other’ arthropods (25%; emmeans (all prey comparisons): p-value = >0.05; ssi = 0.27; Figure 3 B, Table 3). It is worth noting that only one *H. infraguttatus* had 18 ants in the stomach, which accounts for nearly half of the total consumed for this species in our data set. When removing this individual, ants made up 36% of the total diet, followed by ‘other’ arthropods (29.6%) and beetles (16.5%).

Code

```
# to visualize the bipartite interaction matrix:

# First, I formatted the diet data to make a sankey plot to visualize connections between preys and populations  
links <- data.frame(sum_diet) %>% 
  rownames_to_column(var = "target") %>% 
  pivot_longer(cols = c(ants, beetle, larvae, mites, other), names_to = "source", values_to = "value") %>% 
  mutate(target = case_when(grepl("ceiba", target) ~ 5, grepl("c.colon", target) ~ 9, 
                            grepl("h.infragutatus", target) ~ 10, grepl("mana", target) ~ 6, 
                            grepl("p.quito", target) ~ 7, grepl("s.domingo", target) ~ 8)) %>% 
  mutate(source = case_when(grepl("ants", source) ~ 0, grepl("beetle", source) ~ 2, 
                            grepl("larvae", source) ~ 3, grepl("mites", source) ~ 1, 
                            grepl("other", source) ~ 4)) %>% 
  data.frame() %>% 
  round() %>% 
  mutate(perc = value+1) %>% #add1 so the 0 are not 0
  arrange(source) %>% 
  select(source, target, value, perc)

#Here I repeated the number of rows according to the number of prey items per category per population
links_rep <- links[rep(row.names(links), times = links$perc), ]

# the Sankey plot
fig3a <- links_rep %>% 
  select(source, target) %>% 
  make_long(source, target) %>% 
  arrange(desc(node)) %>% 
  ggplot(aes(x = x, 
               next_x = next_x, 
               node = node, 
               next_node = next_node,
               fill = factor(node))) +
  geom_sankey(alpha=0.4, col = "black", lwd = 0.03) +
  theme_sankey(base_size = 16) +
  scale_fill_manual(values= c("#3E63A6", "#6AB897", "#A4BB7A","#E5C049", "#FDE725FF", 
                              rep("black",6))) + 
  theme(legend.position = "none")

##Alternative Chord diagram

sumdiet2 <- t(sum_diet)[ c(3,5,2,4,1),c(1,5,6,2,4,3)] 

chord.plot <- chordDiagram(sumdiet2, grid.col = c("#E5C049", "#FDE725FF", "#A4BB7A", "#6AB897", "#3E63A6",    
                                    rep("black",6)))
```


Code

```
# Plot of emmeans with 95% CIs 
fig3b <- ggplot(emm1, aes(prey, emmean, color = prey))  +
  geom_hline(yintercept = 0, linetype = 3, col = "grey") +
  geom_linerange(aes(ymin=asymp.LCL, ymax=asymp.UCL), linewidth=4,show.legend = F, alpha = 0.6) +
  geom_point(col = "black", size = 1) +
  facet_grid(.~population, scales = "free") +
  theme_bw() +
  theme(panel.grid.major = element_blank(),
                                    panel.grid.minor = element_blank(),
                                    axis.text.x = NULL, legend.position = "none") + 
  scale_color_manual(values=c("#3E63A6","#6AB897", "#A4BB7A", "#E5C049",  "#FDE725FF"))
```

Code

```
t3 <- data.frame(emmeans(sd1, list(pairwise ~ population*prey), 
                           adjust = "tukey")$`pairwise differences of population, prey`)[c(6,12,18,24,35,41,47,53,90,96,102,108,116,122,128,134,141,147,153,159,63,69,75,81),]# Manually choose the contrasts per population

T3 <- t3 %>%
  rename(Contrasts = X1) %>% 
  select(Contrasts, estimate, SE, p.value) %>% 
  remove_rownames() %>% 
  mutate(p.value = c(rep("<0.001",20), rep(">0.05", 4))) %>% #to read easily p.values
  mutate(Contrasts = rep(c("Ants vs. Beetles", "Ants vs. Larvae", 
                   "Ants vs. Mites", "Ants vs. Other"), 6)) %>% 
  kable(digits = 3, 
      table.attr = 'data-quarto-disable-processing="true"', "html",
      caption = "Table S3: Summarized results of estimated marginal means between frog species and prey items. P-values were adjusted using Tukey’s method") %>% 
  kable_classic(full_width = F, html_font = "Cambria") %>% 
  row_spec(0, bold = T) %>% 
  pack_rows("Ceiba",1,4) %>% 
  pack_rows("C. Colón",5,8) %>%
  pack_rows("La Maná",9,12) %>%
  pack_rows("P. Quito",13,16) %>% 
  pack_rows("S. Domingo",17,20) %>%
  pack_rows("H. infraguttatus",21,24)

T3
```

Table 3: Summarized results of estimated marginal means between frog species and prey items. P-values were adjusted using Tukey’s method

Table S3: Summarized results of estimated marginal means between frog species and prey items. P-values were adjusted using Tukey’s method

| Contrasts | estimate | SE | p.value |
| --- | --- | --- | --- |
| **Ceiba** | | | |
| Ants vs. Beetles | 3.729 | 0.515 | <0.001 |
| Ants vs. Larvae | 4.489 | 0.720 | <0.001 |
| Ants vs. Mites | 2.027 | 0.312 | <0.001 |
| Ants vs. Other | 4.502 | 0.720 | <0.001 |
| **C. Colón** | | | |
| Ants vs. Beetles | 2.415 | 0.493 | <0.001 |
| Ants vs. Larvae | 2.401 | 0.493 | <0.001 |
| Ants vs. Mites | 1.226 | 0.366 | <0.001 |
| Ants vs. Other | 2.201 | 0.457 | <0.001 |
| **La Maná** | | | |
| Ants vs. Beetles | 2.991 | 0.306 | <0.001 |
| Ants vs. Larvae | 4.001 | 0.461 | <0.001 |
| Ants vs. Mites | 1.485 | 0.204 | <0.001 |
| Ants vs. Other | 3.432 | 0.368 | <0.001 |
| **P. Quito** | | | |
| Ants vs. Beetles | 3.094 | 0.435 | <0.001 |
| Ants vs. Larvae | 4.914 | 1.012 | <0.001 |
| Ants vs. Mites | 1.783 | 0.308 | <0.001 |
| Ants vs. Other | 21.165 | 3378.581 | <0.001 |
| **S. Domingo** | | | |
| Ants vs. Beetles | 2.574 | 0.448 | <0.001 |
| Ants vs. Larvae | 2.926 | 0.530 | <0.001 |
| Ants vs. Mites | 1.795 | 0.368 | <0.001 |
| Ants vs. Other | 4.414 | 1.018 | <0.001 |
| **H. infraguttatus** | | | |
| Ants vs. Beetles | 0.552 | 0.488 | >0.05 |
| Ants vs. Larvae | 1.460 | 0.656 | >0.05 |
| Ants vs. Mites | 0.868 | 0.518 | >0.05 |
| Ants vs. Other | 0.311 | 0.477 | >0.05 |

Table 3

##Frogs’ alkaloid composition suggests a predominantly ant-based diet

We found that across all frog populations, including the “non-toxic” species H. infraguttatus, ant-derived alkaloids appeared to represent the most abundant component of their chemical profile, while a smaller but important contribution was attributed to mite-derived alkaloids (Figure 3 C). Tukey’s post hoc test revealed that the proportion of ant-derived alkaloids was significantly higher than mite-derived alkaloids (p<0.001). Neither total alkaloid abundance nor abundance within structural families showed a significant correlation with the number of ants or mites consumed across populations (All p-values > 0.05; Table 4).

Code

```
#Visualization
#Make subset matrices for every population
allpop.chord <- alkadiver %>% 
  select(-latitude, -longitude, -altitude) %>% 
  mutate(total = select(., contains(".")) %>%
           rowSums(na.rm = TRUE)) %>%
  mutate(population = fct_relevel(population, "ceiba", "c_colon", "p_quito", "s_domingo", 
                                  "la_mana", "H_infra")) %>% 
  pivot_longer(cols = c(indo.137:pyrrz.526), 
               names_to = "alkaloid", values_to = "abundance") %>% 
  bind_cols(feat_tab %>% 
              select(family, santos2016_source) %>%
  slice(rep(1:n(), times = 59))) %>% 
  group_by(population, family, santos2016_source) %>% 
  summarize(abundance = sum(abundance)) %>% 
  mutate(family = factor(family, levels = c('HTX', 'DHQ', '5,8-I', 'Pyr', 
                                            '3,5-P', 'Lehm', 'Pip', '3,5-I',  
                                            '5,6,8-I', '1,4-Q', 'PTX', 
                                            '4,6-Q', 'aPTX'))) %>% 
  #count(population, family, saporito2017_source) %>%
  pivot_wider(names_from = family, values_from = abundance, values_fill = 0) %>%
  ungroup() %>% 
  group_by(population) %>%
  group_split() %>%
  set_names(map_chr(., ~ unique(.x$population))) %>%
  map(~ .x %>%
        select(-population) %>%
        column_to_rownames("santos2016_source") %>%
        as.matrix()
      )

# Assign a color to each sector, using a named vector
colors <- c("#6AB897", "sienna3", "#3E63A6", rep("black",13))


## Ceiba
ceiba =allpop.chord$`1`

# Check the sector names (rows and columns of the matrix)
sec.ceiba <- union(rownames(ceiba), colnames(ceiba))


# Set names
names(colors) <- sec.ceiba 

ceiba <- ceiba[c("Both", "Mites", "Ants"), c("HTX", "DHQ", "Lehm", "3,5-P", 
                                             "Pip", "5,6,8-I", "1,4-Q", "5,8-I", 
                                             "Pyr", "3,5-I", "PTX", "4,6-Q", 
                                             "aPTX")]

# Run the chord diagram
ch_cei <- chordDiagram(ceiba, grid.col = colors)
```


Code

```
## C. Colón
C.colon =allpop.chord$`2`

# Check the sector names (rows and columns of the matrix)
sec.colon <- union(rownames(C.colon), colnames(C.colon))

# Set names
names(colors) <- sec.colon

C.colon <- C.colon[c("Both", "Mites", "Ants"), c("HTX", "DHQ", "Lehm", "3,5-P", 
                                             "Pip", "5,6,8-I", "1,4-Q", "5,8-I", 
                                             "Pyr", "3,5-I", "PTX", "4,6-Q", 
                                             "aPTX")]

# Run the chord diagram
ch_col <- chordDiagram(C.colon, grid.col = colors)
```


Code

```
## P. Quito
P.quito =allpop.chord$`3`

# Check the sector names (rows and columns of the matrix)
sec.quito <- union(rownames(P.quito), colnames(P.quito))
# Set names
names(colors) <- sec.quito

P.quito <- P.quito[c("Both", "Mites", "Ants"), c("HTX", "DHQ", "Lehm", "3,5-P", 
                                             "Pip", "5,6,8-I", "1,4-Q", "5,8-I", 
                                             "Pyr", "3,5-I", "PTX", "4,6-Q", 
                                             "aPTX")]
# Run the chord diagram
ch_qui <- chordDiagram(P.quito, grid.col = colors)
```


Code

```
## S. Domingo
S.domingo =allpop.chord$`4`

# Check the sector names (rows and columns of the matrix)
sec.domingo <- union(rownames(S.domingo), colnames(S.domingo))
# Set names
names(colors) <- sec.domingo

S.domingo <- S.domingo[c("Both", "Mites", "Ants"), c("HTX", "DHQ", "Lehm", "3,5-P", 
                                             "Pip", "5,6,8-I", "1,4-Q", "5,8-I", 
                                             "Pyr", "3,5-I", "PTX", "4,6-Q", 
                                             "aPTX")]
# Run the chord diagram
ch_dom <- chordDiagram(S.domingo, grid.col = colors)
```


Code

```
## Maná
Mana =allpop.chord$`5`

# Check the sector names (rows and columns of the matrix)
sec.mana <- union(rownames(Mana), colnames(Mana))
# Set names
names(colors) <- sec.mana

Mana <- Mana[c("Both", "Mites", "Ants"), c("HTX", "DHQ", "Lehm", "3,5-P", 
                                             "Pip", "5,6,8-I", "1,4-Q", "5,8-I", 
                                             "Pyr", "3,5-I", "PTX", "4,6-Q", 
                                             "aPTX")]
# Run the chord diagram
ch_man <- chordDiagram(Mana, grid.col = colors)
```


Code

```
## H. infra
H.infra =allpop.chord$`6`

# Check the sector names (rows and columns of the matrix)
sec.infra <- union(rownames(H.infra), colnames(H.infra))
# Set names
names(colors) <- sec.infra

H.infra <- H.infra[c("Both", "Mites", "Ants"), c("HTX", "DHQ", "Lehm", "3,5-P", 
                                             "Pip", "5,6,8-I", "1,4-Q", "5,8-I", 
                                             "Pyr", "3,5-I", "PTX", "4,6-Q", 
                                             "aPTX")]
# Run the chord diagram
ch_inf <- chordDiagram(H.infra, grid.col = colors)
```

Figure 3: **Diet differs between frog species and across diablito frog populations.** **(A)** Chord diagram showing a bipartite network of the dietary interactions between two frog species, the non-toxic *H. infraguttatus* and the toxic *O. sylvatica*, and their arthropod prey in five localities. The thickness of the connecting bars represents the average number of prey items consumed, indicating its relative importance in the network. Species specificity index (ssi) is shown next to each frog species, ranging from 0 (generalist) to 1 (specialist). **(B)** Pairwise comparison of prey item consumption between frog species. Dots represent the mean and bars represent 95% confidence intervals for the estimated marginal means. Non-overlapping bars indicate statistically significant differences. **(C)** Chord diagrams showing the most abundant alkaloid families and their putative arthropod source, across frog populations (sensu Santos et al. 2016). The thickness of the connecting bars represents the summed alkaloid abundance per structural class. Histrionicotoxins (HTX); 5,8-disubstituted indolizidines (5,8-I); 3,5-disubstituted indolizidine (3,5-I); Decahydroquinoline (DHQ); 5,6,8-trisubstituted indolizidines (5,6,8-I); Pyrrolidine (Pyr); 3,5-disubstituted pyrrolizidine (3,5-P); Lehmizidine (Lehm); 1,4-disubstituted quinolizidine (1,4-Q); Piperidine (Pip); Pumiliotoxin (PTX); Allopumilliotoxin (aPTX); 4,6-disubstituted quinolizidine (4,6-Q).

Figure 3.png

Code

```
T4 <- diet %>% 
  filter(prey %in% c("ants", "mites")) %>% 
  pivot_wider(names_from = "prey", values_from = "count", 
              id_cols = c(frog_id, population)) %>% 
  left_join(alkadiver %>% 
              select(-latitude, -longitude, -altitude) %>% 
              mutate(total = select(., contains(".")) %>%
                       rowSums(na.rm = TRUE)), by = "frog_id") %>% 
  na.omit() %>% 
  select(ants, mites, c(indo.137: total)) %>% 
  setNames(c(names(.)[1:2], feat_tab %>% 
               select(family) %>% 
               pull(), "total")) %>%
  {split.default(., names(.))} %>%                    # group columns by name
  map_dfc(~ rowMeans(as.data.frame(.), na.rm = TRUE)) %>% 
  corr.test() %>% 
  .$ci %>%
  rownames_to_column("correlations") %>%
  filter(str_detect(correlations, "mite|ant")) %>% 
  kable(digits = 3, 
      table.attr = 'data-quarto-disable-processing="true"', "html") %>% 
  kable_classic(full_width = F, html_font = "Cambria") %>% 
  row_spec(0, bold = T)

T4
```

Table 4: Summary of the results of pairwise correlations between each alkaloid class and ant & mite abundance.

| correlations | lower | r | upper | p |
| --- | --- | --- | --- | --- |
| 1,4-Q-ants | -0.165 | 0.095 | 0.342 | 0.475 |
| 1,4-Q-mites | -0.270 | -0.015 | 0.242 | 0.911 |
| 3,5-I-ants | -0.318 | -0.067 | 0.192 | 0.612 |
| 3,5-I-mites | -0.382 | -0.140 | 0.120 | 0.290 |
| 3,5-P-ants | -0.234 | 0.024 | 0.278 | 0.858 |
| 3,5-P-mites | -0.220 | 0.038 | 0.292 | 0.773 |
| 4,6-Q-ants | -0.056 | 0.203 | 0.436 | 0.123 |
| 4,6-Q-mites | -0.278 | -0.023 | 0.234 | 0.860 |
| 5,6,8-ants | -0.199 | 0.060 | 0.312 | 0.650 |
| 5,6,8-mites | -0.348 | -0.101 | 0.159 | 0.446 |
| 5,8-I-ants | -0.207 | 0.051 | 0.304 | 0.699 |
| 5,8-I-mites | -0.304 | -0.052 | 0.207 | 0.697 |
| ants-aPTX | -0.314 | -0.063 | 0.196 | 0.636 |
| ants-DHQ | -0.464 | -0.236 | 0.021 | 0.072 |
| ants-HTX | -0.326 | -0.076 | 0.183 | 0.565 |
| ants-Lehm | -0.411 | -0.173 | 0.087 | 0.191 |
| ants-mites | 0.093 | 0.341 | 0.549 | 0.008 |
| ants-Pip | -0.330 | -0.081 | 0.179 | 0.543 |
| ants-PTX | -0.422 | -0.186 | 0.074 | 0.158 |
| ants-Pyr | -0.296 | -0.044 | 0.215 | 0.743 |
| ants-total | -0.319 | -0.068 | 0.191 | 0.608 |
| aPTX-mites | -0.297 | -0.044 | 0.215 | 0.741 |
| DHQ-mites | -0.403 | -0.164 | 0.097 | 0.216 |
| HTX-mites | -0.386 | -0.144 | 0.116 | 0.276 |
| Lehm-mites | -0.329 | -0.080 | 0.180 | 0.549 |
| mites-Pip | -0.301 | -0.048 | 0.210 | 0.716 |
| mites-PTX | -0.230 | 0.027 | 0.281 | 0.838 |
| mites-Pyr | -0.364 | -0.119 | 0.141 | 0.368 |
| mites-total | -0.394 | -0.153 | 0.107 | 0.248 |

Table 4

#### Alkaloid diversity among sites is associated with variation in leaf litter ant communities

As the diet of *O. sylvatica* is mainly composed of ants, we looked at compositional differences in leaf litter ant communities across the geographical gradient. A total of 46 ant genera were recovered from Winkler traps in leaf litter communities. The NMDS suggested a two-dimensional solution (stress = 0.130), representing partially overlapping clusters of ant communities across localities in the ordination space (Figure 4). PERMANOVA and post-hoc comparisons suggested significant differences in the composition of ant communities between all pairs of localities (PERMANOVA, F(3) = 5.0525, p < 0.001; Table 5, Figure 4 A). Precipitation (r² = 0.41, p = 0.001), altitude (r² = 0.458, p = 0.001) and temperature (r² = 0.49, p = 0.001) significantly influenced ant availability across the geographical gradient (Figure 4 B). A subset of ant genus, including *Octostruma* (r² = 0.59, p = 0.001), *Pachycondyla* (r² = 0.54, p = 0.001), *Rogeria* (r² = 0.51, p = 0.001) and *Solenopsis* (r² = 0.48, p = 0.001) significantly contributed to the ordination. We next asked whether availability of leaf litter ant communities between locations is associated with skin alkaloid diversity. The procrustes analysis suggested that variation in alkaloid diversity among sites is significantly correlated with variation in availability of ant communities (r = 0.47, p-value = 0.001, 999 permutations; Figure 4 B).

Code

```
# To make the plot in ggplot, we extract the dimension scores from the NMDS and add them to the main data
ants.nmds <- ant_co %>%
  mutate(NMDS1 = scores(ord.co2)$sites[,1],
         NMDS2 = scores(ord.co2)$sites[,2]) %>% 
  data.frame()

# to extract the coordinates for the vectors of the environmental variables
en_coord.ant <-  as.data.frame(scores(en.co, "vectors"))

en_coord.ant2 <- as.data.frame(scores(en.co.ants, "vectors"))

#and the plot
fig4a <- ggplot(ants.nmds, aes(x = NMDS1, y = NMDS2*-1)) + 
  #stat_ellipse(aes(colour = SITE, fill = SITE), 
  #             geom = "polygon", level = 0.95, alpha = 0.1, type = "t") +
  geom_point(aes(fill = SITE), 
             pch = 24, col = "black", size = 2, alpha = 0.7) + 
  scale_colour_manual(values = c("#a9d8c8", "#e8c95d", "#b9b09f", "#433447")) + 
  scale_fill_manual(values = c("#a9d8c8", "#e8c95d", "#b9b09f", "#433447")) + 
  geom_segment(aes(x = 0, y = 0, xend = NMDS1, yend = NMDS2*-1), 
               data = en_coord.ant, linewidth = 1, alpha = 0.5, colour = "grey30",
               arrow = arrow(type = "open", length = unit(0.1, "inches"))) + 
  geom_text(data = en_coord.ant, aes(x = NMDS1-0.1, y = NMDS2*-1), colour = "grey30", 
            fontface = "bold", label = row.names(en_coord.ant)) + 
  geom_segment(aes(x = 0, y = 0, xend = NMDS1, yend = NMDS2*-1), 
               data = en_coord.ant2[c(13,7,2,5),], linewidth = 1, 
               alpha = 0.5, colour = "red",
               arrow = arrow(type = "open", length = unit(0.1, "inches"))) + 
  geom_text(data = en_coord.ant2[c(13,7,2,5),], aes(x = NMDS1-0.1, y = NMDS2*-1), 
            colour = "red", 
            fontface = "bold", label = row.names(en_coord.ant2[c(13,7,2,5),])) +
  theme_bw() +
  theme(panel.grid.major = element_blank(), panel.grid.minor = element_blank(),
        axis.text.x = NULL, axis.text.y = NULL, legend.position = "none")

###-----------------------###
###  Procrustes analysis  ###
###-----------------------###

#NMDS of alkaloids excluding ceiba and infragutatus
ord.alk <- metaMDS(alkadiver %>% 
                      filter(!(population=="ceiba")) %>%
                      filter(!(population=="H_infra")) %>%
                      select(where(is.numeric), -altitude,
                             -latitude,-longitude), 
  perm=9999, distance = "bray", 
               k = 2, autotransform = FALSE)

#the ordination was ran a second time starting at the previous best solution to ensure the stability and reliability of the results in the NMDS 
ord.alk2<-metaMDS(alkadiver %>% 
                      filter(!(population=="ceiba")) %>% 
                      filter(!(population=="H_infra")) %>%
                      select(where(is.numeric), -altitude,
                             -latitude,-longitude), 
                  previous.best = TRUE)

## NMDS of ants removing an outlier
set.seed(123)

ant_co2 <- ant_co %>% #remove one sample from P. quito that is an outlier in the ordination (only 1 wasmannia)
  group_by(SITE) %>%
  filter(!(SITE=="Puerto Quito")) %>% 
  slice_sample(n = 10) %>% 
  bind_rows(ant_co %>% 
              filter(SITE=="Puerto Quito")) %>% 
  mutate(SITE = factor(SITE)) %>% 
  ungroup() %>% 
  mutate(SITE = fct_relevel(SITE, "Mana", "Colon", "Santo Domingo", 
                            "Puerto Quito")) %>%
  arrange(as.integer(SITE))

#the ordination was performed using Bray-Curtis dissimilarities
ord.co3 <- metaMDS(ant_co2 %>%
                     select(where(is.numeric), -altitude, 
                            -latitude, -longitude), 
                  perm=9999, distance = "bray",
                  k = 2, autotransform = FALSE)

proc <- procrustes(ord.co3,ord.alk2, scale = T)


procru.test <- protest(ord.co3,ord.alk2, permutations = 999)


Yrot <- data.frame(proc$Yrot) %>% 
  bind_cols(alkadiver %>% 
                      filter(!(population=="ceiba")) %>% 
                      filter(!(population=="H_infra")) %>%
              select(population)) %>% 
  bind_cols(data.frame(proc$X))

procu.plot <- ggplot(Yrot, aes(x = NMDS1, y = NMDS2, colour = population)) +
  geom_hline(yintercept = 0, linetype = "dashed", color = "gray", alpha = 0.5) +
  geom_vline(xintercept = 0, linetype = "dashed", color = "gray", alpha = 0.5) +
  geom_segment(aes(x = X1, y = X2, 
                   xend = NMDS1, yend = NMDS2,
                   colour = population), 
               alpha = 0.2) + 
  geom_point(aes(fill = population), pch = 24, size = 1.3) +
  geom_point(aes(X1, X2, fill = population), 
             pch = 21, col = "black", size = 3, alpha = 0.7) +
  theme_bw() +
  theme(panel.grid.major = element_blank(), panel.grid.minor = element_blank(),
        axis.text.x = NULL, axis.text.y = NULL, legend.position = "none") +
  scale_fill_manual(values=c("#e8c95d", "#a9d8c8", "#b9b09f", "#433447")) +
  scale_color_manual(values=c("#e8c95d", "#a9d8c8", "#b9b09f", "#433447"))

fig4c <- ant_all %>%
  filter(TRAP == "Winkler") %>%
  select(SITE, GENERA, X.) %>% 
  group_by(SITE, GENERA) %>% 
  mutate(total = sum(X.)) %>% 
  select(-X.) %>% 
  distinct() %>% 
  filter(total>10) %>% 
  mutate(GENERA = factor(GENERA, levels = c("Solenopsis", "Wasmannia", "Pheidole", 
                                        "Nylanderia", "Carebara", "Rogeria", "Strumigenys", "Hypoponera",
                                        "Octostruma", "Apterostigma", "Cyphomyrmex", "Prionopelta",
                                        "Stenamma", "Anochetus"))) %>% 
  mutate(SITE = factor(SITE, levels = c("Mana", "Santo Domingo","Puerto Quito", 
                                        "Colon"))) %>% 
  ggplot(aes(x = GENERA, y = SITE, colour = total)) + 
  geom_point(aes(size = total, colour = total)) +
  scale_color_gradientn(colours = pal) +
  theme_bw() + 
  theme(axis.text.x = element_text(angle = 45, hjust = 1, vjust = 1),
        legend.text.position = "right")
```

Figure 4: **Ant community composition varies across localities.** **(A)** Non-metric multidimensional scaling (NMDS) biplot based on Bray-Curtis dissimilarities showing ant communities across locations based on the relative abundances of 113 ant species (Stress = 0.156). Each triangle represents ant communities at one sampling point. Environmental factors (black arrows) and ant genera (blue arrows) in correlation with the ordination are shown. **(B)** Procrustes ordination plot showing correspondence between frog alkaloids and ant communities ordinations across localities. Procrustes residuals (lines) connect paired points between frog alkaloid profiles (circles) and ant community composition (triangles) at each site. Shorter lines indicate better alignment between the ordinations, suggesting stronger correspondence. **(C)** Bubble - heatmap shows the number of ants, grouped by genus (with ≥10 individuals), across frog populations. Each ant community was sampled using a trap placed next to the corresponding frog’s capture site.

Figure 4.png

Code

```
colnames(pair.per.ants) <- c("Contrasts", "df", "Sums of sq.", "F", 
                        "R2","p-value", "p.adjusted", "Sig.")

T5 <- pair.per.ants %>%
  select(Contrasts, df, F, R2, p.adjusted) %>% 
  mutate(Contrasts = c("La Maná vs. C. colón","La Maná vs. S. Domingo", 
                       "La Maná vs. P. Quito","C. colón vs. S. Domingo",
                       "C. Colón vs. P. Quito","S. Domingo vs. P. Quito")) %>% 
  kable(digits = 3, 
      table.attr = 'data-quarto-disable-processing="true"', "html",
      caption = "Table S5: Summary of the results of pairwise comparisons of a PERMANOVA in leaf litter ant composition between study sites") %>% 
  kable_classic(full_width = F, html_font = "Cambria") %>% 
  row_spec(0, bold = T) 

T5
```

Table 5: Summary of the results of pairwise comparisons of a PERMANOVA in leaf litter ant composition between study sites

Table S5: Summary of the results of pairwise comparisons of a PERMANOVA in leaf litter ant composition between study sites

| Contrasts | df | F | R2 | p.adjusted |
| --- | --- | --- | --- | --- |
| La Maná vs. C. colón | 1 | 2.988 | 0.083 | 0.011 |
| La Maná vs. S. Domingo | 1 | 5.695 | 0.160 | 0.002 |
| La Maná vs. P. Quito | 1 | 4.699 | 0.164 | 0.002 |
| C. colón vs. S. Domingo | 1 | 6.891 | 0.165 | 0.002 |
| C. Colón vs. P. Quito | 1 | 4.059 | 0.123 | 0.005 |
| S. Domingo vs. P. Quito | 1 | 5.911 | 0.185 | 0.002 |

Table 5

#### Frogs show different dietary selectivity for particular ant genera

We next asked if frogs eat specific ant genera selectively or if their ant diet reflects the genera of the surrounding leaf litter communities. From the 46 ant genera recovered from Winkler traps in leaf litter communities, only 17 of these were consumed by frogs across different species and populations. Our results indicated no significant differences in the total abundance of ants between frog stomach contents and leaf litter across all populations, except for Santo Domingo and *H. infraguttatus* where frogs had lower abundance of ants in their stomachs compared to the leaf litter (Table 6; Figure 5 A). We found that *Solenopsis* was the most selected genus across all *O. sylvatica* populations, while *Pheidole* was the only selected genus by *H. infraguttatus*. Particularly, frogs from the Cristóbal Colón population showed selectivity for *Paratrachymyrmex*, *Crematogaster* and *Pheidole*, whereas *Strumigenys* and *Cyphomyrmex* were selected in La Maná and Puerto Quito populations, respectively (Figure 5 B).

Code

```
f5a <- sum_abun %>% 
  ggplot(aes(x = group, y = abundance, fill = population)) +
  geom_boxplot(outlier.shape = NA) + 
  theme_bw() +
  theme(panel.grid.major = element_blank(),
                                    panel.grid.minor = element_blank(),
                                    axis.text.x = NULL, legend.position = "none") +
  geom_jitter(position=position_jitterdodge(0.1), shape = 21, 
              size = 2.5, alpha=0.6) +
  scale_fill_manual(values = c("#e8c95d", "#d16b54","#ffffff", "#a9d8c8",
                               "#433447", "#b9b09f")) +
  facet_grid(.~population) +
  scale_y_sqrt() + 
  scale_y_continuous(trans = 'sqrt') +
  ylab(NULL)

#ggsave("Fig4a.svg", f4a, units = "cm", width = 18, height = 8)


f5b <- ggplot() + 
  geom_point(data = Lsel_sim, aes(x = value, y = Genus), col = "gray", 
             pch = 15, size = 4) +
  geom_point(data = Lsel_obs, aes(x = Linear, y = Genus, fill = Cat.Linear), size = 2.5, pch = 21) +
  scale_fill_manual(values=c("black", "white", "blue")) +
  facet_grid(.~population, scales = "free")  +
  theme_bw() + 
  theme(panel.grid.major = element_blank(),
        panel.grid.minor = element_blank(),
        axis.text.x = NULL, 
        legend.position = "none") +
  xlab("Linear Selectivity") +
  ylab("Ant species")

#ggsave("Fig4b.svg", f4b, units = "cm", width = 20, height = 7)
```

Figure 5: **Relative abundance and selectivity for ant genera differs across localities.** (A)\*\* Boxplot showing differences across populations in total abundance of ants within 17 ant genera found in both leaf litter and frog stomach samples. The y axis is square-root transformed for visual clarity. n.s. = non-significant. \* p-value<0.05 **(B)** Linear selectivity index for 17 ant genera eaten by the toxic *O. sylvatica* populations and the non-toxic *H. infraguttatus*. Grey bars denote simulated null distribution. Points denote categorical selectivity as follows: ‘non-selected’ if they are below the null distribution (red dots), ‘neutral’ if they are within (black dots), and ‘selected’ if the values are above (blue dots). Blue arrows indicate overall selected ant genera.

Figure 5.png

On the other hand, The remaining ants were either not selected or occasionally consumed (Figure 5 B). For example, *Wasmannia* ants were avoided by all populations, except for Santo Domingo frogs where it was consumed in proportion to its availability (i.e., neutral). Similarly, *Apterostigma* ants were avoided by frogs in Santo Domingo and in the sympatric populations of *O. sylvatica* and *H. infraguttatus* in La Maná.

Code

```
t6 <- data.frame(sumabun.emmeans$`pairwise differences of group, population`)

conts <- c("frog c_colon - leaf c_colon", "frog p_quito - leaf p_quito", "frog s_domingo - leaf s_domingo", "frog la_mana - leaf la_mana", "frog H_infraguttatus - leaf H_infraguttatus")

T6 <- t6 %>%
  select(X1, estimate, SE, p.value) %>% 
  mutate(Contrasts = X1) %>% 
  select(Contrasts, estimate, SE, p.value) %>% 
  filter(Contrasts %in% conts) %>% 
  mutate(Contrasts = c("frog C. Colón vs. leaf C. Colón", "frog P. Quito vs. leaf P. Quito", 
                   "frog S. Domingo vs. leaf S. Domingo", "frog La Maná vs. leaf La Maná", 
                   "frog H. infraguttatus vs. leaf H. infraguttatus")) %>% 
  select(Contrasts, estimate, SE, p.value) %>% 
  kable(digits = 3, 
      table.attr = 'data-quarto-disable-processing="true"', "html",
      caption = "Table S6: Summary of the results of pairwise comparisons of a Negative Binomial regresion comparing ant abundance between *O. sylvatica* populations. P-values were adjusted using Tukey’s method") %>% 
  kable_classic(full_width = F, html_font = "Cambria") %>% 
  row_spec(0, bold = T)

T6
```

Table S6: Summary of the results of pairwise comparisons of a Negative Binomial regresion comparing ant abundance between \*O. sylvatica\* populations. P-values were adjusted using Tukey’s method

| Contrasts | estimate | SE | p.value |
| --- | --- | --- | --- |
| frog C. Colón vs. leaf C. Colón | -0.452 | 0.263 | 0.828 |
| frog P. Quito vs. leaf P. Quito | -1.220 | 0.320 | 0.007 |
| frog S. Domingo vs. leaf S. Domingo | 0.533 | 0.230 | 0.423 |
| frog La Maná vs. leaf La Maná | -0.668 | 0.259 | 0.261 |
| frog H. infraguttatus vs. leaf H. infraguttatus | -1.084 | 0.262 | 0.002 |

Table 6: Summary of the results of pairwise comparisons of a Negative Binomial regresion comparing ant abundance between *O. sylvatica* populations. P-values were adjusted using Tukey’s method

Table 6

#### Ant morphology influences prey selectivity in frogs

Finally, we asked whether *O. sylvatica* frogs select for specific ant traits. A principal components analysis on 17 morphological and life history ant traits yielded three principal components, each with eigenvalues exceeding 1. After inverting the scores of these components to aid interpretability, we found that the first component accounted for 55.66% of the total variance, primarily reflecting size and texture traits, with higher scores indicating larger ants. Component number two explained 11.02% of the variance and was strongly associated with color traits, where higher scores corresponded to ants with dark-brown body coloration. The third principal component, explaining 7.15% of the variance, was largely indicative of the presence of spines, with higher scores representing spiny ants (Figure 6). A pairwise Wilcoxon test on these principal components revealed that ants categorized as “selected” were significantly smaller than those with neutral selectivity (p-value = 0.03; Figure 6 A; Table 7), but not than those not selected (p-value = 0.162; Figure 6 A). Contrary, no differences were found in the traits of color and the presence of spines across the categories of selectivity (all p values >0.05; Figure 6 B & C; Table 7).

Code

```
# Plot contributions of each variable to each component 
fig6a <- PCAtraits$co %>% 
  data.frame() %>% 
  rownames_to_column(var = "Trait") %>% 
  select(Trait, Comp1, Comp2, Comp3) %>%
  melt() %>% 
  arrange(desc(value)) %>% 
  arrange(desc(variable)) %>% 
  filter(variable %in% c("Comp1", "Comp2", "Comp3")) %>% 
  mutate(Trait = fct_relevel(Trait, "WebersL", "BodyL", "HeadL", "HindFemurL", "PronotumW",
                                "HeadW", "InterOcularW", "MandibleL", "EyeL", "ScapeL", "ClypeusL",
                                "Pilosity", "Sculpturing", "nSpines", "ColourGaster", "Colour.Mesosoma",
                                "ColourHead")) %>% 
  mutate(value = value*-1) %>% 
  mutate(sign = ifelse(value >= 0, "Positive", "Negative")) %>% 
  ggplot(aes(x = Trait, y = value, fill = interaction(variable, sign))) + 
  geom_bar (stat="identity",position = position_dodge(0.9), col = "black", lwd = 0.2) +
  scale_y_continuous(limits = c(-1,1)) +
  facet_grid(.~variable) + coord_flip() +
  geom_hline(yintercept = 0, linetype = 3, col = "grey") +
  scale_fill_manual(values = c("Comp1.Positive" = "#a9d8c8", "Comp2.Positive" = "orange", 
                               "Comp3.Positive" = "beige", "Comp1.Negative" = "gray", 
                               "Comp2.Negative" = "gray", "Comp3.Negative" = "gray")) + 
  theme_bw() + theme(panel.grid.major = element_blank(),
                     panel.grid.minor = element_blank(),
                     axis.text.x = NULL, legend.position = "none") 

#Generate data with the mean of first three components for every Genus and site, and linear selectivity (continuous and categorical)
elect.morph <- na.omit(ant_traits) %>% 
  filter(Genus %in% Lsel_obs$Genus) %>% 
  bind_cols(PCAtraits$li) %>% # merge with principal components
  select(Genus, HeadW:ColourGaster, Axis1, Axis2, Axis3) %>% #select only the first three components
  mutate(Axis1 = Axis1*-1, Axis2 = Axis2*-1, Axis3 = Axis3*-1) %>% #PCs are multiplied by -1 for better interpretation 
  rename(PC1 =Axis1, PC2 = Axis2, PC3 = Axis3) %>% 
  group_by(Genus) %>% 
  summarise(mPC1 = mean(PC1), mPC2 = mean(PC2), mPC3 = mean(PC3)) %>%
  left_join(Lsel_obs) %>% #merge with linear selectivity data
  group_by(population) %>% 
  arrange(desc(Linear)) %>%
  mutate(Genus = reorder(Genus, Linear)) %>%
  ungroup() %>% 
  mutate(Cat.Linear = factor(Cat.Linear, levels = c("preferred", "neutral", "avoid")))

# Plot principal components against electivity categories
fig6b <- elect.morph %>% 
  select(-variable) %>% 
  melt(variable.name = "Component") %>% 
  filter(Component %in% c("mPC1", "mPC2", "mPC3")) %>% 
  ggplot(aes(Cat.Linear, value, fill = Cat.Linear)) + 
  geom_boxplot(outlier.shape = NA) + 
  scale_fill_manual(values = c("#dfb92aff", "#88b000ff", "#3f98c8ff")) +
  geom_jitter(position=position_jitterdodge(0.5), shape = 21, 
              size = 4, alpha=0.6) + 
  facet_wrap(.~Component, scales = "free_y") +
  theme_bw() + theme(panel.grid.major = element_blank(),
                     panel.grid.minor = element_blank(),
                     axis.text.x = NULL, legend.position = "none")
```

Figure 6: **Boxplots showing differences between categorical selectivity in principal component scores on 17 ant traits.** (A)\*\* First principal component (PC1) representing size and texture traits. High score values depict larger ants and low score values depict smaller ants. **(B)** Second principal component (PC2) representing body coloration. High score values depict dark-brown ants, middle scores depict light-brown ants, and low score values depict black ants. **(C)** Third principal component (PC3) representing number of spines. High score values depict *spiny* ants, whule low score values depict *non-spiny* ants. Groups not connected by the same letter are significantly different.

Figure 6.png

Code

```
# Pairwise comparisons between linear selectivity categories in the principal components

pwt1 <- pairwise.wilcox.test(elect.morph$mPC1, elect.morph$Cat.Linear, p.adjust.method = "BH")

pwt2 <- pairwise.wilcox.test(elect.morph$mPC2, elect.morph$Cat.Linear, p.adjust.method = "BH")

pwt3 <- pairwise.wilcox.test(elect.morph$mPC3, elect.morph$Cat.Linear, p.adjust.method = "BH")

T7 <- cbind(pwt1$p.value, pwt2$p.value, pwt3$p.value) %>%
  set_colnames(sub("preferred", "selected", colnames(.))) %>% 
  kable(digits = 3, 
      table.attr = 'data-quarto-disable-processing="true"', "html",
      caption = "Table S7: P-values of pairwise Wilcoxon test on differences in principal components between linear selectivity categories") %>% 
  kable_classic(full_width = F, html_font = "Cambria") %>% 
  row_spec(0, bold = T) %>% 
  add_header_above(c("", "PC1 (size & texture)" = 2, 
                     "PC2 (Color)" = 2,
                     "PC3 (Spines)" = 2))

T7
```

Table S7: P-values of pairwise Wilcoxon test on differences in principal components between linear selectivity categories

|  | PC1 (size & texture) | | PC2 (Color) | | PC3 (Spines) | |
| --- | --- | --- | --- | --- | --- | --- |
|  | selected | neutral | selected | neutral | selected | neutral |
| neutral | 0.008 | - | 0.865 | - | 0.403 | - |
| avoid | 0.251 | 0.113 | 0.157 | 0.106 | 0.547 | 0.388 |

Table 7: P-values of pairwise Wilcoxon test on differences in principal components between linear selectivity categories

##### Funding

This work was supported by the National Science Foundation (IOS-1557684 and IOS-2337580), Pew Charitable Trusts, and the New York Stem Cell Foundation to LAO. NAM and AAB are supported by National Science Foundation graduate research fellowships (DGE-1656518). AAB is supported by a Gilliam Fellowship from the Howard Hughes Medical Institute (GT13330). LAO is a New York Stem Cell Foundation – Robertson Investigator.

##### Author contributions

Conceptualization: NAM, LAO. Methodology: NAM, LAO, DAD, AAB. Formal Analysis: NAM, AAB,CR. Investigation: NAM, AAB, AC, JR, DAD. Resources: LAO, DAD. Data Curation: NAM, AAB, CM, DAD, CR. Writing – original draft preparation: NAM, LAO, AAB, DAD. Writing – review and editing: NAM, LAO, AAB, DAD, ET, AC, JR, LAC, CR, KF. Visualization: NAM, LAO, CR. Supervision: LAO. Project administration: LAO. Funding acquisition: LAO.

Deslippe, Richard J, and Yu-Jie Guo. 2000. “Venom Alkaloids of ®Re Ants in Relation to Worker Size and Age.”

Donnelly, Maureen A. 1991. “Feeding Patterns of the Strawberry Poison Frog, Dendrobates Pumilio (Anura: Dendrobatidae).” *Copeia* 1991 (3): 723. https://doi.org/10.2307/1446399.

Donoso, David A. 2017. “Tropical Ant Communities Are in Long-Term Equilibrium.” *Ecological Indicators* 83 (December): 515–23. https://doi.org/10.1016/j.ecolind.2017.03.022.

Donoso, David A., and Giovanni Ramón. 2009. “Composition of a High Diversity Leaf Litter Ant Community (Hymenoptera: Formicidae) from an Ecuadorian Pre-Montane Rainforest.” *Annales de La Société Entomologique de France (N.S.)* 45 (4): 487–99. https://doi.org/10.1080/00379271.2009.10697631.

Dray, Stéphane, and Anne–Béatrice Dufour. 2007. “The Ade4 Package: Implementing the Duality Diagram for Ecologists” 22. https://doi.org/10.18637/jss.v022.i04.

Fick, Stephen E., and Robert J. Hijmans. 2017. “WorldClim 2: New 1-Km Spatial Resolution Climate Surfaces for Global Land Areas.” *International Journal of Climatology* 37 (12): 4302–15. https://doi.org/10.1002/joc.5086.

Funkhouser, John W. 1956. “New Frogs from Ecuador and Southwestern Colombia.” *Zoologica : Scientific Contributions of the New York Zoological Society* 41 (9). https://doi.org/10.5962/p.190356.

Gibb, Heloise, Nathan J. Sanders, Robert R. Dunn, Simon Watson, Manoli Photakis, Silvia Abril, Alan N. Andersen, et al. 2015. “Climate Mediates the Effects of Disturbance on Ant Assemblage Structure.” *Proceedings of the Royal Society B: Biological Sciences* 282 (1808): 20150418. https://doi.org/10.1098/rspb.2015.0418.

Hoenle, Philipp O., David A. Donoso, Adriana Argoti, Michael Staab, Christoph von Beeren, and Nico Blüthgen. 2022. “Rapid Ant Community Reassembly in a Neotropical Forest: Recovery Dynamics and Land-Use Legacy.” *Ecological Applications* 32 (4): e2559. https://doi.org/10.1002/eap.2559.

Jeckel, Adriana M., Sophie Kocheff, Ralph A. Saporito, and Taran Grant. 2019. “Geographically Separated Orange and Blue Populations of the Amazonian Poison Frog Adelphobates Galactonotus (Anura, Dendrobatidae) Do Not Differ in Alkaloid Composition or Palatability.” *Chemoecology* 29 (5-6): 225–34. https://doi.org/10.1007/s00049-019-00291-3.

Jones, Tappey H., Murray S. Blum, and Henry M. Fales. 1982. “Ant Venom Alkaloids from *Solenopsis* and *Monorium* Species.” *Tetrahedron*, The organic chemistry of animal defense mechanisms, 38 (13): 1949–58. https://doi.org/10.1016/0040-4020(82)80044-6.

Jones, Tappey H., Jeffrey S. T. Gorman, Roy R. Snelling, Jacques H. C. Delabie, Murray S. Blum, H. Martin Garraffo, Poonam Jain, John W. Daly, and Thomas F. Spande. 1999. “Further Alkaloids Common to Ants and Frogs: Decahydroquinolines and a Quinolizidine.” *Journal of Chemical Ecology* 25 (5): 1179–93. https://doi.org/10.1023/A:1020898229304.

Lawrence, J. P., Bibiana Rojas, Annelise Blanchette, Ralph A. Saporito, Johanna Mappes, Antoine Fouquet, and Brice P. Noonan. 2023. “Linking Predator Responses to Alkaloid Variability in Poison Frogs.” *Journal of Chemical Ecology* 49 (3): 195–204. https://doi.org/10.1007/s10886-023-01412-7.

Lenth, Russell V. 2025. “Emmeans: Estimated Marginal Means, Aka Least-Squares Means.” https://CRAN.R-project.org/package=emmeans.

Mackay, William P., Solange Silva, David C. Lightfoot, Maria Inez Pagani, and Walter G. Whitford. 1986. “Effect of Increased Soil Moisture and Reduced Soil Temperature on a Desert Soil Arthropod Community.” *American Midland Naturalist* 116 (1): 45. https://doi.org/10.2307/2425936.

McElroy, Matthew T., and David A. Donoso. 2019. “Ant Morphology Mediates Diet Preference in a Neotropical Toad (Rhinella Alata).” *Copeia* 107 (3): 430. https://doi.org/10.1643/CH-18-162.

McGugan, Jenna R., Gary D. Byrd, Alexandre B. Roland, Stephanie N. Caty, Nisha Kabir, Elicio E. Tapia, Sunia A. Trauger, Luis A. Coloma, and Lauren A. O’Connell. 2016. “Ant and Mite Diversity Drives Toxin Variation in the Little Devil Poison Frog.” *Journal of Chemical Ecology* 42 (6): 537–51. https://doi.org/10.1007/s10886-016-0715-x.

Mebs, D. 2002. *Venomous and Poisonous Animals: A Handbook for Biologists, Toxicologists and Toxinologists, Physicians and Pharmacists*. CRC Press.

Meurer, William, Felipe G. Gonçalves, Ricardo S. Bovendorp, Alexandre R. Percequillo, and Jaime Bertoluci. 2021. “Diet Electivity and Preferences for Food Resources in Chiasmocleis Leucosticta (Anura: Microhylidae).” *Journal of Herpetology* 55 (4): 325–29. https://doi.org/10.1670/20-106.

Moses, Jimmy, Tom M. Fayle, Vojtech Novotny, and Petr Klimes. 2021. “Elevation and Leaf Litter Interact in Determining the Structure of Ant Communities on a Tropical Mountain.” *Biotropica* 53 (3): 906–19. https://doi.org/10.1111/btp.12914.

Moskowitz, Nora A., Rachel D’Agui, Aurora Alvarez-Buylla, Katherine Fiocca, and Lauren A. O’Connell. 2022. “Poison Frog Dietary Preference Depends on Prey Type and Alkaloid Load.” Edited by Mainul Haque. *PLOS ONE* 17 (12): e0276331. https://doi.org/10.1371/journal.pone.0276331.

Moskowitz, Nora A., Barbara Dorritie, Tammy Fay, Olivia C. Nieves, Charles Vidoudez, Cambridge Rindge Latin 2017 Biology Class, Masconomet 2017 Biotechnology Class, et al. 2020. “Land Use Impacts Poison Frog Chemical Defenses Through Changes in Leaf Litter Ant Communities.” *Neotropical Biodiversity* 6 (1): 75–87. https://doi.org/10.1080/23766808.2020.1744957.

Moskowitz, Nora A., Alexandre B. Roland, Eva K. Fischer, Ndimbintsoa Ranaivorazo, Charles Vidoudez, Marianne T. Aguilar, Sophia M. Caldera, et al. 2018. “Seasonal Changes in Diet and Chemical Defense in the Climbing Mantella Frog (Mantella Laevigata).” Edited by Alex V. Chaves. *PLOS ONE* 13 (12): e0207940. https://doi.org/10.1371/journal.pone.0207940.

Myers, Charles W., and John W. Daly. 1976. “Preliminary evaluation of skin toxins and vocalizations in taxonomic and evolutionary studies of poison-dart frogs (Dendrobatidae). Bulletin of the AMNH ; v. 157, article 3.” http://hdl.handle.net/2246/622.

Nayik, Gulzar Ahmad, and Jasmeet Kour, eds. 2022. *Handbook of Plant and Animal Toxins in Food: Occurrence, Toxicity, and Prevention*. Boca Raton: CRC Press. https://doi.org/10.1201/9781003178446.

Oksanen, Jari, Gavin L. Simpson, F. Guillaume Blanchet, Roeland Kindt, Pierre Legendre, Peter R. Minchin, R. B. O’Hara, et al. 2024. “Vegan: Community Ecology Package.” https://CRAN.R-project.org/package=vegan.

Olson, David M. 1991. “A Comparison of the Efficacy of Litter Sifting and Pitfall Traps for Sampling Leaf Litter Ants (Hymenoptera, Formicidae) in a Tropical Wet Forest, Costa Rica.” *Biotropica* 23 (2): 166. https://doi.org/10.2307/2388302.

Parr, Catherine L., Robert R. Dunn, Nathan J. Sanders, Michael D. Weiser, Manoli Photakis, Tom R. Bishop, Matthew C. Fitzpatrick, et al. 2017. “GlobalAnts: A New Database on the Geography of Ant Traits (Hymenoptera: Formicidae).” *Insect Conservation and Diversity* 10 (1): 5–20. https://doi.org/10.1111/icad.12211.

Prates, Ivan, Andrea Paz, Jason L. Brown, and Ana C. Carnaval. 2019. “Links Between Prey Assemblages and Poison Frog Toxins: A Landscape Ecology Approach to Assess How Biotic Interactions Affect Species Phenotypes.” *Ecology and Evolution* 9 (24): 14317–29. https://doi.org/10.1002/ece3.5867.

Roberts, Margaret F., and Michael Wink, eds. 1998. *Alkaloids: Biochemistry, Ecology, and Medicinal Applications*. Boston, MA: Springer US. https://doi.org/10.1007/978-1-4757-2905-4.

Sabu, Thomas K., and Raj T. Shiju. 2010. “Efficacy of Pitfall Trapping, Winkler and Berlese Extraction Methods for Measuring Ground-Dwelling Arthropods in Moistdeciduous Forests in the Western Ghats.” *Journal of Insect Science* 10 (1): 98. https://doi.org/10.1673/031.010.9801.

Salazar, Fernanda, Fabian Reyes-Bueno, Daniel Sanmartin, and David A Donoso. 2015. “Mapping Continental Ecuadorian Ant Species.” *Sociobiology* 62 (2): 132–62. https://doi.org/10.13102/sociobiology.v62i2.132-162.

Sanches, Patrick R., Luã E. Santos-Guerra, Fillipe Pedroso-Santos, Igor L. Kaefer, and Carlos E. Costa-Campos. 2023. “What Do Co-Mimics Eat? Trophic Ecology of Ameerega Pulchripecta (Anura, Dendrobatidae) and Allobates Femoralis (Anura, Aromobatidae) in Eastern Brazilian Amazonia.” *Journal of Herpetology* 57 (4). https://doi.org/10.1670/22-074.

Sánchez Loja, Santiago, David A. Donoso, and Monica Isabel Paez Vacas. 2023. “Conspicuous and Cryptic Poison Frogs Are Picky and Prefer Different Meals in Syntopy,” December. https://doi.org/10.1007/s10682-023-10282-0.

Santos, Juan Carlos, Luis A. Coloma, and David C. Cannatella. 2003. “Multiple, Recurring Origins of Aposematism and Diet Specialization in Poison Frogs.” *Proceedings of the National Academy of Sciences* 100 (22): 12792–97. https://doi.org/10.1073/pnas.2133521100.

Santos, Juan C., Rebecca D. Tarvin, and Lauren A. O’Connell. 2016. “A Review of Chemical Defense in Poison Frogs (Dendrobatidae): Ecology, Pharmacokinetics, and Autoresistance.” In, edited by Bruce A. Schulte, Thomas E. Goodwin, and Michael H. Ferkin, 305–37. Cham: Springer International Publishing. https://doi.org/10.1007/978-3-319-22026-0\_21.

Saporito, Ralph A., Maureen A. Donnelly, H. Martin Garraffo, Thomas F. Spande, and John W. Daly. 2006. “Geographic and Seasonal Variation in Alkaloid-Based Chemical Defenses of Dendrobates Pumilio from Bocas Del Toro, Panama.” *Journal of Chemical Ecology* 32 (4): 795–814. https://doi.org/10.1007/s10886-006-9034-y.

Saporito, Ralph A., Maureen A. Donnelly, Poonam Jain, H. Martin Garraffo, Thomas F. Spande, and John W. Daly. 2007. “Spatial and Temporal Patterns of Alkaloid Variation in the Poison Frog *Oophaga Pumilio* in Costa Rica and Panama over 30 Years.” *Toxicon* 50 (6): 757–78. https://doi.org/10.1016/j.toxicon.2007.06.022.

Saporito, Ralph A., Maureen A. Donnelly, Anne A. Madden, H. Martin Garraffo, and Thomas F. Spande. 2010. “Sex-Related Differences in Alkaloid Chemical Defenses of the Dendrobatid Frog Oophaga Pumilio from Cayo Nancy, Bocas Del Toro, Panama.” *Journal of Natural Products* 73 (3): 317–21. https://doi.org/10.1021/np900702d.

Saporito, Ralph A., Maureen A. Donnelly, Roy A. Norton, H. Martin Garraffo, Thomas F. Spande, and John W. Daly. 2007. “Oribatid Mites as a Major Dietary Source for Alkaloids in Poison Frogs.” *Proceedings of the National Academy of Sciences* 104 (21): 8885–90. https://doi.org/10.1073/pnas.0702851104.

Saporito, Ralph A., H. Martin Garraffo, Maureen A. Donnelly, Adam L. Edwards, John T. Longino, and John W. Daly. 2004. “Formicine Ants: An Arthropod Source for the Pumiliotoxin Alkaloids of Dendrobatid Poison Frogs.” *Proceedings of the National Academy of Sciences* 101 (21): 8045–50. https://doi.org/10.1073/pnas.0402365101.

Saporito, Ralph A., Roy A. Norton, Nirina R. Andriamaharavo, Hugo Martin Garraffo, and Thomas F. Spande. 2011. “Alkaloids in the Mite Scheloribates Laevigatus: Further Alkaloids Common to Oribatid Mites and Poison Frogs.” *Journal of Chemical Ecology* 37 (2): 213–18. https://doi.org/10.1007/s10886-011-9914-7.

Saporito, Ralph A., Roy A. Norton, Martin H. Garraffo, and Thomas F. Spande. 2015. “Taxonomic Distribution of Defensive Alkaloids in Nearctic Oribatid Mites (Acari, Oribatida).” *Experimental and Applied Acarology* 67 (3): 317–33. https://doi.org/10.1007/s10493-015-9962-8.

Savitzky, Alan H., Akira Mori, Deborah A. Hutchinson, Ralph A. Saporito, Gordon M. Burghardt, Harvey B. Lillywhite, and Jerrold Meinwald. 2012. “Sequestered Defensive Toxins in Tetrapod Vertebrates: Principles, Patterns, and Prospects for Future Studies.” *Chemoecology* 22 (3): 141–58. https://doi.org/10.1007/s00049-012-0112-z.

Silva, Rogério R., and Carlos Roberto F. Brandão. 2014. “Ecosystem-Wide Morphological Structure of Leaf-Litter Ant Communities Along a Tropical Latitudinal Gradient.” *PLOS ONE* 9 (3): e93049. https://doi.org/10.1371/journal.pone.0093049.

Spande, Thomas F., Poonam Jain, H. Martin Garraffo, Lewis K. Pannell, Herman J. C. Yeh, John W. Daly, Sinji Fukumoto, et al. 1999. “Occurrence and Significance of Decahydroquinolines from Dendrobatid Poison Frogs and a Myrmicine Ant:  Use of 1H and 13C NMR in Their Conformational Analysis.” *Journal of Natural Products* 62 (1): 5–21. https://doi.org/10.1021/np980298v.

Strauss, Richard E. 1979. “Reliability Estimates for Ivlevˈs Electivity Index, the Forage Ratio, and a Proposed Linear Index of Food Selection.” *Transactions of the American Fisheries Society* 108 (4): 344–52. https://doi.org/10.1577/1548-8659(1979)108<344:REFIEI>2.0.CO;2.

Stuckert, Adam MM, Ralph A. Saporito, Pablo J. Venegas, and Kyle Summers. 2014. “Alkaloid Defenses of Co-Mimics in a Putative Müllerian Mimetic Radiation.” *BMC Evolutionary Biology* 14 (1): 76. https://doi.org/10.1186/1471-2148-14-76.

Tarvin, Rebecca D, Jeffrey L Coleman, David A Donoso, Mileidy Betancourth-Cundar, Karem López-Hervas, Kimberly S Gleason, J Ryan Sanders, et al. 2024. “Passive Accumulation of Alkaloids in Inconspicuously Colored Frogs Refines the Evolutionary Paradigm of Acquired Chemical Defenses.” https://doi.org/10.7554/eLife.100011.2.

Tiede, Yvonne, Jan Schlautmann, David A. Donoso, Christine I. B. Wallis, Jörg Bendix, Roland Brandl, and Nina Farwig. 2017. “Ants as Indicators of Environmental Change and Ecosystem Processes.” *Ecological Indicators* 83 (December): 527–37. https://doi.org/10.1016/j.ecolind.2017.01.029.

Toft, Catherine A. 1995. “Evolution of Diet Specialization in Poison-Dart Frogs (Dendrobatidae).” *Herpetologica* 51 (2): 202–16. https://www.jstor.org/stable/3892588.

Venables, W. N., and B. D. Ripley. 2002. “Modern Applied Statistics with s.” https://www.stats.ox.ac.uk/pub/MASS4/.

Virjamo, V., and R. Julkunen-Tiitto. 2016. “Variation in Piperidine Alkaloid Chemistry of Norway Spruce (Picea Abies) Foliage in Diverse Geographic Origins Grown in the Same Area.” *Canadian Journal of Forest Research* 46 (4): 456–60. https://doi.org/10.1139/cjfr-2015-0388.

Walsh, Christopher T., and Yi Tang. 2017. *Natural Product Biosynthesis: Chemical Logic and Enzymatic Machinery*. Royal Society of Chemistry.

Wang, Mingxun, Jeremy J. Carver, Vanessa V. Phelan, Laura M. Sanchez, Neha Garg, Yao Peng, Don Duy Nguyen, et al. 2016. “Sharing and Community Curation of Mass Spectrometry Data with Global Natural Products Social Molecular Networking.” *Nature Biotechnology* 34 (8): 828–37. https://doi.org/10.1038/nbt.3597.

Weldon, Paul J. 2017. “Poison Frogs, Defensive Alkaloids, and Sleepless Mice: Critique of a Toxicity Bioassay.” *Chemoecology* 27 (4): 123–26. https://doi.org/10.1007/s00049-017-0238-0.

Wise, David H., and Janet R. Lensing. 2019. “Impacts of Rainfall Extremes Predicted by Climate-Change Models on Major Trophic Groups in the Leaf Litter Arthropod Community.” *Journal of Animal Ecology* 88 (10): 1486–97. https://doi.org/10.1111/1365-2656.13046.


###### Source Code

```
---
title: "Poison frog chemical defenses are influenced by environmental availability and dietary selectivity for ants"
toc: true
toc-depth: 4
toc-location: left
freeze: true
format: 
  html:
    embed-resources: true
    code-fold: true
    code-copy: true
    code-tools: true
    highlight-style: tango
    self-contained: true
    standalone: true
    fig-responsive: true
editor: visual
bibliography: references.bib
editor_options: 
  chunk_output_type: console
---

```{=html}
<style>
table caption {
    text-align: left !important;
}
</style>
```

```{=html}
<style>
p {
  text-indent: 1.5em;
}

.authors-section p {
  text-indent: 0; /* Remove indentation for paragraphs inside authors-section */
}
</style>
```

::: authors-section
Nora A. Martin†¹, Camilo Rodríguez†¹, Aurora Alvarez-Buylla¹, Katherine Fiocca¹, Colin R. Morrison², Adolfo Chamba-Carrillo³, Ana B. García-Ruilova⁴, Janet Rentería⁵, Elicio E. Tapia⁶, Luis A. Coloma⁶, David A. Donoso\*⁷⁺⁸, Lauren A. O’Connell\*¹<br><br>

¹ Department of Biology, Stanford University, Stanford, CA 94305, USA<br> ² Department of Integrative Biology, The University of Texas at Austin, Austin, TX 78712, USA<br> ³ Programa de Posgrado en Biodiversidad y Cambio Climático, Universidad Indoamérica, Quito, Ecuador<br> ⁴ División de Entomología, Instituto Nacional de Biodiversidad, Pje. Rumipamba N. 341 y Av. de los Shyris, Quito, Ecuador<br> ⁵ School of Biological Sciences, University of Bristol, Bristol, UK <br> ⁶ Centro Jambatu de Investigación y Conservación de Anfibios, Fundación Otonga, San Rafael, Quito, Ecuador<br> ⁷ Departamento de Biología, Escuela Politécnica Nacional, Ladrón de Guevara E11-253, Quito, Ecuador<br> ⁸ Grupo de Investigación en Ecología y Evolución en los Trópicos -EETrop-, Universidad de las Américas, Quito, Ecuador<br><br>

† Joint first authorship<br> \* Corresponding authors: <a href="mailto:">david.donosov\@gmail.com</a>, <a href="mailto:">loconnel\@stanford.edu</a>
:::

```{r}
#| echo: false
#| fig-align: default
#| fig-show: "hold"
#| column: body-outset-right
#| fig-cap: "Graphical abstract"

library(grid)
library(png)

grid.raster(readPNG('/Users/camilorl/Library/CloudStorage//Shared drives/LOBSU Manuscripts/O. sylvatica vs. H. infraguttatus diet (Nora)/Submission 2-JAE/Graphical_abstract.png')) 
```

# Abstract

::: {.callout-note appearance="minimal" icon="false"}
The ability to use small molecule alkaloids as defensive chemicals, often acquired via trophic interactions, has evolved in many organisms. Animals with diet-derived defenses must balance food choices to maintain their defense reservoirs along with other physiological needs. Poison frogs accumulate skin alkaloids from their arthropod diet, but whether they show preference for specific prey remains unexplored. Here, we explore the role of leaf litter prey availability and diet preferences in shaping poison frog chemical defenses along a geographic gradient. We examined skin alkaloid composition, stomach contents and leaf litter ants in aposematic diablito frogs (*Oophaga sylvatica*) at five sites in northwestern Ecuador, and in sympatric, cryptic Chimbo rocket frogs (*Hyloxalus infraguttatus*) at one site. We found that differential availability of leaf litter ants influenced alkaloid profiles across diablito populations, and low levels of alkaloids were observed in the sympatric, ‘undefended’ Chimbo rocket frog. Ants were the primary dietary component of the defended species, while ‘undefended’ species ate other prey categories including beetles and larvae in addition to ants. A prey selection analysis suggested that defended and ‘undefended' frogs both feed on a high proportion of specific small ant genera that naturally contain alkaloids, suggesting that selectivity for toxic prey is not restricted to classically aposematic and highly toxic species. These findings suggest that poison frogs’ use of feeding resources relative to availability may be an understudied and important selection factor in the evolution of acquired defenses.
:::

# Introduction

Many organisms use chemical defenses to protect themselves from predators or pathogens [@mebs2002]. These defenses often involve small molecule alkaloids synthesized by plants or microbes, and some taxa can acquire them through dietary sequestration [@roberts_alkaloids_1998; @agrawal_toxic_2012; @santos2016]. Phytophagous insects represent the most well-studied taxa, including some species that specialize in particular plant species and accumulate specific secondary metabolites for chemical communication or defense [@roberts_alkaloids_1998; @walsh2017; @beran2022]. Although our understanding of chemical defense in vertebrates is more sparse than invertebrates, poison frogs are a well known example of chemical defenses acquired through an arthropod-based diet [@savitzky2012]. A key component of poison frog diets includes alkaloid-rich arthropods, and these frogs have evolved the physiological mechanisms to tolerate and integrate the toxin in their tissues to deter predators [@alvarez-buylla2023]. Yet, it remains unclear whether the ecological principles underlying arthropod diet specialization on toxic plants also apply to vertebrates that sequester their chemical defenses from arthropods.

The chemical repertoire of alkaloid-defended species varies within and between populations. For example, the composition and concentration of piperidine alkaloids in the Norwegian spruce (*Picea abies*) differ by location [@virjamo2016], while in fire ants (*Solenopsis spp.*) vary within species [@deslippe2000]. In species with acquired chemical defenses, variation in alkaloid profiles is generally attributed to spatio-temporal shifts in the availability of alkaloid-containing food, often resulting from environmental variation in temperature, rainfall, and other climatic factors. For instance, in Argentine *Melanophryniscus* toads [@daly2007], Malagasy *Mantella laevigata* frogs [@moskowitz2018], and various species of Neotropical poison frogs (Family Dendrobatidae) [@saporito_geographic_2006; @saporito_spatial_2007; @moskowitz_land_2020; @prates2019], differences in the composition of skin alkaloids across localities and seasons correspond with differences in stomach contents and leaf litter arthropod communities. Dietary selectivity may also influence the alkaloid profile of chemically defended species by favoring the consumption of food items that contain specific defensive alkaloids. Yet, organisms must make dietary decisions based on handling time and nutritional value, in addition to food availability and maintenance of chemical defenses. For instance, *Chiasmocleis leucosticta* frogs preferred smaller ants over larger, more aggressive genera [@meurer2021], while lab-reared non-toxic *Dendrobates tinctorius* preferred protein-rich larvae over other prey types including ants [@moskowitz2022]. As these studies suggest, variation in the environment, the availability of alkaloid-containing prey, prey phenotype and foraging behavior are likely important factors in diet-acquired defense evolution. However, there is limited understanding of how these factors interact to influence species' food choices and their ability to sequester chemical defenses from specific dietary sources.

Neotropical poison frogs acquire alkaloids from alkaloid-containing arthropods rather than synthesizing them de novo [@daly1994]. Chemical defenses in dendrobatids have evolved independently at least four times in parallel with dietary specialization on ants and mites [@santos2003; @darst2005]. However, recent evidence suggests that diet specialization alone does not explain the defended phenotype, as ‘undefended’ species often consume alkaloid-containing arthropods and have low but detectable alkaloid levels [@sanches_what_2023; @tarvin_passive_2024]. Yet, it is unclear if selectivity for specific alkaloid-containing prey plays a role in the ability of poison frogs to dietarily acquire their chemical defenses, as most studies focus on diet without assessing environmental availability of arthropod prey [@mcelroy_ant_2019]. This is especially important as temporal and geographic variations in abiotic factors such as temperature, altitude, and precipitation affect the composition and richness of leaf litter arthropod communities [@brühl1999; @silva2014; @gibb2015; @tiede2017; @hoenle2022; @basset2023]. Thus, we currently lack a framework for understanding the evolution of diet-acquired defenses in poison frogs, as the role of prey availability in diet specialization has not been studied in depth.

Here, we tested whether dendrobatid poison frogs that acquire chemical defenses from their diet exhibit dietary prey selectivity for alkaloid-containing prey. To test this hypothesis, we sampled stomach contents, skin alkaloids, and surrounding leaf litter ant communities from five populations of the aposematic, chemically defended *Oophaga sylvatica* and one sympatric population of the cryptic, chemically ‘undefended’ *Hyloxalus infraguttatus*. We compared skin alkaloid profiles among poison frog populations along a geographical gradient, predicting intraspecific variation linked to environmental factors, such as altitude, temperature and precipitation, and higher alkaloid composition in the sympatric defended species. We compared stomach contents across poison frog populations and predicted within and between species variation, where chemically defended frogs would eat more ants and mites than ‘undefended’ ones. We sorted ants to genus level from stomachs and leaf litter, characterized their morphology, and predicted that sympatric defended and ‘undefended’ frogs will show distinct dietary selectivity for ant genera despite access to the same ant communities. We further predicted that differences in ant communities across localities correspond with differences in chemical defense between poison frogs. Together, our between- and within-species comparisons of poison frog stomach contents and prey availability aim to disentangle the relationship between dendrobatid diet and alkaloid acquisition, which has broader implications for our general understanding of trophic interactions and the evolution of chemical defenses across taxa.

```{r fig1}
#| echo: false
#| warning: false
#| fig-show: "hold"
#| fig-cap: "Map of frog populations and experimental workflow. **(A)** Collection sites are shown on a topographic map of western Ecuador. Note that both *Hyloxalus infraguttatus* and *Oophaga sylvatica* were collected from La Maná. m.a.s.l. = meters above the sea level. **(B)** A flowchart depicts the main steps of data collection for frog stomach content and leaf litter samples, in order to compare ant genera abundances between these groups."
#| cap-location: margin
#| label: fig-1
#| fig-width: 8
#| fig-align: left 

library(grid)
library(png)

grid.raster(readPNG('/Users/camilorl/Library/CloudStorage//Shared drives/LOBSU Manuscripts/O. sylvatica vs. H. infraguttatus diet (Nora)/Submission 2-JAE/Reviews/Fig1_neu.png')) 
```

[Figure 1.](https://drive.google.com/open?id=1oz7vn6lXDQIToIUbWO45K2Yb2mEjzmmm&usp=drive_fs){.external target="_blank"}

# Methods

## Study system and sample collection

Diablito frogs (*Oophaga sylvatica*) were collected in May 2019, during daylight in secondary forests near the towns of Ceiba (N=10; 207 m.a.s.l), Cristóbal Colón (N=11; 221 m.a.s.l), Puerto Quito (N=11; 302 m.a.s.l), Santo Domingo de los Tsáchilas (N=10; 632 m.a.s.l), and La Maná (N=20; 480 m.a.s.l), in Northwestern Ecuador (@fig-1 A). As *O. sylvatica* suffers from illegal poaching for the pet trade, coordinates for collection can be obtained from the corresponding authors. Chimbo rocket frogs (*Hyloxalus infraguttatus*) were collected during daylight from La Maná (N=9). While behavioral observations were not performed, frogs were collected during active foraging hours (06:00 - 19:00) [@funkhouser1956; @amphibia2025]. Frogs were anesthetized 3 – 6 hours after collection with 20% benzocaine gel applied to the ventral skin and euthanized. For each individual, the dorsal skin was dissected and stored in methanol in glass vials. The stomach contents were stored in 100% ethanol in 1.5 ml plastic tubes. Remaining frog tissues were either preserved in 100% ethanol or RNAlater (Thermo Scientific, Waltham, MA, USA), or deposited in the amphibian collection of Centro Jambatu de Investigación y Conservación de Anfibios in Quito, Ecuador. Collections and exportation of specimens were done under permits (Collection permit: No. 0013-18 IC-FAU-DNB/MA; Export permit: No. 214-2019-EXP-CM-FAU-DNB/MA; CITES export permit No. 19EC000036/VS) issued by the Ministerio de Ambiente de Ecuador. The Administrative Panel on Laboratory Animal Care of Stanford University approved all frog-related procedures (Protocol #34153).

## Alkaloid extraction and quantification

Skins were removed from methanol with forceps and weighed. From the methanol in which the skin was stored, 1 ml was syringe filtered through a 0.45 um PTFE filter (Thermo Scientific, 44504-NP) into the new glass vial (Wheaton, PTFE caps, 60940A-2) supplemented with 25 ug (-)-nicotine (Sigma Aldrich, N3876-100ML). All tubes were then capped, vortexed, and stored for 24 hours at -80ºC to precipitate lipids and proteins. After precipitating for 24 hours, the supernatant was filtered through a 0.45 um PTFE syringe filter into a new glass vial. A 100 uL aliquot was added to a gas chromatography/mass spectrometry (GC/MS) autosampler vial, and the remaining solution was stored at -80ºC.

Alkaloid detection was performed using gas chromatography-mass spectrometry (GC-MS) following the protocol described elsewhere [@saporito2010; @alvarez-buylla2023], and using a Shimadzu GCMS-QP2020 instrument with a Shimadzu 30m x 0.25 mmID SH-Rxi-5Sil MS column. In brief, the separation of alkaloids was achieved with helium as the carrier gas (flow rate: 1 mL/min) using a temperature program increasing from 100 to 280°C at a rate of 10°C/minute. This was followed by a 2-minute hold and an additional ramp to 320°C at a rate of 10°C/minute for column protection reasons, and no alkaloids appeared during this part of the method. Compounds were analyzed with electron impact-mass spectrometry (EI-MS). The GC-MS data files were exported as .CDF files, and the Global Natural Products Social Molecular Networking (GNPS) software was used to perform the deconvolution and library searching against the AMDIS (NIST) database to identify all compounds [@wang2016; @aksenov2021]. For deconvolution (identification of peaks and abundance estimates), the default parameters were used. Through the deconvolution process, molecular features were reported as rows/observations, while m/z intensities were reported as columns/variables. Automatic library search was obtained from reference libraries of natural products (NIST, Wiley, University of CORSICA, GNPS), and our resulting dataset was filtered to keep only the nicotine standard and alkaloids previously found in poison frogs or compounds with the same base ring structure and R group positions as those classes defined in the Daly poison frog alkaloid database [@daly2005]. Once the feature table from the GNPS deconvolution was filtered to include only poison frog alkaloids and nicotine, the abundance values (ion counts) were normalized by dividing by the nicotine standard and skin weight. The resulting filtered and normalized feature table was used for all further analyses and visualizations.

## Frog stomach contents identification

Whole stomachs were stored in 100% ethanol at -20ºC until processing. Stomach contents were sorted and photographed. Prey items in the photographs were identified to the lowest possible taxonomic rank and then grouped into broad diet categories: ants, mites, larvae, beetles, and “other” (i.e., any non-larval arthropods not described by the other four groups). Note that most larvae belong to Dipteran or Coleopteran taxa, but are considered in our distinct “larvae” category given their differences in appearance from adults. The vast majority of identifiable prey remained whole, with the exception of ants, whose heads frequently detached from their bodies upon ingestion. To prevent overcounting, only whole ant specimens, partial ant specimens with heads, or individual ant heads were counted. Ant specimens were identified to genus using a reference collection of Ecuadorian ants [@donoso2009; @salazar2015; @donoso2017].

## Leaf litter communities and ant morphology

Leaf litter samples were extracted and collected using 18 - 20 Winkler sacs from all localities except for La Ceiba due to time and resource limitations. Samples were collected in May–June 2019 within one square meter of where a frog had been previously collected that day. Leaf litter arthropods were extracted from 1 m2 and hung within Winkler sacs for 24 hours, during which the arthropods were collected into 70% ethanol. Only ants were identified, as described above. Collection of ant specimens was done under permits issued by the Ministerio de Ambiente de Ecuador to Museo de Historia Natural Gustavo Orcés at Escuela Politécnica Nacional (MAE-DNB-CM-2017-0068). Ant morphology was characterized by 17 traits related to size, texture, spine count, and coloration for leaf litter ant genera that were also found in frog stomach contents. Morphological traits were measured at the species level using data from the Global Ants Database, which provides standardized trait information across ant species [@parr2017].

## Data Analysis

All statistics and figures were generated in R Studio (version 1.1.442) running R (version 3.5.2).

```{r}
#| warning: false
#| message: false
#| label: load-pkgs
#| code-summary: "Packages"

library(Hmisc)
library(corrr)
library(corrplot)
library(bipartite)
library(sjPlot)
library(sjmisc)
library(sjlabelled)
library(sjtable2df)
library(caret)
library(MASS)
library(ggplot2)
library(emmeans)      ## estimated marginal means (least-squares means)
library(lmerTest)     ## p-values for lme4 models
library(lme4)         ## Mixed models 
library(lmtest)
library(MuMIn)
library(ggpubr)
library(nlme)
library(GGally)
library(PerformanceAnalytics)
library(psych)
library(knitr)
library(performance)
library(see)
library(viridis)       ## Viridis colour palette
library(car)
library(mvnTest)
library(rstatix)       ## different analyses (e.g., pairwise.t.test, Dunn's test)
library(FSA)           ## alternative for Dunn's test
library(repmis)
library(tidyverse)     ## the tidyverse
library(vegan)         ## Ordination, dissimilarity analyses
library(colorspace)    ## adjust colors
library(rcartocolor)   ## Carto palettes
library(ggforce)       ## sina plots
library(ggdist)        ## halfeye plots
library(ggridges)      ## ridgeline plots
library(ggbeeswarm)    ## beeswarm plots
library(gghalves)      ## off-set jitter
library(systemfonts)   ## custom fonts
library(kableExtra)
library(ggh4x)
library(ape)            
library(wesanderson)    ## Colour palette
library(grid)
library(png)
library(devtools)
library(pairwiseAdonis)
library(reshape2)       ## function melt
library(readr)          ## to export kable
library(rptR)           ## Repeatability
library(DescTools)
library(igraph)
library(remotes)
library(tinytable)
library(stats)
library(ggcorrplot)
library(glmmTMB)
library(DHARMa)
library(dietr)
library(FactoMineR)
library(factoextra)
library(modeldb)
library(ade4)
library(conflicted)
library(ggsankey)
library(raster)
library(circlize)
library(scico)
library(magrittr)

conflicts_prefer(dplyr::select)# Will prefer dplyr::select over any other package. 

conflicts_prefer(dplyr::filter)# Will prefer dplyr::filter over any other package. 

conflicts_prefer(base::attr)

conflicts_prefer(dplyr::summarize)

conflicts_prefer(dplyr::mutate)

conflicts_prefer(dplyr::rename)

conflicts_prefer(dplyr::summarise)

conflicts_prefer(base::union)

conflicts_prefer(magrittr::set_names)

```

All data sets used for this analyses can be accessed in: <https://drive.google.com/open?id=1d0iVHDFW4sX9hHQyasMv9oO-aWF1Ff_5&usp=drive_fs>

```{r}
#| warning: false
#| message: false
#| label: Data
#| code-summary: "data"
#| cache: false

#LOBSU Mac
feat_tab <- read.csv('/Users/camilorl/Library/CloudStorage//Shared drives/LOBSU Manuscripts/O. sylvatica vs. H. infraguttatus diet (Nora)/Submission 2-JAE/Code&Data/featureTableY_normalized.csv')

#LOBSU Mac 
alkadiver <- read.csv(
  "/Users/camilorl/Library/CloudStorage//Shared drives/LOBSU Manuscripts/O. sylvatica vs. H. infraguttatus diet (Nora)/Submission 2-JAE/Code&Data/alkadiver.csv")

#My Mac 
#alkadiver <- read.csv(
###  '/Users/camilorodriguezlopez/Library/CloudStorage//Shared drives/LOBSU Manuscripts/O. sylvatica vs. H. infraguttatus diet (Nora)/Submission 2-JAE/Code&Data/alkadiver.csv') # has compositional differences in skin alkaloid profiles between *O. sylvatica* populations and *H.infragutatus*, and altitude for every locality.

#LOBSU Mac 
L_antdiver <- read.csv("/Users/camilorl/Library/CloudStorage//Shared drives/LOBSU Manuscripts/O. sylvatica vs. H. infraguttatus diet (Nora)/Submission 2-JAE/Code&Data/L_antdiver.csv")

#My Mac 
#L_antdiver <- read.csv('/Users/camilorodriguezlopez/Library/CloudStorage//Shared drives/LOBSU Manuscripts/O. sylvatica vs. H. infraguttatus diet (Nora)/Submission 2-JAE/Code&Data/L_antdiver.csv')#has the abundance of all ant genera collected from the leaf litter and from frogs' stomach contents in every locality.

#LOBSU Mac 
diet <- read.csv(
  "/Users/camilorl/Library/CloudStorage//Shared drives/LOBSU Manuscripts/O. sylvatica vs. H. infraguttatus diet (Nora)/Submission 2-JAE/Code&Data/diet.csv")

#My Mac 
#diet <- read.csv(
###  '/Users/camilorodriguezlopez/Library/CloudStorage//Shared drives/LOBSU Manuscripts/O. sylvatica vs. H. infraguttatus diet (Nora)/Submission 2-JAE/Code&Data/diet.csv')#has the type and number of prey items found in the stomach contents of frog individuals from all populations.

#LOBSU Mac 
ant_traits <- read.csv("/Users/camilorl/Library/CloudStorage//Shared drives/LOBSU Manuscripts/O. sylvatica vs. H. infraguttatus diet (Nora)/Submission 2-JAE/Code&Data/ant_traits.csv")

#My Mac 
#ant_traits <- read.csv('/Users/camilorodriguezlopez/Library/CloudStorage//Shared drives/LOBSU Manuscripts/O. sylvatica vs. H. infraguttatus diet (Nora)/Submission 2-JAE/Code&Data/ant_traits.csv')#has 17 morphological traits of all ant genera collected from the leaf litter in every locality.


#LOBSU Mac
ant_all <- read.csv("/Users/camilorl/Library/CloudStorage//Shared drives/LOBSU Manuscripts/O. sylvatica vs. H. infraguttatus diet (Nora)/Submission 2-JAE/Code&Data/ant_all.csv")

#My Mac
#ant_all <- read.csv('/Users/camilorodriguezlopez/Library/CloudStorage//Shared drives/LOBSU Manuscripts/O. sylvatica vs. H. infraguttatus diet (Nora)/Submission 2-JAE/Code&Data/ant_all.csv')# has abundance of all ant species captured in pitfall and Winkler traps for every place.
```

### Alkaloid comparisons

We used a Kruskal-Wallis test to examine overall differences between *O. sylvatica* and *Hyloxalus infraguttatus* populations in summed toxicity across the 79 alkaloids in the 13 structural families. A pairwise Wilcoxon test was used to determine *O. sylvatica* population differences between the 13 structural families in both species, with p-values adjusted for multiple testing using the False Discovery Rate control (FDR).

```{r}
#| warning: false
#| message: false
#| output: false

#Kruskal-Wallis test to compare skin alkaloids accross populations
kw_test <- alkadiver %>% 
  select(-latitude, -longitude, -altitude) %>% 
  mutate(total = select(., contains(".")) %>%
           rowSums(na.rm = TRUE)) %>% 
  do(tidy(kruskal.test(.$total ~ .$population)))

###-------------------------##
#### NMDS for skin alkaloids ##
##-------------------------##

#the ordination was performed using Bray-Curtis dissimilarities
ord <- metaMDS((alkadiver %>% 
  select(where(is.numeric), -altitude, -latitude, -longitude)), 
  perm=9999, distance = "bray", 
               k = 2, autotransform = FALSE)

#the ordination was ran a second time starting at the previous best solution to ensure the stability and reliability of the results in the NMDS 
ord2<-metaMDS((alkadiver %>% 
  select(where(is.numeric), -altitude, -latitude, -longitude)), previous.best = TRUE)


#Include environmental variables 

### Create a vector with the "environmental variables" 
#For this I will use the coordinates of every locality and extract average historical temperature and precipitation, in addition to altitude, from World Clim.  

### Load average temperature and precipitation rasters from : https://www.worldclim.org/data/worldclim21.html 

### Adjust this path accordingly:

#My_Mac
#prec <- stack('/Users/camilorodriguezlopez/Library/CloudStorage//Shared drives/LOBSU Manuscripts/O. sylvatica vs. H. infraguttatus diet (Nora)/Submission 2-JAE/Code&Data/wc2.1_30s_prec_05.tif')  # layer for May
#temp <- stack('/Users/camilorodriguezlopez/Library/CloudStorage//Shared drives/LOBSU Manuscripts/O. sylvatica vs. H. infraguttatus diet (Nora)/Submission 2-JAE/Code&Data/wc2.1_30s_tavg_05.tif')  # layer for May

#LOBSU_Mac
prec <- stack('/Users/camilorl/Library/CloudStorage//Shared drives/LOBSU Manuscripts/O. sylvatica vs. H. infraguttatus diet (Nora)/Submission 2-JAE/Code&Data/wc2.1_30s_prec_05.tif')  # layer for May
temp <- stack('/Users/camilorl/Library/CloudStorage//Shared drives/LOBSU Manuscripts/O. sylvatica vs. H. infraguttatus diet (Nora)/Submission 2-JAE/Code&Data/wc2.1_30s_tavg_05.tif')  # layer for May

### Coordinates for every locality
coords <- alkadiver %>% 
  rename(lon = longitude , lat = latitude) %>% 
  select(lon, lat)

### Define a point (longitude, latitude)
coordinates(coords) <- ~lon + lat

### Extract temperature and precipitation for May (when the sampling was made) at the specified location
temp_value <- raster::extract(temp, coords)
prec_value <- raster::extract(prec, coords)

### Add location and altitude
env.vars <- alkadiver %>%
  select(altitude) %>% 
  bind_cols(data.frame(temp_value), data.frame(prec_value)) %>% 
  rename(Avtemp = wc2.1_30s_tavg_05, Avprec = wc2.1_30s_prec_05)

### fit environmental variables in the ordination
en <- envfit(ord2, env.vars, perm = 999) # We can see that altitude, latitude and longitude, are significantly correlated with the dimensions 


### fit alkaloids in the ordination to see their individual contribution
en.alka <- envfit(ord2, alkadiver %>%
  select(-latitude, -longitude, -altitude) %>%
  mutate(population = fct_relevel(population, "ceiba", "c_colon", "p_quito", "s_domingo", "la_mana", "H_infra")) %>%
  pivot_longer(cols = indo.137:pyrrz.526, names_to = "alkaloid", values_to = "abundance") %>%
  bind_cols(
    feat_tab %>%
      select(family, santos2016_source) %>%
      slice(rep(1:n(), times = 59))
  ) %>%
  select(frog_id,population,family, abundance) %>%
  pivot_wider(
    names_from = family,
    values_from = abundance,
    values_fn = sum,
    values_fill = 0
  ) %>%
    select(where(is.numeric), -frog_id, -population), perm = 999) 

sort(en.alka$vectors$r, decreasing = T) %>% 
  data.frame()

#PERMANOVA
permanova.output<-adonis2(alkadiver[,6:81]~alkadiver$population, 
                          permutations = 9999,method="bray")

#Pairwise adonis
pair.mod <-pairwise.adonis(alkadiver[,6:81],factors=alkadiver$population,
                           p.adjust = "BH")

```

### Frogs’ diet comparisons

We visualized interactions between arthropod prey taxa and poison frog species in a bipartite network based on the average abundance of arthropod prey for every frog population. We used the species specificity index (*ssi*), as implemented in the **specieslevel()** function of the ‘*bipartite*’ package, to quantify the variability in dietary interactions between arthropod prey taxa and poison frog species in a bipartite network. This index reflects the degree to which each frog species interacts unevenly across prey taxa, with values ranging from 0 (indicating low variability and generalist behavior) to 1 (indicating high variability and specialist behavior). Additionally, we used generalized linear mixed models (GLMM) to test for compositional differences of frog diet categories using the function **glmmTMB()** within the package ‘*glmmTMB*’ [@glmmTMB]. We used a negative binomial distribution appropriate for count data with overdispersion. We tested for diet differences by including species/population, prey type, and their interaction as main effects. Frog individual tags were included as a random variable to account for repeated sampling of prey categories within individuals. We computed estimated marginal means to test for pairwise comparisons between populations using the ‘emmeans’ package [@emmeans].

```{r}
#| warning: false
#| message: false
#| output: false

### to calculate the mean abundance of consumed prey for every frog population

sum_diet <- diet %>% # [-311,] remove the frog with 18 ants 
  group_by(population, prey) %>% 
  summarise(count_mean = mean(count)) %>%
  pivot_wider(id_cols = population, names_from = prey, 
              values_from = count_mean) %>% 
  data.matrix()


rownames(sum_diet) <- c("ceiba", "c.colon", "h.infragutatus", "mana",  
                         "p.quito", "s.domingo")

sum_diet <- sum_diet[,-c(1)]

### To calculate the specialization index
#dprime <- dfun(sum_diet, abuns=c(1,1,1,1,1))$dprime #I set hypothetical abundances of items in the leaf litter because we don't have the real information.
#However, even if I double the abundance of ants "abuns=c(2,1,1,1,1)", the specialization index is still higher for sylvatica and very low for infra.

#I now use the species specificity index, which which measures the variability in interaction strengths for a species, normalized to range from 0 (Low specificity - interactions are evenly distributed across all partners, i.e., generalist behavior) to 1 (High specificity - interactions are concentrated on a few partners, i.e., specialist behavior)

ssi <- specieslevel(sum_diet, index = "species specificity")

#To test for species differences (O. sylvatica vs. H. infraguttatus) in the number of prey items consumed

sd1 <- glmmTMB(count ~ population*prey,
               family=nbinom1(link="log"), data=diet)

anova.sd1 <- glmmTMB:::Anova.glmmTMB(sd1, typr="II")

#Check model fit by testing overdispersion 
disp.test <- function() {testDispersion(sd1)} #Remove "function() {}"
simulationOutputspp <- simulateResiduals(fittedModel = sd1, plot = F)
test.plots <- function() {plot(simulationOutputspp)} #Remove "function() {}"

#Calculate pairwise comparisons between prey items, within populations 
emm1 <- data.frame(emmeans(sd1, list(pairwise ~ population*prey), 
                           adjust = "tukey")$`emmeans of population, prey`)

emm1$prey <- factor(emm1$prey, levels = c("ants", "mites", "beetle", "other", "larvae"))

emm1$population <- factor(emm1$population, levels = c("ceiba", "puerto_quito","santo_domingo",  "cristobal_colon", "la_mana_Os", "la_mana_Hi"))

### To get rid of the very high values of puerto quito - other prey items
emm1$emmean[emm1$emmean < -10] <-  0
emm1$asymp.LCL[emm1$asymp.LCL < -10] <-  0
emm1$asymp.UCL[emm1$asymp.UCL > 10] <-  0
```

### Ant- vs mite -derive alkaloids

To infer potential dietary origins of frogs’ skin alkaloids, we cross-referenced the structural classes of alkaloids with the determined arthropod source reported by [@santos2016], which assigns compounds as derived from ants, mites, or both. We tested whether the proportion of ant-based alkaloids is greater than the proportion of mite-based alkaloids across frog populations using an anova followed by a Tukey post-hoc test for multiple comparisons. We visualized the proportion of ant- and mite-derived alkaloids using a chord diagram. Additionally, we tested whether total alkaloid abundance, as well as abundance within structural families, correlated with the number of ants and mites consumed across populations.

```{r}
#| warning: false
#| message: false

#### Test if the proportion of ant-based alkaloids is greater than the proportion of mite-based alkaloids across populations

antmitest <- alkadiver %>% 
  select(-latitude, -longitude, -altitude) %>% 
  mutate(total = select(., contains(".")) %>%
           rowSums(na.rm = TRUE)) %>%
  mutate(population = fct_relevel(population, "ceiba", "c_colon", "p_quito", "s_domingo", 
                                  "la_mana", "H_infra")) %>% 
  pivot_longer(cols = c(indo.137:pyrrz.526), 
               names_to = "alkaloid", values_to = "abundance") %>% 
  bind_cols(feat_tab %>% 
              select(family, santos2016_source) %>%
  slice(rep(1:n(), times = 59))) %>% 
  group_by(population, family, santos2016_source) %>% 
  summarize(abundance = sum(abundance)) %>% 
  mutate(family = factor(family, levels = c('HTX', '3,5-I', 'DHQ', 'Pyr', 
                                            '3,5-P', 'Lehm', 'Pip', '5,8-I', 
                                            '5,6,8-I', '1,4-Q', 'PTX', 
                                            '4,6-Q', 'aPTX'))) %>%
  group_by(population, santos2016_source) %>%
  summarise(total_abund = sum(abundance), .groups = "drop") %>%
  pivot_wider(names_from = santos2016_source, 
              values_from = total_abund, values_fill = 0) %>%
  mutate(total = Ants + Mites + Both,
         prop_ant = Ants / total,
         prop_mite = Mites / total,
         prop_both = Both / total) %>% 
  select(population, prop_ant, prop_mite, prop_both) %>% 
  pivot_longer(c(prop_ant:prop_both), names_to = "source", values_to = "proportion") %>%
  aov(proportion ~ source, data = .) %>%
  #plot()# to check residual normality
  #summary()
  TukeyHSD() %>%
  broom::tidy() %>% 
  kable(digits = 3, 
      table.attr = 'data-quarto-disable-processing="true"', "html",
      caption = "Summary of the results of Tukey post-hoc comparisons of an ANOVA in proportion of potential alkaloid arthropod source between frog populations") %>% 
  kable_classic(full_width = F, html_font = "Cambria") %>% 
  row_spec(0, bold = T)

### Correlation between #ants & # mites with alkaloids abundance 
cor_dietalk <- diet %>% 
  filter(prey %in% c("ants", "mites")) %>% 
  pivot_wider(names_from = "prey", values_from = "count", 
              id_cols = c(frog_id, population)) %>% 
  left_join(alkadiver %>% 
              select(-latitude, -longitude, -altitude) %>% 
              mutate(total = select(., contains(".")) %>%
                       rowSums(na.rm = TRUE)), by = "frog_id") %>% 
  na.omit() %>% 
  select(ants, mites, c(indo.137: total)) %>% 
  setNames(c(names(.)[1:2], feat_tab %>% 
               select(family) %>% 
               pull(), "total")) %>%
  {split.default(., names(.))} %>%                    # group columns by name
  map_dfc(~ rowMeans(as.data.frame(.), na.rm = TRUE)) %>% 
  corr.test() %>% 
  .$ci %>%
  rownames_to_column("correlations") %>%
  filter(str_detect(correlations, "mite|ant")) %>% 
  kable(digits = 3, 
      table.attr = 'data-quarto-disable-processing="true"', "html") %>% 
  kable_classic(full_width = F, html_font = "Cambria") %>% 
  row_spec(0, bold = T)

```

### Skin alkaloids vs. leaf litter ant communities along a geographical gradient

We looked at compositional differences in skin alkaloid profiles of *O. sylvatica* populations and *H. infraguttatus*, and their surrounding leaf litter ant communities in two separate non-metric multidimensional scaling (NMDS), using the function **metaMDS()** within the package '*vegan*' [@vegan]. Statistical differences between and within populations were assessed using a permutational multivariate analysis of variance (PERMANOVA) on Bray-Curtis dissimilarities. P-values for pairwise comparisons were adjusted using the function **pairwise.adonis()**. Additionally, we used the **envfit()** function to understand the influence of altitude, ambient temperature and precipitation on both alkaloid composition and leaf litter ant communities. These environmental variables were selected to capture spatial and temporal variation in ecological conditions that could influence ant availability and frogs’ foraging behavior. Ambient temperature and precipitation data was extracted from WorldClim 2.1, at a spatial resolution of 30 arc-seconds (\~1 km²) for each study site [@fick2017]. Similarly, we used envfit() to explore the contribution of specific alkaloid classes and ant genus to the ordination spaces. We used Procrustes analysis to test similarities between the NMDS ordinations of skin alkaloids and ant community composition across sites. The Procrustes correlation was calculated using the **procrustes()** function from the ‘*vegan*’ package, which aligns the two ordinations by scaling, rotating, and translating one configuration to best match the other. Statistical significance of the correlation was evaluated with 999 permutations using the **protest()** function.

```{r}
#| warning: false
#| message: false
#| output: false

##### Winkler ###
#NOTE: S. Domingo and P. Quito seem to have one outlier each when performing the ordination. I remove them.
ant_co <- (ant_all %>%
  filter(TRAP == "Winkler") %>%
  select(SITE, SPECIES, X., altitude, latitude, longitude) %>% # important to avoid duplicates
  group_by(SITE,SPECIES) %>% # important to avoid duplicates
  mutate(row = row_number()) %>% 
  pivot_wider(names_from = SPECIES, values_from = X.) %>%
  select(-row) %>% 
  replace(is.na(.), 0) %>% 
  mutate(SITE = factor(SITE)) %>% 
  ungroup() %>% 
  mutate(SITE = fct_relevel(SITE, "Mana", "Colon", "Santo Domingo", 
                            "Puerto Quito")) %>%
  arrange(as.integer(SITE)) %>% 
      as.data.frame())[-c(47,65),]
  

#the ordination was performed using Bray-Curtis dissimilarities
ord.co <- metaMDS(ant_co %>%
                     select(where(is.numeric), -altitude, 
                            -latitude, -longitude), 
                  perm=9999, distance = "bray",
                  k = 2, autotransform = FALSE)

  #the ordination was ran a second time starting at the previous best solution to ensure the stability and reliability of the results in the NMDS 
ord.co2<-metaMDS(ant_co %>%
                     select(where(is.numeric), -altitude, 
                            -latitude, -longitude), 
                 previous.best = TRUE)

#Include environmental variables - altitude, latitude and longitude: 
### Create a vector with the "environmental variables" 

coords.2 <- ant_co %>% 
  rename(lon = longitude , lat = latitude) %>% 
  select(lon, lat)

### Define a point (longitude, latitude)
coordinates(coords.2) <- ~lon + lat

### Extract temperature and precipitation for May at the specified location
temp_value.2 <- raster::extract(temp, coords.2)
prec_value.2 <- raster::extract(prec, coords.2)

### Add location and altitude

env.vars.co <- ant_co %>%
  bind_cols(data.frame(temp_value.2), data.frame(prec_value.2)) %>% 
  rename(Avtemp = wc2.1_30s_tavg_05, Avprec = wc2.1_30s_prec_05) %>% 
  select(altitude, Avtemp, Avprec) 

en.co <- envfit(ord.co2, env.vars.co, perm = 999) # We can see that altitude, latitude and longitude, are significantly correlated with the dimensions 

en.co.ants <- envfit(ord.co2, ant_co %>%
         select(where(is.numeric), -altitude,
                -latitude, -longitude), perm = 999)

sort(en.co.ants$vectors$r, decreasing = T) %>% 
  data.frame() 


en.co$vectors$arrows %>%
  data.frame() %>%
  mutate(r2 = en.co$vectors$r) %>% 
  mutate("p-value" = en.co$vectors$pvals) %>% 
  kable(digits = 3,caption = "") %>% 
  kable_classic(full_width = F, html_font = "Cambria") %>% 
  row_spec(0, bold = T) 

#PERMANOVA
permanova.ants<-adonis2((ant_co %>% 
                          select(-SITE,-altitude, -latitude, -longitude)) ~ant_co$SITE, 
                          permutations = 9999,method="bray")

#Pairwise adonis
pair.per.ants <-pairwise.adonis((ant_co %>% select(-SITE,-altitude, -latitude, -longitude)),
                                 factors=ant_co$SITE, p.adjust = "BH")

```

### Stomach vs. leaf litter ant communities and frogs’ preference for ant genera

To test for within and between species differences in ant abundance between leaf litter and stomach contents, we employed a negative binomial generalized linear model using the **glm.nb()** function from the ‘*MASS*’ package [@MASS]. The model used total ant abundance per sample as the response variable, with group (stomach vs. leaf) and population as predictors. Pairwise comparisons were conducted using estimated marginal means. To examine if frogs show selectivity for consuming specific ant genera, we calculated a linear selectivity index by subtracting the relative abundance of ants found on the leaf litter from those found in frog stomach contents [@strauss1979]. The interpretation of these selectivity values was guided by the methodology in [@mcelroy_ant_2019], which involves generating a null distribution of selectivity values via simulation for each ant species. By comparing the observed selectivity values to this null distribution, we classified the ants as ‘selected’ if the values were above, ‘neutral’ if they were within, and ‘avoided’ if they were below the null distribution, thereby delivering a statistically robust assessment of the frogs’ selective foraging behaviors.

```{r}
#| warning: false
#| message: false


### To calculate the total ant abundance per sample only in Winkler sacs
sum_abun <-  L_antdiver %>% 
  rowwise(Trap, group, population) %>% 
  summarise(abundance = sum(c_across(Anochetus:Wasmannia))) %>% 
  filter(!(Trap == "Pitfall"))

#To check normality
shapiro.ants <- shapiro.test(sum_abun$abundance)

Histant <- sum_abun %>% 
  ggplot(aes(x=abundance)) +
  geom_histogram(aes(x=abundance, y=..density..), 
                 bins=5,col="black", fill="black",alpha = 0.15) +
  theme_bw(14) + 
  theme(panel.grid.major = element_blank(), 
        panel.grid.minor = element_blank(), 
        axis.text.x = NULL, legend.position = "none") +
  ylab("Density") + 
  theme(axis.text=element_text(size=9),
        axis.title=element_text(size=11)) +
  annotate("text", x = 250, y = 0.009, label = "Shapiro-Wilk test", 
           size = 3.5) +
  annotate("text", x = 250, y = 0.007, label = "p-value = <0.001", 
           size = 3.5) +
  annotate("text", x = 250, y = 0.005, label = "W = 0.83", 
           size = 3.5)

###### Ant abundance comparison between leafs and stomachs
### We used the data base "L_antdiver"

### Negative binomial logistic regression
sumabun.mod <- glm.nb(abundance ~ group*population, data = sum_abun)
sumabun.aov <- anova(sumabun.mod)

### Estimated marginal means
sumabun.emmeans <- emmeans(sumabun.mod, list(pairwise ~ group*population), 
                           adjust = "tukey")

### Calculate linear selectivity based on (McElroy & Donoso, 2019)

### Function to calculate null distribution 
custom_loop <- function(data) {
  for (i in 1:1000){                                          # number of loops
    j <- sample(2000:5000,1)                                  # number of ants sampled
    randomdraw <- sample(data$Genus,                             # draw species from env. proportion with replacement
                         size = j,
                        replace = TRUE, 
                        prob = data$leafprop)                      
    df_randomdraw <- as.data.frame(table(randomdraw))         # dataframe of species counts
    preyprop_sim <- df_randomdraw$Freq/j                      # vector of simulated prey species proportions
    Linear_sim <- as.data.frame(preyprop_sim - data$leafprop)       # simulated LinearSelectivity by subtracting REAL env_prop from SIMULATEd prey_prop + make dataframe
    colnames(Linear_sim) <- paste0("sim",i)                   # rename column from LinearS_sim --> sim1, sim2, sim3,...sim1000
  
  # add the simulated dataset to the dataframe with species, LinearS, sim1, sim2,...etc....  
    data <- cbind(data,Linear_sim) 
  }
  return(data)
}


### Calculate linear selectivity and null distribution using bootstrap with 1000 iterations 
L_simulations <- L_antdiver %>%
  group_by(group, population) %>% 
  select(where(is.numeric)) %>% 
  summarise_at(vars(Anochetus:Wasmannia), sum) %>%
  rowwise() %>% 
  mutate(across(Anochetus:Wasmannia, ~./sum(c_across(Anochetus:Wasmannia)))) %>%
  ungroup() %>% 
  melt() %>% 
  pivot_wider(names_from = group, values_from = value) %>% 
  rename(frogprop = frog, leafprop = leaf, Genus = variable) %>% 
  mutate(Linear = frogprop - leafprop) %>%
  group_by(population) %>% 
  do(custom_loop(.))

### melt all simulated linear selectivity for plotting null distribution 
Lsel_sim <- L_simulations %>% 
  select(-Linear) %>% 
  melt(id.vars = c("population", "Genus"), measure.vars = paste0("sim",c(1:1000))) %>% 
  group_by(population) %>% 
  arrange(desc(value)) %>%
  mutate(Genus = reorder(Genus, value)) 

### melt observed linear selectivity and assign category based on null distribution
Lsel_obs <- L_simulations %>% 
  melt(id.vars = c("population", "Genus"), measure.vars = c("Linear")) %>% 
  group_by(population) %>% # group by population to rearrange values
  arrange(desc(value)) %>%
  mutate(Genus = reorder(Genus, value)) %>%
  rename(Linear = value) %>% # arrange values by genus
  group_by(population, Genus) %>% 
  mutate(Cat.Linear = case_when(
    Linear > max(Lsel_sim$value) ~ "preferred", # selected are all values falling above null distribution   
    Linear >= min(Lsel_sim$value) & Linear <= max(Lsel_sim$value) ~ "neutral", # neutral are all values falling within null distribution
    Linear < min(Lsel_sim$value) ~ "avoid")) # avoid are all values falling under null distribution

#to arrange genus based on linear selectivity


Lsel_sim <- Lsel_sim %>%
  mutate(Genus = factor(Genus, levels = c("Hypoponera",
                                          "Carebara", "Rogeria",
                                          "Myrmicocrypta", "Anochetus",
                                          "Apterostigma", "Gnamptogenys",
                                          "Hylomyrma", "Octostruma",
                                          "Sericomyrmex",
                                          "Wasmannia","Pheidole", 
                                          "Cyphomyrmex",
                                          "Crematogaster", "Trachymyrmex",
                                          "Strumigenys", "Solenopsis")))

Lsel_obs <- Lsel_obs %>%
  mutate(Genus = factor(Genus, levels = c("Hypoponera",
                                          "Carebara", "Rogeria",
                                          "Myrmicocrypta", "Anochetus",
                                          "Apterostigma", "Gnamptogenys",
                                          "Hylomyrma", "Octostruma",
                                          "Sericomyrmex",
                                          "Wasmannia","Pheidole", 
                                          "Cyphomyrmex",
                                          "Crematogaster", "Trachymyrmex",
                                          "Strumigenys", "Solenopsis")))

```

### Ant morphology vs frog selectivity

To determine whether the morphological and life history traits of ant species in the leaf litter influences frog selectivity, we performed a Principal Component Analysis (PCA). The PCA aimed at reducing the dimensionality of 17 morphological and life history traits across 17 ant genera found both in the leaf litter and in *O. sylvatica* stomachs. To build the PCA, we used the function **dudi.pca()** from the ‘*ade4*’ package [@ade4]. Scores of components with eigenvalues higher than 1 were selected and used as response variables in a pairwise Wilcoxon test to determine differences between selectivity categories for ants in all diablito populations. P-values were adjusted for multiple testing using the False Discovery Rate control (FDR).

```{r}
#| warning: false
#| message: false
#| fig-cap: "Eigenvalues of the first 5 principal components. The first three components have eigenvalues higher than one"
#| fig-height: 2
#| fig-width: 3
#| column: margin

### Perform Principal Components Analysis for ant traits 
PCAtraits <- na.omit(ant_traits) %>% 
  filter(Genus %in% Lsel_obs$Genus) %>%  
  select(where(is.numeric)) %>% # retain only numeric columns
  dudi.pca(scale = T, center = T, scannf = FALSE, nf = 3) # we keep the first three components as they explain most of the variance (see scree plot)


### Plot eigenvalues
eigenvalues <- data.frame(eigenvalues = PCAtraits$eig) %>% 
  mutate("Component" = paste0("PC", 1:length(eigenvalues))) %>% 
  filter(Component %in% c("PC1", "PC2", "PC3", "PC4", "PC5")) %>%
  arrange(eigenvalues) %>%
  mutate(Component = reorder(Component, eigenvalues, decreasing = T)) %>%  
  ggplot(aes(x = Component, y = eigenvalues)) +
  geom_bar(stat = "identity", col = "black", fill = "darkgoldenrod", size = 0.3) + 
  geom_hline(yintercept = 1, lty = 2, col = "black") +
  theme_bw() + 
  theme(panel.grid.major = element_blank(),
        panel.grid.minor = element_blank(),
        axis.text.x = NULL, 
        plot.title = element_blank())

eigenvalues

```

# Results

## Alkaloids differ between species and across diablito frog populations

Skin extracts consisted of 79 alkaloids. The summed amount of alkaloids varied across species and populations (Kruskal-Wallis; X2(5) = 41.542, p \< 0.001; @fig-2 A), with *O. sylvatica* having more alkaloids than *H. infraguttatus* (*H. infraguttatus vs. all other O. sylvatica populations*, p \< 0.001; @tbl-1). Within *O. sylvatica*, the Ceiba population had less toxins than all others (p \< 0.001; @tbl-1), while frogs from Santo Domingo had on average the highest alkaloid load (@tbl-1).

```{r}
#| warning: false
#| message: false
#| tbl-cap: "P-values of pairwise Wilcoxon test on differences in summed alkaloids between frog populations"
#| tbl-cap-location: top
#| label: tbl-1

#Pairwise Wilcoxon test
pwise.test <- alkadiver %>% 
  select(-latitude, -longitude, -altitude) %>% 
  mutate(total = select(., contains(".")) %>%
           rowSums(na.rm = TRUE)) %>%
  mutate(population = fct_relevel(population, "ceiba", "c_colon", "p_quito", "s_domingo", 
                                  "la_mana", "H_infra")) %>% 
  do(tidy(pairwise.wilcox.test(.$total, .$population,
                               p.adjust.method = "BH")$p.value)) %>% 
  mutate(rowName = c("C. Colón", "P. Quito", "S. Domingo", "La Maná", "H. infraguttatus")) %>% 
  column_to_rownames(var = "rowName") %>% 
  data.frame(.$x) %>% 
  select(-x) %>% 
  rename("C. colón" = c_colon, "Ceiba" = ceiba, "S. Domingo" = s_domingo, 
         "La Maná" = la_mana, "P. Quito" = p_quito)
  

### Results of pairwise comparisons

options(knitr.kable.NA = "-")# to remove "NA's" from the table

T1 <- pwise.test %>% 
  mutate(across(where(is.numeric), round, 3)) %>% 
  mutate(across(where(is.numeric), ~ ifelse(. < 0.001, "<0.001", as.character(.)))) %>% 
  kable(digits = 2, 
      table.attr = 'data-quarto-disable-processing="true"', "html",
      caption = "Table S1:P-values of pairwise Wilcoxon test on differences in summed alkaloids between frog populations") %>% 
  kable_classic(full_width = F, html_font = "Cambria") %>% 
  row_spec(0, italic = T, bold = T) %>% 
  column_spec(1, italic = T, bold = T)

T1

```

[Table 1](https://drive.google.com/open?id=1M1z2h2whc_-izPeJOMjDzu4XLhZP1gLt&usp=drive_fs){.external target="_blank"}

```{r fig2}
#| echo: false
#| fig-show: "hold"
#| fig-cap: "**Alkaloid profiles differ between *Oophaga sylvatica* populations.** **(A)** The summed relative abundances of 79 alkaloids are visualized as box plots, where each dot represents a frog. The y axis is square-root transformed for visual clarity and summed alkaloids are expressed in arbitrary units (a.u.). Populations are represented with different colors. *Hyloxalus infraguttatus*, collected near the town of La Maná, is shown on the far right. Groups not connected by the same letter are significantly different (Table 1). **(B)** Non-metric multidimensional scaling (NMDS) biplot based on Bray-Curtis dissimilarities show alkaloid profile differences between populations based on the relative abundances of 79 unique alkaloids (Stress = 0.156). Each dot represents a single frog’s alkaloid profile Ellipses represent 95% confidence intervals. Environmental factors (black arrows) and alkaloids (blue arrows) in correlation with the ordination are shown. **(C)** Bubble - heatmap shows percent of all summed alkaloids grouped by alkaloid structural class and population. Histrionicotoxins (HTX); 5,8-disubstituted indolizidines (5,8-I); 3,5-disubstituted indolizidine (3,5-I); Decahydroquinoline (DHQ); 5,6,8-trisubstituted indolizidines (5,6,8-I); Pyrrolidine (Pyr); 3,5-disubstituted pyrrolizidine (3,5-P); Lehmizidine (Lehm); 1,4-disubstituted quinolizidine (1,4-Q); Piperidine (Pip); Pumiliotoxin (PTX); Allopumilliotoxin (aPTX); 4,6-disubstituted quinolizidine (4,6-Q)."
#| cap-location: margin
#| label: fig-2
#| fig-height: 5
#| fig-align: center

grid.raster(readPNG('/Users/camilorl/Library/CloudStorage//Shared drives/LOBSU Manuscripts/O. sylvatica vs. H. infraguttatus diet (Nora)/Submission 2-JAE/Reviews/Fig2_neu.png')) 
```

[Figure 2.png](https://drive.google.com/open?id=1wVHeGUraM4OIw7SmkQ0PHNPpPkhignve&usp=drive_fs){.external target="_blank"}

We next visualized overall alkaloid compositional differences across *O. sylvatica* populations and *H. infraguttatus* using an NMDS (@fig-2 B). The NMDS suggested a two dimensional solution (stress = 0.156) and showed distinct clusters of alkaloid composition. The abundance of all 79 alkaloids varied significantly across groups (PERMANOVA, F(4) = 10.178, p \< 0.001). A post-hoc pairwise comparison indicated significant differences between all possible population pairs (@tbl-2; @fig-2 B), suggesting each group has a unique alkaloid profile. Fitting environmental variables into the NMDS indicated that altitude (r² = 0.72, p = 0.001) and temperature (r² = 0.68, p = 0.001) significantly influenced alkaloid composition across the geographical gradient (@fig-2 B). Given their strong inverse correlation and closely aligned vectors in NMDS space, we report both as reflecting a shared environmental gradient.

Skin alkaloids fell into one of 13 structural families: Histrionicotoxins (HTX); 5,8-disubstituted indolizidines (5,8-I); 3,5-disubstituted indolizidine (3,5-I); Decahydroquinoline (DHQ); 5,6,8-trisubstituted indolizidines (5,6,8-I); Pyrrolidine (Pyr); 3,5-disubstituted pyrrolizidine (3,5-P); Lehmizidine (Lehm); 1,4-disubstituted quinolizidine (1,4-Q); Piperidine (Pip); Pumiliotoxin (PTX); Allopumilliotoxin (aPTX); 4,6-disubstituted quinolizidine (4,6-Q). When examined more closely, histrionicotoxins, decahydroquinolines and indolizidines were the most abundant alkaloid classes relative to the total alkaloid content in both H. infraguttatus and all O. sylvatica populations sampled (@fig-2 C). Several indolizidines, including 5,6,8-I (r² = 0.54, p = 0.001), 5,8-I (r² = 0.54, p = 0.001) & 3,5-I (r² = 0.46, p = 0.001) contributed significantly to alkaloid ordination (@fig-2 B).

```{r}
#| warning: false
#| message: false
#| tbl-cap: "Summary of the results of pairwise comparisons of a PERMANOVA in alkaloid composition between *O. sylvatica* populations"
#| tbl-cap-location: top
#| label: tbl-2

colnames(pair.mod) <- c("Contrasts", "df", "Sums of sq.", "F", 
                        "R2","p-value", "p.adjusted", "Sig.")

T2 <- pair.mod %>%
  select(Contrasts, df, F, R2, p.adjusted) %>% 
  mutate(Contrasts = c("La Maná vs. Ceiba", "La Maná vs. C. Colón", 
                   "La Maná vs. S. Domingo", "La Maná vs.P. Quito",
                   "La Maná vs. H. infraguttatus", "Ceiba vs. C. Colón", 
                   "Ceiba vs. S. Domingo", "Ceiba vs. P. Quito", 
                   "Ceiba vs. H. infraguttatus", "C. Colón vs. S. Domingo", 
                   "C. Colón vs. P. Quito", "C. Colón vs. H.infraguttatus",
                   "S. Domingo vs. P. Quito", "S. Domingo vs. H. infraguttatus",
                   "P. Quito vs. H. infraguttatus")) %>% 
  kable(digits = 3, 
      table.attr = 'data-quarto-disable-processing="true"', "html",
      caption = "Table S2: Summary of the results of pairwise comparisons of a PERMANOVA in alkaloid composition between frog populations") %>% 
  kable_classic(full_width = F, html_font = "Cambria") %>% 
  row_spec(0, bold = T) 

T2
```

[Table 2](https://drive.google.com/open?id=1SeyK8a8lDdn4KqyvY4ouAEG3wtJsgJ7R&usp=drive_fs){.external target="_blank"}

```{r}
#| warning: false
#| message: false

#Boxplots 
f2a <- alkadiver %>% 
  select(-latitude, -longitude, -altitude) %>% 
  mutate(total = select(., contains(".")) %>%
           rowSums(na.rm = TRUE)) %>% 
  mutate(population = factor(population, 
                levels = c("ceiba", "c_colon", "p_quito",
                           "s_domingo", "la_mana", "ala_manaHI"))) %>% 
  ggplot(aes(y=total/10000000, x=population, fill=population)) + 
  geom_boxplot(outlier.shape = NA) + 
  geom_jitter(position=position_jitterdodge(0.1), shape = 21, 
              size = 4, alpha=0.6) +
  theme_classic(20) + 
  ylab("summed alkaloids (a.u.)") +
  scale_fill_manual(values=c("#d16b54", "#e8c95d", "#433447", "#b9b09f",
                             "#a9d8c8", "#ffffff"),
                    name = "populations", 
                    labels = c("Ceiba", "C. colon", "P. Quito", 
                               "S. Domingo", "La Maná", "H. infragutatus")) + 
  scale_y_sqrt() + 
  scale_y_continuous(trans = 'sqrt') +
  theme(axis.title.x = element_blank(), 
        axis.text.x = element_blank(), axis.title = element_text(size=15),
        plot.title = element_text(size=28,hjust=0.5),
        legend.position = "none")


### To make the NMDS plot in ggplot, we extract the dimension scores from the NMDS and add them to the main data
NMDSdims <- alkadiver %>% 
  mutate(NMDS1 = scores(ord2)$sites[,1], NMDS2 = scores(ord2)$sites[,2]) %>%
  data.frame()

### to extract the coordinates for the vectors of the environmental variables
en_coord <-  as.data.frame(scores(en, "vectors"))

en_alk_coord <-  as.data.frame(scores(en.alka, "vectors"))[c(1,4,6),]

### The contribution of every environmental variable
env.alka <- en$vectors$arrows %>%
  data.frame() %>%
  mutate(r2 = en$vectors$r) %>% 
  mutate("p-value" = en$vectors$pvals) %>% 
  kable(digits = 3, 
      table.attr = 'data-quarto-disable-processing="true"', "html") %>% 
  kable_classic(full_width = F, html_font = "Cambria") %>% 
  row_spec(0, bold = T)


#and the plot
f2b <- ggplot(NMDSdims, aes(x = NMDS1, y = NMDS2)) + 
  stat_ellipse(aes(colour = population, fill = population), 
               geom = "polygon", level = 0.95, alpha = 0.2, type = "t") +
  geom_point(aes(fill = population), pch = 21, size = 4, alpha = 0.5) + 
  scale_colour_manual(values = c("#e8c95d", "#d16b54", "gray90", "#a9d8c8", "#433447", "#b9b09f")) + 
  scale_fill_manual(values = c("#e8c95d", "#d16b54", "gray90", "#a9d8c8", "#433447", "#b9b09f")) + 
  geom_segment(aes(x = 0, y = 0, xend = NMDS1, yend = NMDS2), 
               data = en_coord, linewidth = 1, alpha = 0.5, colour = "grey30",
               arrow = arrow(type = "open", length = unit(0.1, "inches"))) + 
  geom_text(data = en_coord, aes(x = NMDS1-0.1, y = NMDS2), colour = "grey30", 
            fontface = "bold", label = row.names(en_coord)) +
  geom_segment(aes(x = 0, y = 0, xend = NMDS1, yend = NMDS2), 
               data = en_alk_coord, linewidth = 1, alpha = 0.5, colour = "red",
               arrow = arrow(type = "open", length = unit(0.1, "inches"))) + 
  geom_text(data = en_alk_coord, aes(x = NMDS1-0.1, y = NMDS2), colour = "red", 
            fontface = "bold", label = row.names(en_alk_coord)) +
  theme_bw() +
  theme(panel.grid.major = element_blank(), panel.grid.minor = element_blank(),
        axis.text.x = NULL, axis.text.y = NULL, legend.position = "none")


#Heatmap to visualize percentage of summed alkaloids grouped by alkaloid class and population

### To create a palette for a gradient color
pal <- wes_palette("Zissou1", 100, type = "continuous")

#By structural family
f2c <- alkadiver %>% 
  select(-latitude, -longitude, -altitude) %>% 
  #mutate(total = select(., contains(".")) %>%
  #         rowSums(na.rm = TRUE)) %>%
  mutate(population = fct_relevel(population, "ceiba", "c_colon", "p_quito", "s_domingo", 
                                  "la_mana", "H_infra")) %>% 
  pivot_longer(cols = c(indo.137:pyrrz.526), 
               names_to = "alkaloid", values_to = "abundance") %>% 
  bind_cols(feat_tab %>% 
              select(family, santos2016_source) %>%
  slice(rep(1:n(), times = 59))) %>% 
  group_by(family, population) %>% 
  summarise(sum.str.alk = sum(abundance)) %>% 
  group_by(population) %>% 
  mutate(percent = (sum.str.alk / sum(sum.str.alk))*100) %>% 
  mutate(family = factor(family, levels = c('HTX', '5,8-I', '3,5-I', 'DHQ', 
                                            '5,6,8-I', 'Pyr', '3,5-P', 'Lehm',
                                            '1,4-Q', 'Pip', 'PTX', '4,6-Q', 
                                            'aPTX'))) %>% 
  mutate(population = factor(population, levels = c("H_infra", "la_mana", "s_domingo", 
                                                    "p_quito", "c_colon", 
                                                    "ceiba"))) %>% 
  ggplot(aes(x = population, y = family, colour = percent)) + 
  geom_point(aes(size = percent, colour = percent)) +
  scale_color_gradientn(colours = pal) +
  theme_bw() + coord_flip() +
  theme(axis.text.x = element_text(angle = 45, hjust = 1, vjust = 1),
        legend.text.position = "right")


#By superfamily
#f2c <- alkadiver %>%  # to get the sum of alkaloids per individual
###  select(-altitude) %>%
###  melt() %>% 
###  mutate(toxin = case_when(grepl("indo", variable) ~ "indolizidine", 
 #                          grepl("allo", variable) ~ "allopumiliotoxin",
#                           grepl("quino", variable) ~ "quinolizidine", 
#                           grepl("pyrrz", variable) ~ "pyrrolizidine",
#                           grepl("deca", variable) ~ "decahydroquinoline", 
#                           grepl("hist", variable) ~ "histrionicotoxin",
#                           grepl("lehm", variable) ~ "lehmizidine", 
#                           grepl("piper", variable) ~ "piperidine", 
#                           grepl("pumi", variable) ~ "pumiliotoxin", 
#                           grepl("pyrro", variable) ~ "pyrrolidine")) %>%
###  select(-variable) %>% 
###  group_by(toxin, population) %>% 
###  summarise(sum.alk = sum(value)) %>% 
###  group_by(population) %>% 
###  mutate(percent = (sum.alk / sum(sum.alk))*100) %>% 
###  mutate(toxin = factor(toxin, levels = c('histrionicotoxin', 'indolizidine',
#                                          'decahydroquinoline', 'pumiliotoxin',
#                                          'pyrrolidine', 'pyrrolizidine', 
#                                          'quinolizidine', 'piperidine', 
#                                          'allopumiliotoxin', 'lehmizidine'))) %>% 
###  mutate(population = factor(population, levels = c("H_infra", "la_mana", "s_domingo", 
#                                                    "p_quito", "c_colon", 
#                                                    "ceiba"))) %>% 
###  ggplot(aes(x = population, y = toxin, colour = percent)) + 
###  geom_point(aes(size = percent, colour = percent)) +
###  scale_color_gradientn(colours = pal) +
###  theme_bw() + coord_flip() +
###  theme(axis.text.x = element_text(angle = 45, hjust = 1, vjust = 1))

```

## Defended frogs consumed more ants relative to other prey types and compared to the diet of the undefended species

We found that the number of prey consumed in different categories differ significantly across populations and between species (GLMM, population x prey type: X2(20) = 83.59, p \< 0.001; @fig-3 A & B). Post hoc pairwise comparisons and the species selectivity index (*ssi*) showed that all *O. sylvatica* populations consumed significantly more ants than other prey categories (x̅ = 75%; emmeans (ants vs. all prey): p-value = \<0.001; ssirange= 0.66 - 0.85; @fig-3 B, @tbl-3), whereas *H. infraguttatus* showed a more generalist dietary pattern, consuming a smaller but diverse array of arthropods including ants (45%), beetles (14%) and ‘other’ arthropods (25%; emmeans (all prey comparisons): p-value = \>0.05; ssi = 0.27; @fig-3 B, @tbl-3). It is worth noting that only one *H. infraguttatus* had 18 ants in the stomach, which accounts for nearly half of the total consumed for this species in our data set. When removing this individual, ants made up 36% of the total diet, followed by ‘other’ arthropods (29.6%) and beetles (16.5%).

```{r}
#| warning: false
#| message: false
#| output: false


### to visualize the bipartite interaction matrix:

### First, I formatted the diet data to make a sankey plot to visualize connections between preys and populations  
links <- data.frame(sum_diet) %>% 
  rownames_to_column(var = "target") %>% 
  pivot_longer(cols = c(ants, beetle, larvae, mites, other), names_to = "source", values_to = "value") %>% 
  mutate(target = case_when(grepl("ceiba", target) ~ 5, grepl("c.colon", target) ~ 9, 
                            grepl("h.infragutatus", target) ~ 10, grepl("mana", target) ~ 6, 
                            grepl("p.quito", target) ~ 7, grepl("s.domingo", target) ~ 8)) %>% 
  mutate(source = case_when(grepl("ants", source) ~ 0, grepl("beetle", source) ~ 2, 
                            grepl("larvae", source) ~ 3, grepl("mites", source) ~ 1, 
                            grepl("other", source) ~ 4)) %>% 
  data.frame() %>% 
  round() %>% 
  mutate(perc = value+1) %>% #add1 so the 0 are not 0
  arrange(source) %>% 
  select(source, target, value, perc)

#Here I repeated the number of rows according to the number of prey items per category per population
links_rep <- links[rep(row.names(links), times = links$perc), ]

### the Sankey plot
fig3a <- links_rep %>% 
  select(source, target) %>% 
  make_long(source, target) %>% 
  arrange(desc(node)) %>% 
  ggplot(aes(x = x, 
               next_x = next_x, 
               node = node, 
               next_node = next_node,
               fill = factor(node))) +
  geom_sankey(alpha=0.4, col = "black", lwd = 0.03) +
  theme_sankey(base_size = 16) +
  scale_fill_manual(values= c("#3E63A6", "#6AB897", "#A4BB7A","#E5C049", "#FDE725FF", 
                              rep("black",6))) + 
  theme(legend.position = "none")

##Alternative Chord diagram

sumdiet2 <- t(sum_diet)[ c(3,5,2,4,1),c(1,5,6,2,4,3)] 

chord.plot <- chordDiagram(sumdiet2, grid.col = c("#E5C049", "#FDE725FF", "#A4BB7A", "#6AB897", "#3E63A6",    
                                    rep("black",6))) 


### Plot of emmeans with 95% CIs 
fig3b <- ggplot(emm1, aes(prey, emmean, color = prey))  +
  geom_hline(yintercept = 0, linetype = 3, col = "grey") +
  geom_linerange(aes(ymin=asymp.LCL, ymax=asymp.UCL), linewidth=4,show.legend = F, alpha = 0.6) +
  geom_point(col = "black", size = 1) +
  facet_grid(.~population, scales = "free") +
  theme_bw() +
  theme(panel.grid.major = element_blank(),
                                    panel.grid.minor = element_blank(),
                                    axis.text.x = NULL, legend.position = "none") + 
  scale_color_manual(values=c("#3E63A6","#6AB897", "#A4BB7A", "#E5C049",  "#FDE725FF"))

```

```{r}
#| warning: false
#| message: false
#| tbl-cap: "Summarized results of estimated marginal means between frog species and prey items. P-values were adjusted using Tukey’s method"
#| tbl-cap-location: top
#| label: tbl-3


t3 <- data.frame(emmeans(sd1, list(pairwise ~ population*prey), 
                           adjust = "tukey")$`pairwise differences of population, prey`)[c(6,12,18,24,35,41,47,53,90,96,102,108,116,122,128,134,141,147,153,159,63,69,75,81),]# Manually choose the contrasts per population

T3 <- t3 %>%
  rename(Contrasts = X1) %>% 
  select(Contrasts, estimate, SE, p.value) %>% 
  remove_rownames() %>% 
  mutate(p.value = c(rep("<0.001",20), rep(">0.05", 4))) %>% #to read easily p.values
  mutate(Contrasts = rep(c("Ants vs. Beetles", "Ants vs. Larvae", 
                   "Ants vs. Mites", "Ants vs. Other"), 6)) %>% 
  kable(digits = 3, 
      table.attr = 'data-quarto-disable-processing="true"', "html",
      caption = "Table S3: Summarized results of estimated marginal means between frog species and prey items. P-values were adjusted using Tukey’s method") %>% 
  kable_classic(full_width = F, html_font = "Cambria") %>% 
  row_spec(0, bold = T) %>% 
  pack_rows("Ceiba",1,4) %>% 
  pack_rows("C. Colón",5,8) %>%
  pack_rows("La Maná",9,12) %>%
  pack_rows("P. Quito",13,16) %>% 
  pack_rows("S. Domingo",17,20) %>%
  pack_rows("H. infraguttatus",21,24)

T3
```

[Table 3](https://drive.google.com/open?id=1trtw6COpQQkHGcUJHPf9u9sUhg3xbkvl&usp=drive_fs){.external target="_blank"}

##Frogs’ alkaloid composition suggests a predominantly ant-based diet

We found that across all frog populations, including the "non-toxic" species H. infraguttatus, ant-derived alkaloids appeared to represent the most abundant component of their chemical profile, while a smaller but important contribution was attributed to mite-derived alkaloids (@fig-3 C). Tukey’s post hoc test revealed that the proportion of ant-derived alkaloids was significantly higher than mite-derived alkaloids (p\<0.001). Neither total alkaloid abundance nor abundance within structural families showed a significant correlation with the number of ants or mites consumed across populations (All p-values \> 0.05; @tbl-4).

```{r}
#| warning: false
#| message: false
#| output: false


#Visualization
#Make subset matrices for every population
allpop.chord <- alkadiver %>% 
  select(-latitude, -longitude, -altitude) %>% 
  mutate(total = select(., contains(".")) %>%
           rowSums(na.rm = TRUE)) %>%
  mutate(population = fct_relevel(population, "ceiba", "c_colon", "p_quito", "s_domingo", 
                                  "la_mana", "H_infra")) %>% 
  pivot_longer(cols = c(indo.137:pyrrz.526), 
               names_to = "alkaloid", values_to = "abundance") %>% 
  bind_cols(feat_tab %>% 
              select(family, santos2016_source) %>%
  slice(rep(1:n(), times = 59))) %>% 
  group_by(population, family, santos2016_source) %>% 
  summarize(abundance = sum(abundance)) %>% 
  mutate(family = factor(family, levels = c('HTX', 'DHQ', '5,8-I', 'Pyr', 
                                            '3,5-P', 'Lehm', 'Pip', '3,5-I',  
                                            '5,6,8-I', '1,4-Q', 'PTX', 
                                            '4,6-Q', 'aPTX'))) %>% 
  #count(population, family, saporito2017_source) %>%
  pivot_wider(names_from = family, values_from = abundance, values_fill = 0) %>%
  ungroup() %>% 
  group_by(population) %>%
  group_split() %>%
  set_names(map_chr(., ~ unique(.x$population))) %>%
  map(~ .x %>%
        select(-population) %>%
        column_to_rownames("santos2016_source") %>%
        as.matrix()
      )

### Assign a color to each sector, using a named vector
colors <- c("#6AB897", "sienna3", "#3E63A6", rep("black",13))


#### Ceiba
ceiba =allpop.chord$`1`

### Check the sector names (rows and columns of the matrix)
sec.ceiba <- union(rownames(ceiba), colnames(ceiba))


### Set names
names(colors) <- sec.ceiba 

ceiba <- ceiba[c("Both", "Mites", "Ants"), c("HTX", "DHQ", "Lehm", "3,5-P", 
                                             "Pip", "5,6,8-I", "1,4-Q", "5,8-I", 
                                             "Pyr", "3,5-I", "PTX", "4,6-Q", 
                                             "aPTX")]

### Run the chord diagram
ch_cei <- chordDiagram(ceiba, grid.col = colors)

#### C. Colón
C.colon =allpop.chord$`2`

### Check the sector names (rows and columns of the matrix)
sec.colon <- union(rownames(C.colon), colnames(C.colon))

### Set names
names(colors) <- sec.colon

C.colon <- C.colon[c("Both", "Mites", "Ants"), c("HTX", "DHQ", "Lehm", "3,5-P", 
                                             "Pip", "5,6,8-I", "1,4-Q", "5,8-I", 
                                             "Pyr", "3,5-I", "PTX", "4,6-Q", 
                                             "aPTX")]

### Run the chord diagram
ch_col <- chordDiagram(C.colon, grid.col = colors)

#### P. Quito
P.quito =allpop.chord$`3`

### Check the sector names (rows and columns of the matrix)
sec.quito <- union(rownames(P.quito), colnames(P.quito))
### Set names
names(colors) <- sec.quito

P.quito <- P.quito[c("Both", "Mites", "Ants"), c("HTX", "DHQ", "Lehm", "3,5-P", 
                                             "Pip", "5,6,8-I", "1,4-Q", "5,8-I", 
                                             "Pyr", "3,5-I", "PTX", "4,6-Q", 
                                             "aPTX")]
### Run the chord diagram
ch_qui <- chordDiagram(P.quito, grid.col = colors)

#### S. Domingo
S.domingo =allpop.chord$`4`

### Check the sector names (rows and columns of the matrix)
sec.domingo <- union(rownames(S.domingo), colnames(S.domingo))
### Set names
names(colors) <- sec.domingo

S.domingo <- S.domingo[c("Both", "Mites", "Ants"), c("HTX", "DHQ", "Lehm", "3,5-P", 
                                             "Pip", "5,6,8-I", "1,4-Q", "5,8-I", 
                                             "Pyr", "3,5-I", "PTX", "4,6-Q", 
                                             "aPTX")]
### Run the chord diagram
ch_dom <- chordDiagram(S.domingo, grid.col = colors)


#### Maná
Mana =allpop.chord$`5`

### Check the sector names (rows and columns of the matrix)
sec.mana <- union(rownames(Mana), colnames(Mana))
### Set names
names(colors) <- sec.mana

Mana <- Mana[c("Both", "Mites", "Ants"), c("HTX", "DHQ", "Lehm", "3,5-P", 
                                             "Pip", "5,6,8-I", "1,4-Q", "5,8-I", 
                                             "Pyr", "3,5-I", "PTX", "4,6-Q", 
                                             "aPTX")]
### Run the chord diagram
ch_man <- chordDiagram(Mana, grid.col = colors)


#### H. infra
H.infra =allpop.chord$`6`

### Check the sector names (rows and columns of the matrix)
sec.infra <- union(rownames(H.infra), colnames(H.infra))
### Set names
names(colors) <- sec.infra

H.infra <- H.infra[c("Both", "Mites", "Ants"), c("HTX", "DHQ", "Lehm", "3,5-P", 
                                             "Pip", "5,6,8-I", "1,4-Q", "5,8-I", 
                                             "Pyr", "3,5-I", "PTX", "4,6-Q", 
                                             "aPTX")]
### Run the chord diagram
ch_inf <- chordDiagram(H.infra, grid.col = colors)

```

```{r fig3}
#| echo: false
#| fig-show: "hold"
#| fig-cap: "**Diet differs between frog species and across diablito frog populations.** **(A)** Chord diagram showing a bipartite network of the dietary interactions between two frog species, the non-toxic *H. infraguttatus* and the toxic *O. sylvatica*, and their arthropod prey in five localities. The thickness of the connecting bars represents the average number of prey items consumed, indicating its relative importance in the network. Species specificity index (ssi) is shown next to each frog species, ranging from 0 (generalist) to 1 (specialist). **(B)** Pairwise comparison of prey item consumption between frog species. Dots represent the mean and bars represent 95% confidence intervals for the estimated marginal means. Non-overlapping bars indicate statistically significant differences. **(C)** Chord diagrams showing the most abundant alkaloid families and their putative arthropod source, across frog populations (sensu Santos et al. 2016). The thickness of the connecting bars represents the summed alkaloid abundance per structural class. Histrionicotoxins (HTX); 5,8-disubstituted indolizidines (5,8-I); 3,5-disubstituted indolizidine (3,5-I); Decahydroquinoline (DHQ); 5,6,8-trisubstituted indolizidines (5,6,8-I); Pyrrolidine (Pyr); 3,5-disubstituted pyrrolizidine (3,5-P); Lehmizidine (Lehm); 1,4-disubstituted quinolizidine (1,4-Q); Piperidine (Pip); Pumiliotoxin (PTX); Allopumilliotoxin (aPTX); 4,6-disubstituted quinolizidine (4,6-Q). "
#| cap-location: margin
#| label: fig-3
#| fig-width: 9
#| fig-height: 9

grid.raster(readPNG('/Users/camilorl/Library/CloudStorage//Shared drives/LOBSU Manuscripts/O. sylvatica vs. H. infraguttatus diet (Nora)/Submission 2-JAE/Reviews/Fig3_neu.png')) 
```

[Figure 3.png](https://drive.google.com/open?id=14ED_0IzafS-Y691-62tPJ6Qijto7JwnM&usp=drive_fs){.external target="_blank"}

```{r}
#| warning: false
#| message: false
#| tbl-cap: "Summary of the results of pairwise correlations between each alkaloid class and ant & mite abundance."
#| tbl-cap-location: top
#| label: tbl-4

T4 <- diet %>% 
  filter(prey %in% c("ants", "mites")) %>% 
  pivot_wider(names_from = "prey", values_from = "count", 
              id_cols = c(frog_id, population)) %>% 
  left_join(alkadiver %>% 
              select(-latitude, -longitude, -altitude) %>% 
              mutate(total = select(., contains(".")) %>%
                       rowSums(na.rm = TRUE)), by = "frog_id") %>% 
  na.omit() %>% 
  select(ants, mites, c(indo.137: total)) %>% 
  setNames(c(names(.)[1:2], feat_tab %>% 
               select(family) %>% 
               pull(), "total")) %>%
  {split.default(., names(.))} %>%                    # group columns by name
  map_dfc(~ rowMeans(as.data.frame(.), na.rm = TRUE)) %>% 
  corr.test() %>% 
  .$ci %>%
  rownames_to_column("correlations") %>%
  filter(str_detect(correlations, "mite|ant")) %>% 
  kable(digits = 3, 
      table.attr = 'data-quarto-disable-processing="true"', "html") %>% 
  kable_classic(full_width = F, html_font = "Cambria") %>% 
  row_spec(0, bold = T)

T4

```

[Table 4](https://drive.google.com/open?id=1IIhI3pdMJwoLEyqgEI7zKh4PeCCdKZ7R&usp=drive_fs){.external target="_blank"}

## Alkaloid diversity among sites is associated with variation in leaf litter ant communities

As the diet of *O. sylvatica* is mainly composed of ants, we looked at compositional differences in leaf litter ant communities across the geographical gradient. A total of 46 ant genera were recovered from Winkler traps in leaf litter communities. The NMDS suggested a two-dimensional solution (stress = 0.130), representing partially overlapping clusters of ant communities across localities in the ordination space (@fig-4). PERMANOVA and post-hoc comparisons suggested significant differences in the composition of ant communities between all pairs of localities (PERMANOVA, F(3) = 5.0525, p \< 0.001; @tbl-5, @fig-4 A). Precipitation (r² = 0.41, p = 0.001), altitude (r² = 0.458, p = 0.001) and temperature (r² = 0.49, p = 0.001) significantly influenced ant availability across the geographical gradient (@fig-4 B). A subset of ant genus, including *Octostruma* (r² = 0.59, p = 0.001), *Pachycondyla* (r² = 0.54, p = 0.001), *Rogeria* (r² = 0.51, p = 0.001) and *Solenopsis* (r² = 0.48, p = 0.001) significantly contributed to the ordination. We next asked whether availability of leaf litter ant communities between locations is associated with skin alkaloid diversity. The procrustes analysis suggested that variation in alkaloid diversity among sites is significantly correlated with variation in availability of ant communities (r = 0.47, p-value = 0.001, 999 permutations; @fig-4 B).

```{r}
#| warning: false
#| message: false
#| output: false

### To make the plot in ggplot, we extract the dimension scores from the NMDS and add them to the main data
ants.nmds <- ant_co %>%
  mutate(NMDS1 = scores(ord.co2)$sites[,1],
         NMDS2 = scores(ord.co2)$sites[,2]) %>% 
  data.frame()

### to extract the coordinates for the vectors of the environmental variables
en_coord.ant <-  as.data.frame(scores(en.co, "vectors"))

en_coord.ant2 <- as.data.frame(scores(en.co.ants, "vectors"))

#and the plot
fig4a <- ggplot(ants.nmds, aes(x = NMDS1, y = NMDS2*-1)) + 
  #stat_ellipse(aes(colour = SITE, fill = SITE), 
  #             geom = "polygon", level = 0.95, alpha = 0.1, type = "t") +
  geom_point(aes(fill = SITE), 
             pch = 24, col = "black", size = 2, alpha = 0.7) + 
  scale_colour_manual(values = c("#a9d8c8", "#e8c95d", "#b9b09f", "#433447")) + 
  scale_fill_manual(values = c("#a9d8c8", "#e8c95d", "#b9b09f", "#433447")) + 
  geom_segment(aes(x = 0, y = 0, xend = NMDS1, yend = NMDS2*-1), 
               data = en_coord.ant, linewidth = 1, alpha = 0.5, colour = "grey30",
               arrow = arrow(type = "open", length = unit(0.1, "inches"))) + 
  geom_text(data = en_coord.ant, aes(x = NMDS1-0.1, y = NMDS2*-1), colour = "grey30", 
            fontface = "bold", label = row.names(en_coord.ant)) + 
  geom_segment(aes(x = 0, y = 0, xend = NMDS1, yend = NMDS2*-1), 
               data = en_coord.ant2[c(13,7,2,5),], linewidth = 1, 
               alpha = 0.5, colour = "red",
               arrow = arrow(type = "open", length = unit(0.1, "inches"))) + 
  geom_text(data = en_coord.ant2[c(13,7,2,5),], aes(x = NMDS1-0.1, y = NMDS2*-1), 
            colour = "red", 
            fontface = "bold", label = row.names(en_coord.ant2[c(13,7,2,5),])) +
  theme_bw() +
  theme(panel.grid.major = element_blank(), panel.grid.minor = element_blank(),
        axis.text.x = NULL, axis.text.y = NULL, legend.position = "none")

###-----------------------###
#####  Procrustes analysis  ###
###-----------------------###

#NMDS of alkaloids excluding ceiba and infragutatus
ord.alk <- metaMDS(alkadiver %>% 
                      filter(!(population=="ceiba")) %>%
                      filter(!(population=="H_infra")) %>%
                      select(where(is.numeric), -altitude,
                             -latitude,-longitude), 
  perm=9999, distance = "bray", 
               k = 2, autotransform = FALSE)

#the ordination was ran a second time starting at the previous best solution to ensure the stability and reliability of the results in the NMDS 
ord.alk2<-metaMDS(alkadiver %>% 
                      filter(!(population=="ceiba")) %>% 
                      filter(!(population=="H_infra")) %>%
                      select(where(is.numeric), -altitude,
                             -latitude,-longitude), 
                  previous.best = TRUE)

#### NMDS of ants removing an outlier
set.seed(123)

ant_co2 <- ant_co %>% #remove one sample from P. quito that is an outlier in the ordination (only 1 wasmannia)
  group_by(SITE) %>%
  filter(!(SITE=="Puerto Quito")) %>% 
  slice_sample(n = 10) %>% 
  bind_rows(ant_co %>% 
              filter(SITE=="Puerto Quito")) %>% 
  mutate(SITE = factor(SITE)) %>% 
  ungroup() %>% 
  mutate(SITE = fct_relevel(SITE, "Mana", "Colon", "Santo Domingo", 
                            "Puerto Quito")) %>%
  arrange(as.integer(SITE))

#the ordination was performed using Bray-Curtis dissimilarities
ord.co3 <- metaMDS(ant_co2 %>%
                     select(where(is.numeric), -altitude, 
                            -latitude, -longitude), 
                  perm=9999, distance = "bray",
                  k = 2, autotransform = FALSE)

proc <- procrustes(ord.co3,ord.alk2, scale = T)


procru.test <- protest(ord.co3,ord.alk2, permutations = 999)


Yrot <- data.frame(proc$Yrot) %>% 
  bind_cols(alkadiver %>% 
                      filter(!(population=="ceiba")) %>% 
                      filter(!(population=="H_infra")) %>%
              select(population)) %>% 
  bind_cols(data.frame(proc$X))

procu.plot <- ggplot(Yrot, aes(x = NMDS1, y = NMDS2, colour = population)) +
  geom_hline(yintercept = 0, linetype = "dashed", color = "gray", alpha = 0.5) +
  geom_vline(xintercept = 0, linetype = "dashed", color = "gray", alpha = 0.5) +
  geom_segment(aes(x = X1, y = X2, 
                   xend = NMDS1, yend = NMDS2,
                   colour = population), 
               alpha = 0.2) + 
  geom_point(aes(fill = population), pch = 24, size = 1.3) +
  geom_point(aes(X1, X2, fill = population), 
             pch = 21, col = "black", size = 3, alpha = 0.7) +
  theme_bw() +
  theme(panel.grid.major = element_blank(), panel.grid.minor = element_blank(),
        axis.text.x = NULL, axis.text.y = NULL, legend.position = "none") +
  scale_fill_manual(values=c("#e8c95d", "#a9d8c8", "#b9b09f", "#433447")) +
  scale_color_manual(values=c("#e8c95d", "#a9d8c8", "#b9b09f", "#433447"))

fig4c <- ant_all %>%
  filter(TRAP == "Winkler") %>%
  select(SITE, GENERA, X.) %>% 
  group_by(SITE, GENERA) %>% 
  mutate(total = sum(X.)) %>% 
  select(-X.) %>% 
  distinct() %>% 
  filter(total>10) %>% 
  mutate(GENERA = factor(GENERA, levels = c("Solenopsis", "Wasmannia", "Pheidole", 
                                        "Nylanderia", "Carebara", "Rogeria", "Strumigenys", "Hypoponera",
                                        "Octostruma", "Apterostigma", "Cyphomyrmex", "Prionopelta",
                                        "Stenamma", "Anochetus"))) %>% 
  mutate(SITE = factor(SITE, levels = c("Mana", "Santo Domingo","Puerto Quito", 
                                        "Colon"))) %>% 
  ggplot(aes(x = GENERA, y = SITE, colour = total)) + 
  geom_point(aes(size = total, colour = total)) +
  scale_color_gradientn(colours = pal) +
  theme_bw() + 
  theme(axis.text.x = element_text(angle = 45, hjust = 1, vjust = 1),
        legend.text.position = "right")
```

```{r fig4}
#| echo: false
#| fig-show: "hold"
#| fig-cap: "**Ant community composition varies across localities.** **(A)** Non-metric multidimensional scaling (NMDS) biplot based on Bray-Curtis dissimilarities showing ant communities across locations based on the relative abundances of 113 ant species (Stress = 0.156). Each triangle represents ant communities at one sampling point. Environmental factors (black arrows) and ant genera (blue arrows) in correlation with the ordination are shown. **(B)** Procrustes ordination plot showing correspondence between frog alkaloids and ant communities ordinations across localities. Procrustes residuals (lines) connect paired points between frog alkaloid profiles (circles) and ant community composition (triangles) at each site. Shorter lines indicate better alignment between the ordinations, suggesting stronger correspondence. **(C)** Bubble - heatmap shows the number of ants, grouped by genus (with ≥10 individuals), across frog populations.  Each ant community was sampled using a trap placed next to the corresponding frog’s capture site."
#| cap-location: margin
#| label: fig-4
#| fig-width: 9
#| fig-height: 7

grid.raster(readPNG('/Users/camilorl/Library/CloudStorage//Shared drives/LOBSU Manuscripts/O. sylvatica vs. H. infraguttatus diet (Nora)/Submission 2-JAE/Reviews/Fig4_neu.png')) 
```

[Figure 4.png](https://drive.google.com/open?id=16c7Gr7NC2oDNSFNwXgQKZ6Zpk6mkSrXo&usp=drive_fs){.external target="_blank"}

```{r}
#| warning: false
#| message: false
#| tbl-cap: "Summary of the results of pairwise comparisons of a PERMANOVA in leaf litter ant composition between study sites" 
#| tbl-cap-location: top
#| label: tbl-5

colnames(pair.per.ants) <- c("Contrasts", "df", "Sums of sq.", "F", 
                        "R2","p-value", "p.adjusted", "Sig.")

T5 <- pair.per.ants %>%
  select(Contrasts, df, F, R2, p.adjusted) %>% 
  mutate(Contrasts = c("La Maná vs. C. colón","La Maná vs. S. Domingo", 
                       "La Maná vs. P. Quito","C. colón vs. S. Domingo",
                       "C. Colón vs. P. Quito","S. Domingo vs. P. Quito")) %>% 
  kable(digits = 3, 
      table.attr = 'data-quarto-disable-processing="true"', "html",
      caption = "Table S5: Summary of the results of pairwise comparisons of a PERMANOVA in leaf litter ant composition between study sites") %>% 
  kable_classic(full_width = F, html_font = "Cambria") %>% 
  row_spec(0, bold = T) 

T5
```

[Table 5](https://drive.google.com/open?id=1KPDjtI3U1hgZR0g7i_-C4SH8ma9fXDmU&usp=drive_fs){.external target="_blank"}

## Frogs show different dietary selectivity for particular ant genera

We next asked if frogs eat specific ant genera selectively or if their ant diet reflects the genera of the surrounding leaf litter communities. From the 46 ant genera recovered from Winkler traps in leaf litter communities, only 17 of these were consumed by frogs across different species and populations. Our results indicated no significant differences in the total abundance of ants between frog stomach contents and leaf litter across all populations, except for Santo Domingo and *H. infraguttatus* where frogs had lower abundance of ants in their stomachs compared to the leaf litter (@tbl-6; @fig-5 A). We found that *Solenopsis* was the most selected genus across all *O. sylvatica* populations, while *Pheidole* was the only selected genus by *H. infraguttatus*. Particularly, frogs from the Cristóbal Colón population showed selectivity for *Paratrachymyrmex*, *Crematogaster* and *Pheidole*, whereas *Strumigenys* and *Cyphomyrmex* were selected in La Maná and Puerto Quito populations, respectively (@fig-5 B).

```{r}
#| warning: false
#| message: false

f5a <- sum_abun %>% 
  ggplot(aes(x = group, y = abundance, fill = population)) +
  geom_boxplot(outlier.shape = NA) + 
  theme_bw() +
  theme(panel.grid.major = element_blank(),
                                    panel.grid.minor = element_blank(),
                                    axis.text.x = NULL, legend.position = "none") +
  geom_jitter(position=position_jitterdodge(0.1), shape = 21, 
              size = 2.5, alpha=0.6) +
  scale_fill_manual(values = c("#e8c95d", "#d16b54","#ffffff", "#a9d8c8",
                               "#433447", "#b9b09f")) +
  facet_grid(.~population) +
  scale_y_sqrt() + 
  scale_y_continuous(trans = 'sqrt') +
  ylab(NULL)

#ggsave("Fig4a.svg", f4a, units = "cm", width = 18, height = 8)


f5b <- ggplot() + 
  geom_point(data = Lsel_sim, aes(x = value, y = Genus), col = "gray", 
             pch = 15, size = 4) +
  geom_point(data = Lsel_obs, aes(x = Linear, y = Genus, fill = Cat.Linear), size = 2.5, pch = 21) +
  scale_fill_manual(values=c("black", "white", "blue")) +
  facet_grid(.~population, scales = "free")  +
  theme_bw() + 
  theme(panel.grid.major = element_blank(),
        panel.grid.minor = element_blank(),
        axis.text.x = NULL, 
        legend.position = "none") +
  xlab("Linear Selectivity") +
  ylab("Ant species")

#ggsave("Fig4b.svg", f4b, units = "cm", width = 20, height = 7)

```

```{r fig5}
#| echo: false
#| fig-show: "hold"
#| fig-cap: "**Relative abundance and selectivity for ant genera differs across localities.  **(A)** Boxplot showing differences across populations in total abundance of ants within 17 ant genera found in both leaf litter and frog stomach samples. The y axis is square-root transformed for visual clarity. n.s. = non-significant. * p-value<0.05 **(B)** Linear selectivity index for 17 ant genera eaten by the toxic *O. sylvatica* populations and the non-toxic *H. infraguttatus*. Grey bars denote simulated null distribution. Points denote categorical selectivity as follows: ‘non-selected’ if they are below the null distribution (red dots), ‘neutral’ if they are within (black dots), and ‘selected’ if the values are above (blue dots). Blue arrows indicate overall selected ant genera."
#| cap-location: margin
#| label: fig-5
#| fig-width: 9
#| fig-height: 7

grid.raster(readPNG('/Users/camilorl/Library/CloudStorage//Shared drives/LOBSU Manuscripts/O. sylvatica vs. H. infraguttatus diet (Nora)/Submission 2-JAE/Reviews/Fig5_neu.png')) 
```

[Figure 5.png](https://drive.google.com/open?id=11sziDPGX2wcmDQwgszWA2hlAB6Xfy6G4&usp=drive_fs){.external target="_blank"}

On the other hand, The remaining ants were either not selected or occasionally consumed (@fig-5 B). For example, *Wasmannia* ants were avoided by all populations, except for Santo Domingo frogs where it was consumed in proportion to its availability (i.e., neutral). Similarly, *Apterostigma* ants were avoided by frogs in Santo Domingo and in the sympatric populations of *O. sylvatica* and *H. infraguttatus* in La Maná.

```{r}
#| warning: false
#| message: false
#| tbl-cap: "Summary of the results of pairwise comparisons of a Negative Binomial regresion comparing ant abundance between *O. sylvatica* populations. P-values were adjusted using Tukey’s method"
#| tbl-cap-location: margin
#| label: tbl-6

t6 <- data.frame(sumabun.emmeans$`pairwise differences of group, population`)

conts <- c("frog c_colon - leaf c_colon", "frog p_quito - leaf p_quito", "frog s_domingo - leaf s_domingo", "frog la_mana - leaf la_mana", "frog H_infraguttatus - leaf H_infraguttatus")

T6 <- t6 %>%
  select(X1, estimate, SE, p.value) %>% 
  mutate(Contrasts = X1) %>% 
  select(Contrasts, estimate, SE, p.value) %>% 
  filter(Contrasts %in% conts) %>% 
  mutate(Contrasts = c("frog C. Colón vs. leaf C. Colón", "frog P. Quito vs. leaf P. Quito", 
                   "frog S. Domingo vs. leaf S. Domingo", "frog La Maná vs. leaf La Maná", 
                   "frog H. infraguttatus vs. leaf H. infraguttatus")) %>% 
  select(Contrasts, estimate, SE, p.value) %>% 
  kable(digits = 3, 
      table.attr = 'data-quarto-disable-processing="true"', "html",
      caption = "Table S6: Summary of the results of pairwise comparisons of a Negative Binomial regresion comparing ant abundance between *O. sylvatica* populations. P-values were adjusted using Tukey’s method") %>% 
  kable_classic(full_width = F, html_font = "Cambria") %>% 
  row_spec(0, bold = T)

T6

```

[Table 6](https://drive.google.com/open?id=1AfF9pDct9Lp07jet2-isSY0DEzimVNhU&usp=drive_fs){.external target="_blank"}

## Ant morphology influences prey selectivity in frogs

Finally, we asked whether *O. sylvatica* frogs select for specific ant traits. A principal components analysis on 17 morphological and life history ant traits yielded three principal components, each with eigenvalues exceeding 1. After inverting the scores of these components to aid interpretability, we found that the first component accounted for 55.66% of the total variance, primarily reflecting size and texture traits, with higher scores indicating larger ants. Component number two explained 11.02% of the variance and was strongly associated with color traits, where higher scores corresponded to ants with dark-brown body coloration. The third principal component, explaining 7.15% of the variance, was largely indicative of the presence of spines, with higher scores representing spiny ants (@fig-6). A pairwise Wilcoxon test on these principal components revealed that ants categorized as “selected” were significantly smaller than those with neutral selectivity (p-value = 0.03; @fig-6 A; @tbl-7), but not than those not selected (p-value = 0.162; @fig-6 A). Contrary, no differences were found in the traits of color and the presence of spines across the categories of selectivity (all p values \>0.05; @fig-6 B & C; @tbl-7).

```{r}
#| warning: false
#| message: false

### Plot contributions of each variable to each component 
fig6a <- PCAtraits$co %>% 
  data.frame() %>% 
  rownames_to_column(var = "Trait") %>% 
  select(Trait, Comp1, Comp2, Comp3) %>%
  melt() %>% 
  arrange(desc(value)) %>% 
  arrange(desc(variable)) %>% 
  filter(variable %in% c("Comp1", "Comp2", "Comp3")) %>% 
  mutate(Trait = fct_relevel(Trait, "WebersL", "BodyL", "HeadL", "HindFemurL", "PronotumW",
                                "HeadW", "InterOcularW", "MandibleL", "EyeL", "ScapeL", "ClypeusL",
                                "Pilosity", "Sculpturing", "nSpines", "ColourGaster", "Colour.Mesosoma",
                                "ColourHead")) %>% 
  mutate(value = value*-1) %>% 
  mutate(sign = ifelse(value >= 0, "Positive", "Negative")) %>% 
  ggplot(aes(x = Trait, y = value, fill = interaction(variable, sign))) + 
  geom_bar (stat="identity",position = position_dodge(0.9), col = "black", lwd = 0.2) +
  scale_y_continuous(limits = c(-1,1)) +
  facet_grid(.~variable) + coord_flip() +
  geom_hline(yintercept = 0, linetype = 3, col = "grey") +
  scale_fill_manual(values = c("Comp1.Positive" = "#a9d8c8", "Comp2.Positive" = "orange", 
                               "Comp3.Positive" = "beige", "Comp1.Negative" = "gray", 
                               "Comp2.Negative" = "gray", "Comp3.Negative" = "gray")) + 
  theme_bw() + theme(panel.grid.major = element_blank(),
                     panel.grid.minor = element_blank(),
                     axis.text.x = NULL, legend.position = "none") 

#Generate data with the mean of first three components for every Genus and site, and linear selectivity (continuous and categorical)
elect.morph <- na.omit(ant_traits) %>% 
  filter(Genus %in% Lsel_obs$Genus) %>% 
  bind_cols(PCAtraits$li) %>% # merge with principal components
  select(Genus, HeadW:ColourGaster, Axis1, Axis2, Axis3) %>% #select only the first three components
  mutate(Axis1 = Axis1*-1, Axis2 = Axis2*-1, Axis3 = Axis3*-1) %>% #PCs are multiplied by -1 for better interpretation 
  rename(PC1 =Axis1, PC2 = Axis2, PC3 = Axis3) %>% 
  group_by(Genus) %>% 
  summarise(mPC1 = mean(PC1), mPC2 = mean(PC2), mPC3 = mean(PC3)) %>%
  left_join(Lsel_obs) %>% #merge with linear selectivity data
  group_by(population) %>% 
  arrange(desc(Linear)) %>%
  mutate(Genus = reorder(Genus, Linear)) %>%
  ungroup() %>% 
  mutate(Cat.Linear = factor(Cat.Linear, levels = c("preferred", "neutral", "avoid")))

### Plot principal components against electivity categories
fig6b <- elect.morph %>% 
  select(-variable) %>% 
  melt(variable.name = "Component") %>% 
  filter(Component %in% c("mPC1", "mPC2", "mPC3")) %>% 
  ggplot(aes(Cat.Linear, value, fill = Cat.Linear)) + 
  geom_boxplot(outlier.shape = NA) + 
  scale_fill_manual(values = c("#dfb92aff", "#88b000ff", "#3f98c8ff")) +
  geom_jitter(position=position_jitterdodge(0.5), shape = 21, 
              size = 4, alpha=0.6) + 
  facet_wrap(.~Component, scales = "free_y") +
  theme_bw() + theme(panel.grid.major = element_blank(),
                     panel.grid.minor = element_blank(),
                     axis.text.x = NULL, legend.position = "none")
```

```{r fig6}
#| echo: false
#| fig-show: "hold"
#| fig-cap: "**Boxplots showing differences between categorical selectivity in principal component scores on 17 ant traits. **(A)** First principal component (PC1) representing size and texture traits. High score values depict larger ants and low score values depict smaller ants. **(B)** Second principal component (PC2) representing body coloration. High score values depict dark-brown ants, middle scores depict light-brown ants, and low score values depict black ants. **(C)** Third principal component (PC3) representing number of spines. High score values depict _spiny_ ants, whule low score values depict _non-spiny_ ants. Groups not connected by the same letter are significantly different."
#| cap-location: margin
#| label: fig-6
#| fig-width: 10

grid.raster(readPNG('/Users/camilorl/Library/CloudStorage//Shared drives/LOBSU Manuscripts/O. sylvatica vs. H. infraguttatus diet (Nora)/Submission 2-JAE/Reviews/Fig6_neu.png')) 
```

[Figure 6.png](https://drive.google.com/open?id=1NVyqYOm7dRriJJjuLjcLpJaAfwLackCe&usp=drive_fs){.external target="_blank"}

```{r}
#| warning: false
#| message: false
#| tbl-cap: "P-values of pairwise Wilcoxon test on differences in principal components between linear selectivity categories"
#| tbl-cap-location: margin
#| label: tbl-7

### Pairwise comparisons between linear selectivity categories in the principal components

pwt1 <- pairwise.wilcox.test(elect.morph$mPC1, elect.morph$Cat.Linear, p.adjust.method = "BH")

pwt2 <- pairwise.wilcox.test(elect.morph$mPC2, elect.morph$Cat.Linear, p.adjust.method = "BH")

pwt3 <- pairwise.wilcox.test(elect.morph$mPC3, elect.morph$Cat.Linear, p.adjust.method = "BH")

T7 <- cbind(pwt1$p.value, pwt2$p.value, pwt3$p.value) %>%
  set_colnames(sub("preferred", "selected", colnames(.))) %>% 
  kable(digits = 3, 
      table.attr = 'data-quarto-disable-processing="true"', "html",
      caption = "Table S7: P-values of pairwise Wilcoxon test on differences in principal components between linear selectivity categories") %>% 
  kable_classic(full_width = F, html_font = "Cambria") %>% 
  row_spec(0, bold = T) %>% 
  add_header_above(c("", "PC1 (size & texture)" = 2, 
                     "PC2 (Color)" = 2,
                     "PC3 (Spines)" = 2))

T7
```

[Table 7](https://drive.google.com/open?id=1AgB7Iy1LzU7pMCsjxp_lNTg1VneUEBm-&usp=drive_fs){.external target="_blank"}

# Discussion

We found that *O. sylvatica* alkaloid profiles varied between populations, corresponding with changes in the availability of leaf litter ants along a geographical gradient of temperature, precipitation, and altitude. Our results align with previous studies [@myers1976; @saporito_geographic_2006; @stuckert2014; @mcgugan2016; @prates2019; @moskowitz_land_2020], and provide further evidence of the importance of environmental availability of alkaloid containing prey in shaping the chemical repertoire in poison frogs. Overall, diablito populations with higher alkaloid loads were found at cooler, high-elevation sites, where leaf litter ant community composition was more diverse, except in La Maná, where both frog alkaloids and leaf litter ants were low despite the high altitude. As altitude and temperature vary along geographical gradients, they can drive changes in frog alkaloid profiles indirectly by shaping the composition and diversity of their arthropod prey [@mackay1986; @brühl1999; @silva2014; @wise2019; @moses2021]. This is consistent with our previous work where we showed that alkaloid profiles, diet, and surrounding leaf litter communities in the diablito population from Santo Domingo were more abundant in frogs from a cooler, humid forest than in a hot, dry pasture [@moskowitz_land_2020]. Other factors like chemical diversity of arthropod prey have been shown to influence alkaloid variability in diablito frogs [@mcgugan2016]. Future work comparing environmental arthropod chemistry is necessary to better understand the interplay between chemical repertoire and environmental availability of prey in organisms with diet-acquired defenses.

We found that *H. infraguttatus*, which is typically considered chemically ‘undefended’, has lower yet detectable amounts of alkaloids in the skin, compared to the sympatric chemically defended *O. sylvatica*. Our results align with the hypothesis that ‘undefended’ dendrobatids can accumulate alkaloids even in small amounts, and that the ability to acquire alkaloids for defense may be more common across dendrobatids [@tarvin_passive_2024]. In this study histrionicotoxins and indolizidines made up the highest proportion of the alkaloid profiles in both *O. sylvatica* populations and *H. infraguttatus*, which correspond with what has been found in other dendrobatids, including *Oophaga* and *Hyloxalus* frogs [@saporito_geographic_2006; @mcgugan2016; @jeckel2019; @moskowitz_land_2020; @tarvin_passive_2024]. Few studies have shown the variability of individual alkaloids and their influence on frog fitness. The potency or organismal effect of most poison frog alkaloids is unknown and there are debates about which assays are most appropriate to test potency [@weldon_poison_2017]. Some poison frog alkaloids are toxic, while other alkaloids are more noxious [@daly2005; @santos2016; @nayik_handbook_2022]. Birds avoided eating food associated with skin secretions from one *Dendrobates tinctorius* population and this difference was partly explained by 15 alkaloids, many of them indolizidines [@lawrence_linking_2023]. Given the variation in *O. sylvatica* alkaloid profiles, it is likely that there are differences in palatability to predators, although more studies on alkaloid ingestion with natural predators are needed to understand this relationship.

Organisms with diet-derived defenses must maintain their defensive reservoirs through balancing food choice and availability. In our study, sympatric groups of defended *O. sylvatica* ate more ants and mites than *H. infraguttatus*, which ate a less specialized diet of ants, beetles and arthropods gathered in the ‘other’ category, demonstrating dietary differences between species with similar prey access. Additionally, although *O. sylvatic*a differed overall in diet and alkaloid profiles across populations, the majority of their stomach contents were constituted by ants, consistent with previous studies [@mcgugan2016; @moskowitz_land_2020]. Our results are also consistent with a broader dietary study of dendrobatids in which chemically defended species, such as *Epipedobates anthonyi*, *Dendrobates auratus*, *O. pumilio*, showed a greater degree of dietary specialization on ants than ‘undefended’ species, such as *Allobates femoralis*, *Allobates zaparo*, *Hyloxalus infraguttatus* [@caldwell_evolution_1996; @darst2005]. However, some cryptically colored dendrobatid species eat mainly ants and some chemically defended species have a broader diet than expected [@toft_evolution_1995; @darst2005; @tarvin_passive_2024]. Our data is consistent with growing evidence that ants make up a large portion of the diet in many *Hyloxalus* species [@darst2005; @tarvin_passive_2024; @sanchez_loja_conspicuous_2023]. Overall, our results support the broad trend of high alkaloid-bearing dendrobatids being ant and mite specialists, while providing further evidence that poison frogs with low alkaloid levels also consume important amounts of ants.

Consuming specific arthropod prey at rates disproportionate to their availability can influence poison frogs’ alkaloid profile, as certain alkaloid classes have known origins in specific arthropod taxa [@blum_alkaloidal_1980; @mcgugan2016; @santos2016; @saporito_oribatid_2007; @saporito_formicine_2004; @spande_occurrence_1999]. In all *O. sylvatica* populations, frogs consistently selected *Solenopsis* ants, while *Strumigenys*, *Paratrachymyrmex*, *Crematogaster*, *Pheidole* and *Cyphomyrmex* ants were selectively consumed in specific populations, a pattern consistent with our previous findings [@mcgugan2016; @moskowitz_land_2020]. These ant genus are known sources of several alkaloid classes particularly abundant across diablito populations, including histrionicotoxins, decahydroquinolines, 3,5-disubstituted indolizidines, and pyrrolidines [@blum_alkaloidal_1980; @jones_ant_1982; @jones_further_1999; @spande_occurrence_1999; @mcgugan2016; @moskowitz_land_2020]. Dietary selectivity in *O. sylvatica* may suggest a preference for specific ant prey based on their alkaloid content, or it may simply reflect that these frogs inhabit microhabitats where alkaloid-rich ants are abundant, leading to incidental consumption without active behavioral preference, as previously observed in *O. pumilio* [@donnelly_feeding_1991]. Further behavioral assays are required to distinguish between these competing hypotheses. Additionally, given the varied, but overall high number of mites recovered from stomach contents across localities, our data suggest that mites are also an important defensive alkaloid source for *O. sylvatica*, probably of 5,8-disubstituted & 5,6,8-trisubstituted indolizidines and pumiliotoxins, as previously found in *O. pumilio* [@saporito_geographic_2006; @saporito_oribatid_2007]. Mite taxonomy and chemistry is drastically understudied compared to ants and future studies should also make efforts to include mites in their analyses [but see @saporito_oribatid_2007; @saporito_taxonomic_2015; @saporito_alkaloids_2011].

We found that *H. infraguttatus* selected *Pheidole* ants, which were also selected by *O. sylvatica* from the ‘Cristobal Colón’ locality within this study. Previous alkaloid sampling of *O. sylvatica* and *Pheidole* ants found evidence of overlapping alkaloids within the same sampling location in both frogs and ants [@moskowitz_land_2020]. Our data suggests that selection for alkaloid-rich ants is not exclusive to aposematic species. Recent evidence found that *Pheidole* ants constituted the largest portion of the diet in the cryptic *Allobates femoralis* frog, although they were not actively selected [@sanches_what_2023]. It is unclear whether *H. infraguttatus* frogs have less alkaloids because their diet is less ant and mite rich or if they lack the physiological mechanisms required for alkaloid sequestration in higher concentrations, as has been noted recently in other cryptic species [@alvarez-buylla2023]. For example, in laboratory toxin feeding trials *A. femoralis* was able to uptake alkaloids as effectively as *O. sylvatica*, but exhibited signs of physiological distress [@caty_alkaloids_2025]. Further captive feeding experiments would be helpful to fully understand the implications of dietary selectivity for the evolution of acquired chemical defenses among poison frogs.

Poison frogs may selectively consume certain ant genera based on their alkaloid content, reflecting underlying feeding preferences. However, dietary selectivity can also arise from factors beyond behavioral preference. For instance, *Solenopsis* and *Pheidole* ants may be consumed more frequently, regardless of their abundance, because they are smaller and require less handling effort than bigger, more aggressive ant genera, as observed in the chemically defended frog, *Chiasmocleis* *leucosticta* [@meurer2021]. In contrast, the Neotropical toad, *Rhinella alata*, selectively consumed larger, more conspicuous ants despite their rarity, while avoiding the more abundant but smaller *Solenopsis* ants [@mcelroy_ant_2019]. Also, as we limited our genus-level analysis to ants here, we cannot determine whether frogs select ants over other prey categories that may offer different metabolic benefits or higher nutritional value. For example, when prey availability is controlled under lab conditions, lab-raised Dendrobates tinctorius frogs gravitate to the most nutritious prey (larvae)​, regardless of size and other prey options including ants (Moskowitz et al., 2022). A careful comparison of frog diet with all available arthropods in the leaf litter through metabarcoding could be useful to determine whether behavioral selectivity is consistent across prey categories. Although we found solid evidence on the relationship between alkaloid defenses, frog diets and arthropod availability, it is worth noting that alkaloid profiles in poison frogs reflect long-term accumulation, whereas dietary composition and prey availability are measured at a single time point, potentially capturing only a subset of the frogs' long-term foraging patterns. Finally, we collected leaf litter samples with Winkler sacs, which often capture abundant small, specialist myrmicines compared to pitfall traps, which are better in capturing bigger scavengers like *Anochetus* or *Hypoponera* ants [@olson_comparison_1991; @sabu_efficacy_2010]. Integrating multiple sampling methods may provide a more ecologically realistic view of the arthropod community available to poison frogs, including variation in foraging patterns that could underestimate prey availability.

# Summary

Our study demonstrates that dietary selectivity and environmental prey availability shape poison frog chemical defenses. *Oophaga sylvatica* showed alkaloid profiles that varied with local leaf litter ant availability, while the sympatric *Hyloxalus infraguttatus*, traditionally considered ‘undefended’, carried small but detectable alkaloid levels. Both species selectively consumed certain small ant genera, suggesting toxic prey selection is not exclusive to aposematic species. These findings highlight dietary selectivity as a key factor in the evolution of chemical defenses. Future studies should replicate these findings in both natural and controlled settings, compare environmental arthropod chemistry to better understand prey availability, and use captive feeding experiments to test how diet influences alkaloid accumulation. 

\

# Acknowledgements

We thank Lola Guarderas (Wikiri) for logistical and resource support in Ecuador. We are thankful to the Peter Dorrestein lab at The University of California at San Diego for creating the GNPS environment that allows us to perform the metabolomic analysis of poison frog alkaloids. We would also like to thank Mabel Gonzalez, who was a visiting researcher in the Dorrestein Lab and helped us implement GNPS in our lab. LAC acknowledges the support of the Saint Louis Zoo. We acknowledge that our work at Stanford University takes place on the ancestral and unceded land of the Muwekma Ohlone tribe.

### Data accessibility

Diet and alkaloid data is included in supplementary materials. Gas chromatography / mass spectrometry files and arthropod photos are available on DataDryad (submission pending acceptance).

### Funding

This work was supported by the National Science Foundation (IOS-1557684 and IOS-2337580), Pew Charitable Trusts, and the New York Stem Cell Foundation to LAO. NAM and AAB are supported by National Science Foundation graduate research fellowships (DGE-1656518). AAB is supported by a Gilliam Fellowship from the Howard Hughes Medical Institute (GT13330). LAO is a New York Stem Cell Foundation – Robertson Investigator.

### Author contributions

Conceptualization: NAM, LAO. Methodology: NAM, LAO, DAD, AAB. Formal Analysis: NAM, AAB,CR. Investigation: NAM, AAB, AC, JR, DAD. Resources: LAO, DAD. Data Curation: NAM, AAB, CM, DAD, CR. Writing – original draft preparation: NAM, LAO, AAB, DAD. Writing – review and editing: NAM, LAO, AAB, DAD, ET, AC, JR, LAC, CR, KF. Visualization: NAM, LAO, CR. Supervision: LAO. Project administration: LAO. Funding acquisition: LAO.

# References
```
